# Synthesis and Evaluation of Carmofur Analogs as Antiproliferative Agents, Inhibitors to the Main Protease (M^pro^) of SARS-CoV-2, and Membrane Rupture-Inducing Agents

**DOI:** 10.1101/2024.10.11.617727

**Authors:** Tiffany Gu, Amber Lu, Xina Wang, Natalie Brahan, Lexi Xu, Leyuan Zhang, Kaitlyn Su, Kody Seow, Julia Vu, Charissa Luk, Yunseo Lee, Anirudh Raman, Joseph Pazzi, Edward Njoo

## Abstract

Initially developed as a derivatized analog of 5-fluorouracil for the treatment of colorectal cancer, carmofur has more recently demonstrated potent covalent inhibition of the main protease (M^pro^) of SARS-CoV-2. Harnessing our previously described workflow for the optimized preparation of carmofur using benchtop ^19^F NMR spectroscopy, here, we prepared and evaluated a synthetic library of nine carmofur analogs with a selection of side chain motifs or single-atom substitution to explore the diversifiability of these compounds as M^pro^ inhibitors, where we discovered that a hexyl carbamate analog outperformed carmofur, and as antiproliferative agents in model human cell lines to identify differences in potency when the carbonyl electrophilicity and/or alkyl side chains are modified. Finally, we describe a novel workflow for the evaluation of membrane-rupturing small molecules through imaging of fluorescently labeled giant unilamellar vesicles (GUVs), and through this, we identified two lipophilic urethane analogs of carmofur bearing dodecyl urethane and octadecyl urethane side chains that have potent membrane-rupturing capability in the nanomolar range, providing insight into a potential mechanism for the *in vitro* activities of lipidated 5-fluorouracil analogs.

## Introduction

The broad applicability of synthetic nucleoside analogs as small molecule antimetabolites for the treatment of cancers and infectious diseases have made them an important chemical platform for the discovery and development of modern therapeutics [1–5]. Carmofur **1** was originally developed to be a lipophilic, orally-available prodrug of the antimetabolite 5-fluorouracil (5-FU) and was approved for clinical use in Japan, China, Korea, and Finland for the treatment of colorectal cancer [6,7]. Carmofur was discovered to putatively inhibit acid ceramidase through carbamoylation of the sulfhydryl group of Cys143 on acid ceramidase, which attacks and the electrophilic carbonyl group of carmofur [8,9]. This leads the electrophile to covalently modify the active site cysteine, rendering the enzyme inactive and ultimately limiting cancer progression through ceramide-induced apoptosis [9–12].

More recently, carmofur was identified through a drug high-throughput repurposing screen to be a potent inhibitor of the SARS-CoV-2 main protease (M^pro^), a key enzyme involved in the cleavage and maturation of functional proteins responsible for viral replication [13–15]. Upon further evaluation, it was determined that the mechanism for the inhibition of M^pro^ by carmofur is also through direct covalent modification, wherein the hexyl urethane tail forms a covalent adduct with Cys145, rendering the enzyme catalytically inactive [16]. The possible function of M^pro^ as a drug target for SARS-CoV-2, alongside other carmofur analog structure-activity-relationship studies, have inspired us to design analogs that could enable more targeted and effective inhibition against acid ceramidase or M^pro^ (**Fig. 1**) [10,17,18].

**Fig. 1.**
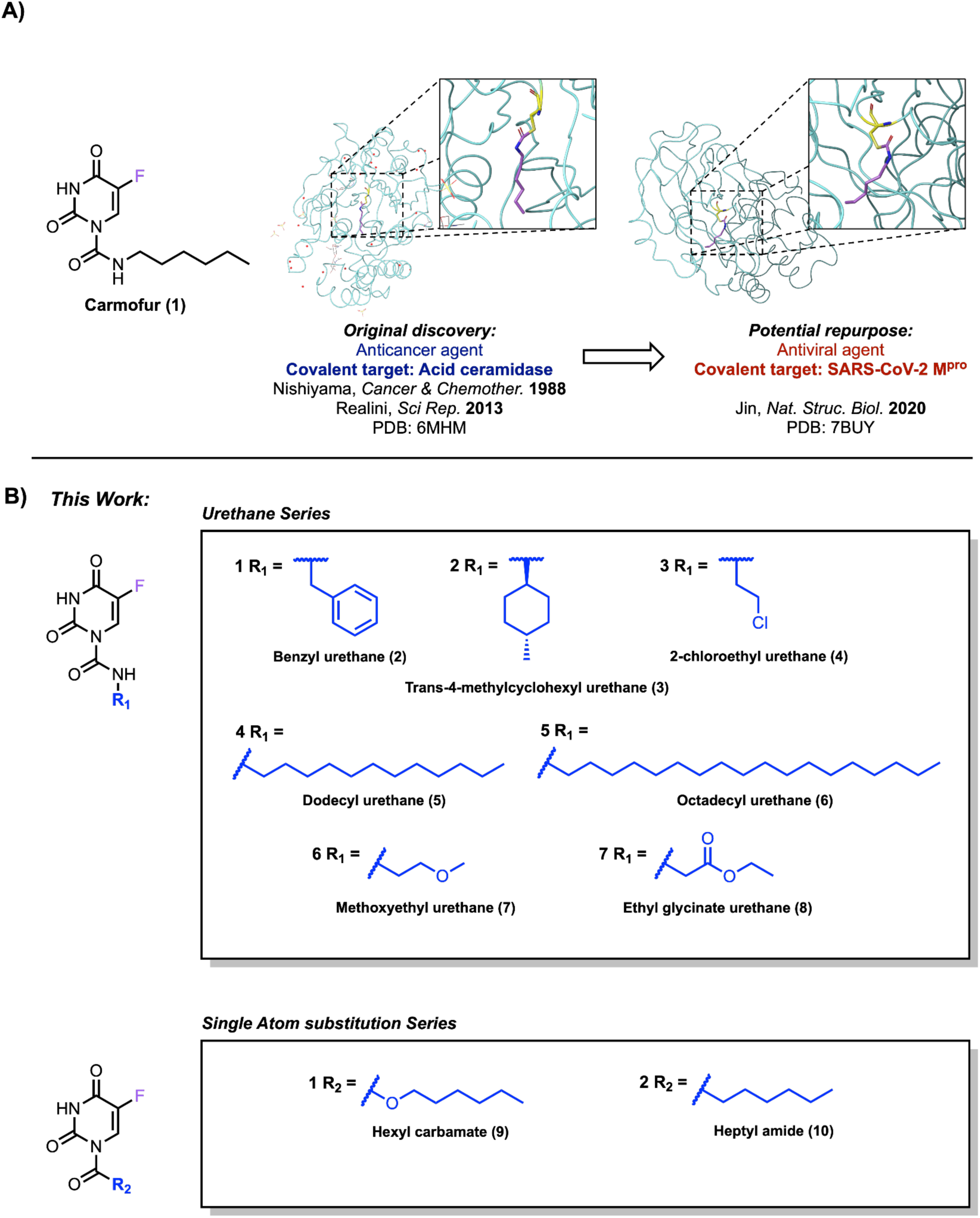
Structure of carmofur and analogs. A) Carmofur was initially discovered as an anticancer agent against colorectal cancer by Nishiyama et al. in 1988 and a covalent inhibitor of acid ceramidase by Realini et al. in 2013. In 2020, carmofur was later found to be a SARS-CoV-2 M^pro^ inhibitor by Jin et al. A magnified view of carmofur covalently bound to acid ceramidase (PDB: 6MHM) and the SARS-CoV-2 main protease (PDB: 7BUY) is shown. B) A series of urethane and single atom-substituted analogs of carmofur were prepared to probe the impact of different substitutions at R_1_ and R_2_

Previously, we reported the use of benchtop ^19^F nuclear magnetic resonance (NMR) spectroscopy as an efficient means of screening reaction conditions that have enabled the monitoring of multicomponent reactions (MCRs), preparation of trifluoromethyl pyrazole non-steroidal anti-inflammatory drugs (NSAIDs), and the optimization of the synthesis of carmofur [19–21]. Here, we generalized this workflow to the synthetic preparation of nine carmofur analogs with a selection of alkyl, aryl, and heteroalkyl side chains, as well as two single-atom substitution isosteres of carmofur to investigate the role of modifications of the urethane motif in the biological activity of our 5-FU derivatives, as a platform for the discovery of potential dual-purpose small molecules targeting acid ceramidase or SARS-CoV-2 M^pro^.

We assessed the anti-proliferative activity of our analogs in four model human colon cancer, embryonic kidney, and breast cancer cell lines as well as their inhibitory activity against SARS-CoV-2 M^pro^. We found that the benzyl urethane **2** analog exhibited moderately more potent activity in cell viability assays compared to carmofur **1** in HT-29 cells, while analogs with increased electrophilicity at the carbonyl, such as the hexyl carbamate analog **9**, exhibited improved SARS-CoV-2 M^pro^ inhibitory activity. Moreover, while most compounds of this series exhibited limited antiproliferative activity, several of the compounds bearing long, linear aliphatic urethane chains such as the dodecyl and octadecyl urethane analogs **5** and **6** were found to be potent membrane disrupting agents, and this was measured on the basis of a cell-free microscopy assay in the lysis of giant unilamellar vesicles (GUVs).

## Results and Discussion

### Chemical Synthesis

**Scheme 1.**
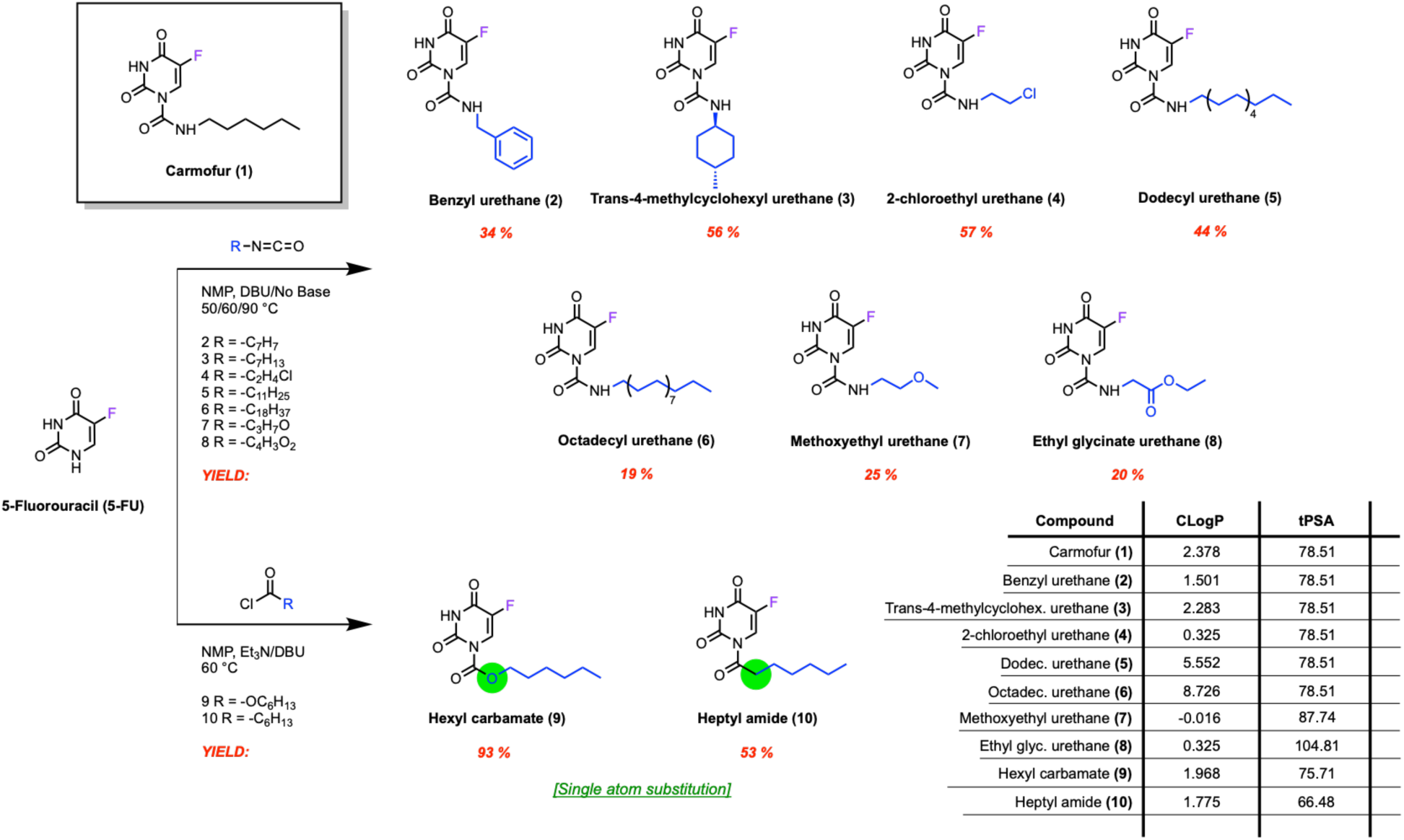
General synthetic workflow for synthesis of carmofur analogs and physicochemical properties of carmofur and its analogs. Analogs were synthesized with aryl, alkyl, or cycloalkyl substituent chains or by substituting the nitrogen atom on carmofur with an oxygen or a methylene. Isolated yields are shown above, and CLogP and tPSA values were calculated using ChemDraw

In order to interrogate the effect of side chain substitution on the biological function of carmofur and its analogs, we prepared a diverse series of analogs with various alkyl, cycloalkyl, and aryl substituents that probe the impact of sterics, lipophilicity, and electrophilicity on biological activity (Scheme 1). To probe the effects of aromaticity on pi-stack interactions within the S2 subsite of M^pro^, we synthesized the benzyl urethane analog **2** that was previously analyzed for its antitumor activity through nucleophilic addition to benzyl isocyanate [22]. Conveniently, we found that treatment of 5-FU with a variety of alkyl, benzyl, or heteroalkyl isocyanates with an amine base in N-methylpyrrolidinone (NMP) to forge the corresponding urethane was easily generalizable in the preparation of benzyl urethane **2** and other urethane analogs. We also desired to probe the impact of a cyclized chain on binding, leading us to synthesize the *trans*-4-methylcyclohexyl urethane analog **3**, with a ring of equivalent carbon length to carmofur, and which was previously reported in literature as a potential anti-tumor agent [23]. Inspired by elements of chlorinated alkylating anticancer agents such as ifosfamide and glufosfamide, we also synthesized the 2-chloroethyl urethane analog **4** [24,25].

We inferred that mimicking the lipophilic structure of ceramide, the endogenous substrate of acid ceramidase, could strengthen hydrophobic interactions with acid ceramidase. Thus, we synthesized the dodecyl and octadecyl urethane analogs **5** and **6,** which both possess a long alkyl side chain, from 5-FU with dodecyl isocyanate and octadecyl isocyanate, respectively. Next, to evaluate the potential effect of a more hydrophilic methoxyethyl ether on an otherwise hydrocarbon side chain, we synthesized a novel methoxyethyl urethane analog **7** in 25 % yield from 5-FU and 2-methoxyethyl isocyanate. In addition, we prepared the ethyl glycinate urethane analog **8** from its corresponding isocyanate to investigate if a peptidomimetic carbonyl-containing compound would better bind with the enzymatic subsite of M^pro^.

Finally, with the understanding that **1** functions through covalent inhibition of nucleophilic residues within both SARS-CoV-2 M^pro^ and acid ceramidase, we prepared hexyl carbamate **9,** where the urethane nitrogen is exchanged for an oxygen atom, and heptyl amide **10**, both of which have the same linear alkyl carbon count as **1** but where single-atom alterations might increase electrophilicity at the reactive carbonyl center [9,16]. The carbamate **9** was prepared in 93 % yield from 5-FU with hexyl chloroformate. Analogously, treatment of 5-FU with heptanoyl chloride afforded heptyl amide **10** in 53 % yield.

### Diversified analogs of carmofur exhibit different inhibition against SARS-CoV-2 M^pro^

To measure the effect our analogs have on SARS-CoV-2 activity, we sought to quantify their effect on the activity of M^pro^ with a cell-free chromogenic assay based on the release of *p-* nitroaniline from an M^pro^ recognition peptide, which is measurable at an optical readout at 410 nm [26]. We evaluated the potency of carmofur and its analogs in relative inhibition of M^pro^ activity by dosing the enzyme with 5 μM compound and measuring the absorbance values representative of the resulting M^pro^ catalytic activity after 160 minutes of incubation with the colorimetric substrate. The inhibitory activity of each compound from this series on M^pro^ activity is depicted in **Fig. 2**.

**Fig. 2.**
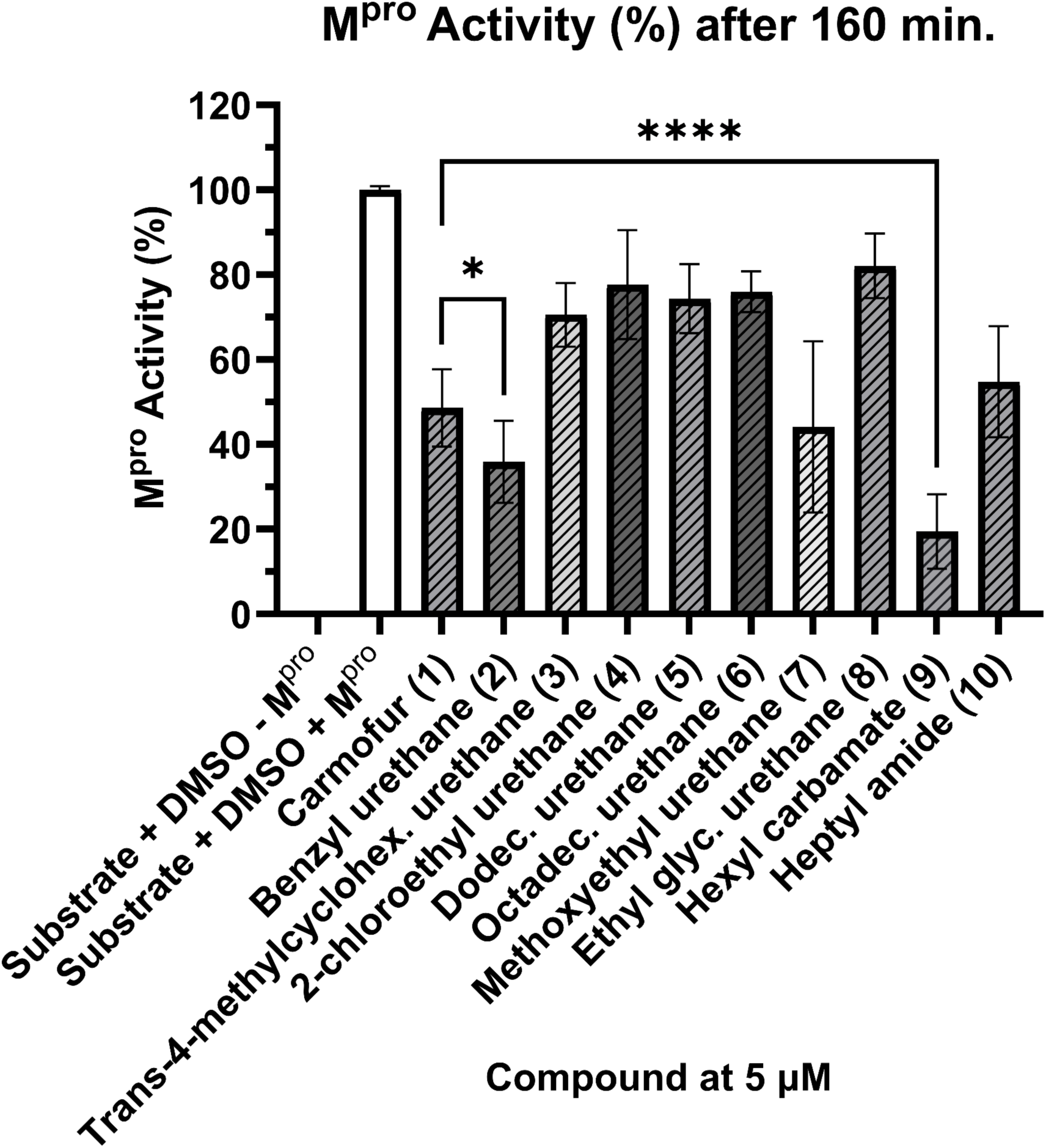
M^pro^ inhibitory activity of carmofur and its analogs. A) Percent activity of the SARS-CoV-2 M^pro^ was measured at a compound concentration of 5 µM after 160 minutes. Data is represented as means relative to the negative control (no enzyme present) ± SD compared with the positive control (1 % DMSO) (n=27) and carmofur **1** using a two-tailed unpaired student’s t test (n=9) (*P<0.05, ****P<0.0001)

After 160 minutes, all compounds significantly decreased the activity of the SARS-CoV-2 main protease compared to the positive control (P<0.0001), and we found that carmofur at a 5 µM concentration lowered M^pro^ activity to 49 ± 9.1 %, which is consistent with literature reported IC_50_ values [13,18]. The benzyl urethane and hexyl carbamate analogs **2** and **9** exhibited more potent inhibition against M^pro^ activity, decreasing it to 36 ± 10. % and 19 ± 8.8 %, respectively (**Fig. 2**). This could be due to their electronic properties; the oxygen on hexyl carbamate **9** increases the electrophilicity at the carbonyl, and the benzyl ring of **2** might make the urethane carbonyl more electrophilic than in alkyl urethane analogs.

The relatively hydrophilic methoxyethyl analog **7** with a CLogP value of −0.016 exhibited similar inhibition of M^pro^ activity compared to carmofur **1**, whereas the dodecyl urethane **5** and octadecyl urethane **6** analogs with higher CLogP values performed worse than **1**, suggesting that longer aliphatic tails are not tolerated in M^pro^ inhibition. Additionally, computer models of compounds **1** through **10** docked to the active site of SARS-CoV-2 M^pro^ (PDB: 7CAM) suggest that large, hydrophobic alkyl groups in the dodecyl **5** and octadecyl **6** urethane analogs are not easily accommodated in the main binding pocket (**Fig. 3**) [27].

**Fig. 3.**
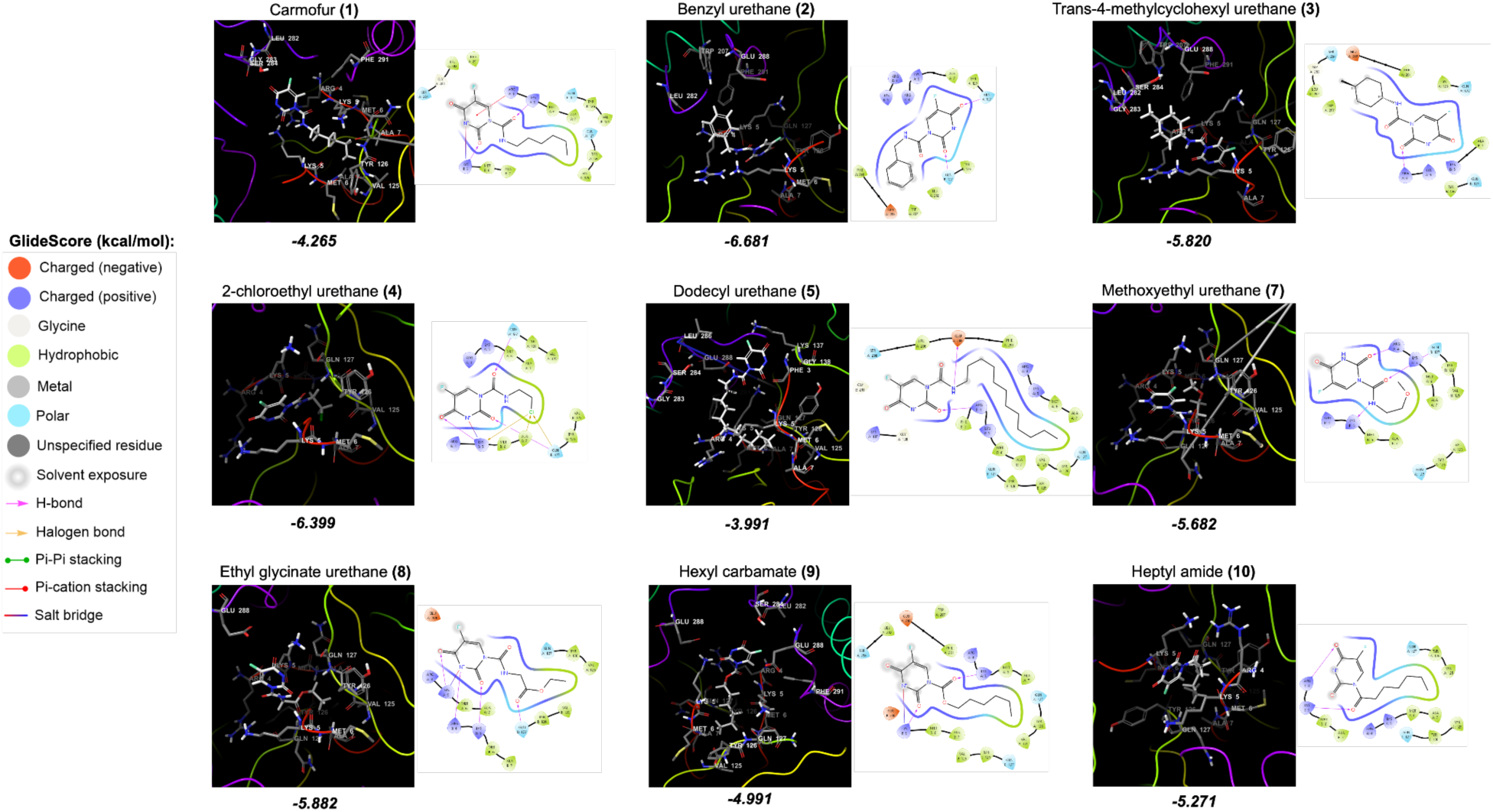
SARS-CoV-2 M^pro^ in complex with carmofur and its analogs. Docked interactions and GlideScores of the compounds against M^pro^ (PDB: 7CAM) were visualized and calculated using Schrödinger Glide. Note: The octadecyl urethane analog **6** has no conformations that bind to the M^pro^ pocket. In addition, when the docked poses were overlapped, it was observed that the 5-FU moiety had a unique conformation for each compound However, no consistent trend was observed in the inhibitory effect of the enhanced electrophilicity of the carbonyl on analogs **9** and **10**. Even though hexyl carbamate **9** performed significantly better than carmofur **1**, the heptyl amide **10**, which is more electrophilic than carmofur **1**, did not, lowering the activity of M^pro^ to 65 ± 13 % with no statistical improvement over the activity of **1**.

### Evaluating antiproliferative activity in model cell lines

To evaluate the efficacy of our analogs for *in vitro* antiproliferative activity, we measured their activity against HCT-116 and HT-29 colorectal cancer cells and MDA-MB-468 breast cancer cells. We additionally probed the potency of these compounds in HEK-293 cells, a noncancerous human embryonic kidney line (**Fig. 4**).

**Fig. 4.**
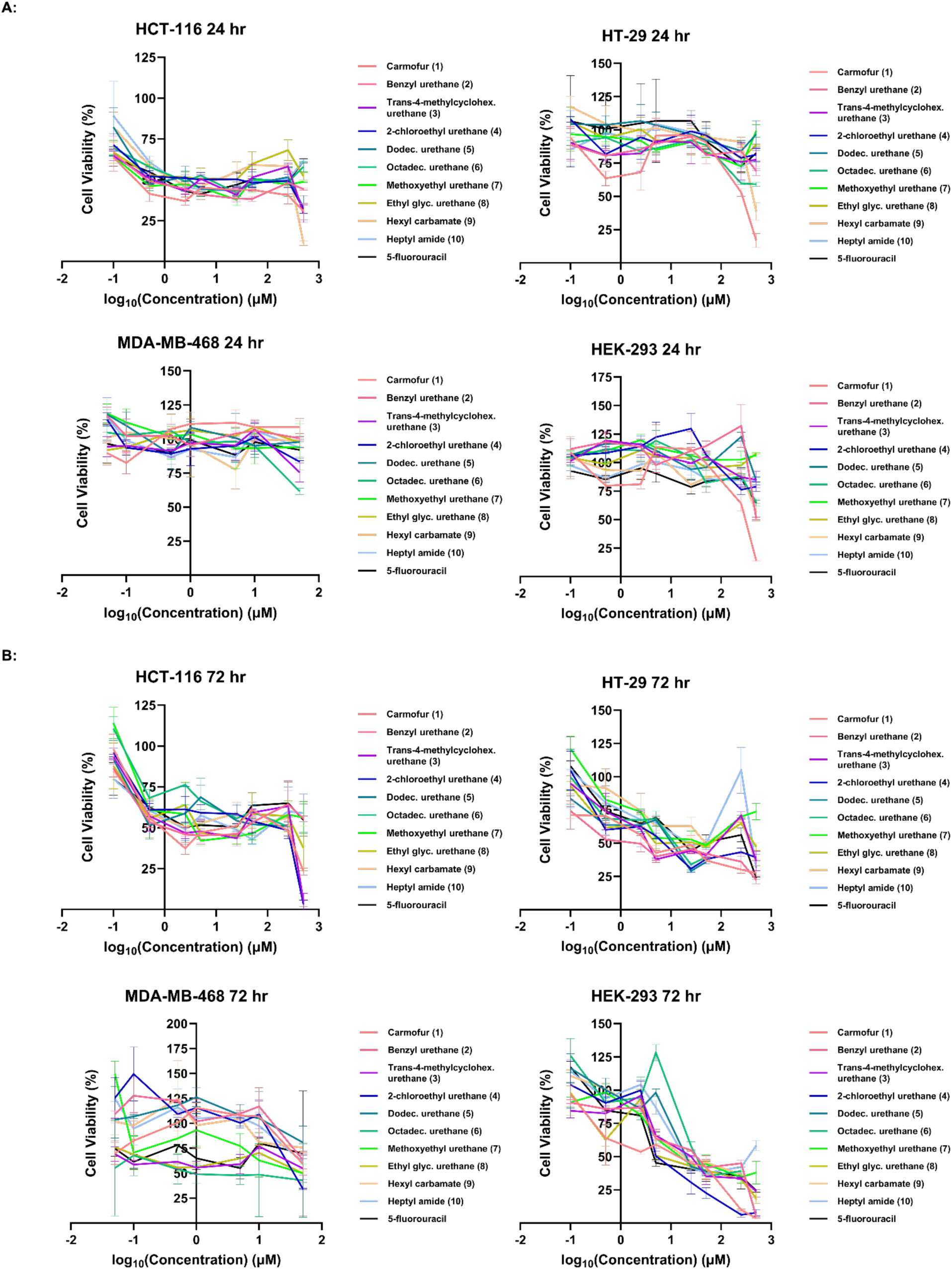
Analogs of carmofur selectively decrease cell viability of human colorectal cancer and embryonic kidney cells while showing marginal effect on non-metastatic and triple negative metastatic breast cancer cells. Cell viabilities after treatment with carmofur, 5-FU, and carmofur analogs at different concentrations (μM) after A) 24 hours and B) 72 hours on HCT-116, HT-29, MDA-MB-468, and HEK-293 cell lines. Cell viability was measured via MTT assay against a DMSO negative control. Note: Compounds **5** and **6** were poorly soluble at final concentrations of 500 µM and 250 μM and were hence excluded at those concentrations from this report. IC_50_ values could not be calculated accurately for some compounds as they exhibited limited antiproliferative activity except at the highest concentrations

**Fig. 5.**
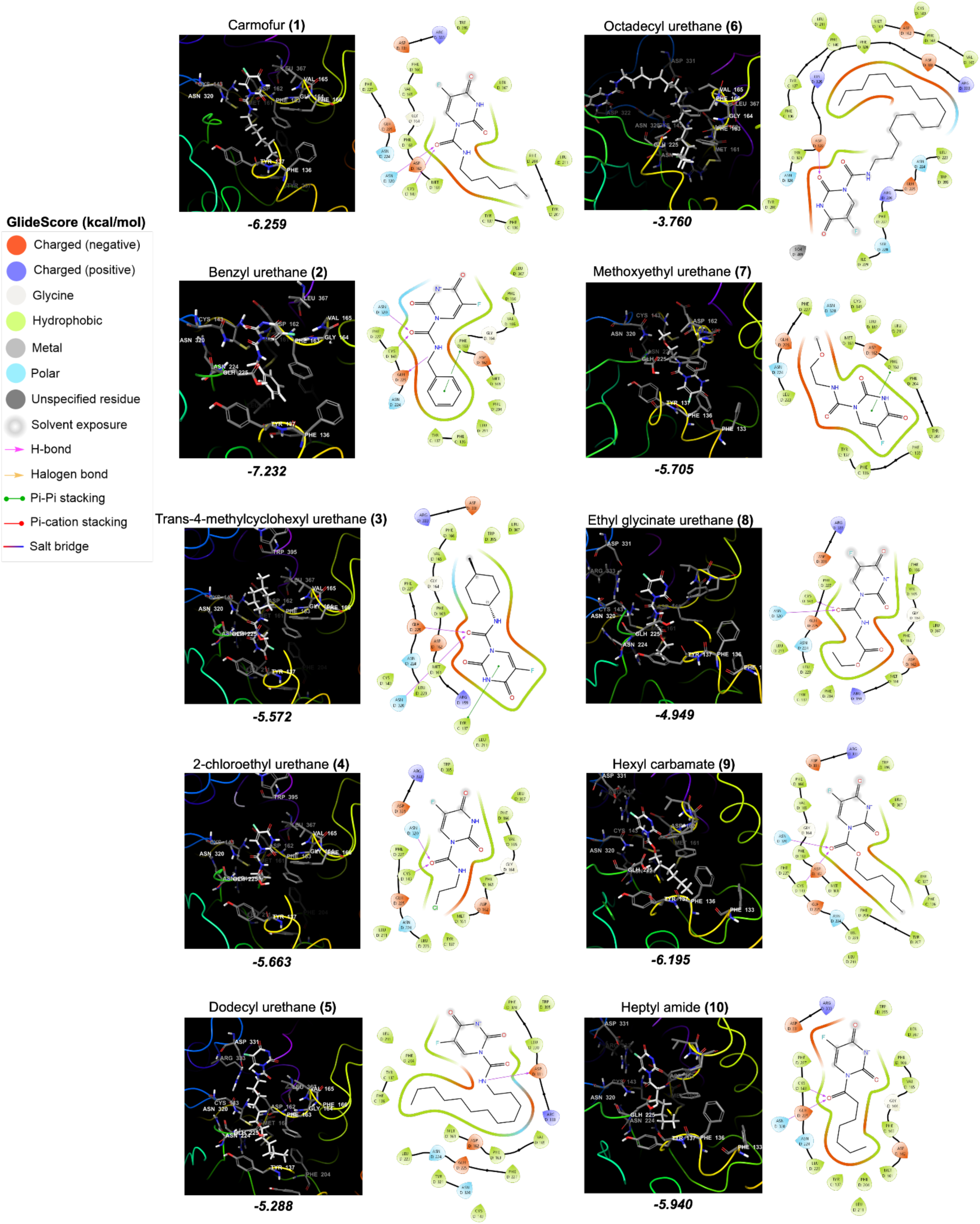
Human acid ceramidase in complex with carmofur and its analogs. Docked interactions and GlideScores of the compounds against acid ceramidase (PDB: 6MHM) were visualized and calculated using Schrödinger Glide. Note: When the docked poses were overlapped, it was observed that the 5-FU moiety had a unique conformation for each compound Docking of our compounds to the catalytic site of human acid ceramide (PDB: 6MHM) was further performed on Schrödinger Glide and suggests that initial binding of these compounds might be mediated by hydrogen bonding to Asn320 in addition to Cys143 and that longer motifs might be accommodated in the binding pocket (**Fig. 5**) [9,28–31].

At 24 hours, compounds **2**, **5**, **6**, **7**, and **9** with IC_50_ values of 6.64 μM, 5.46 μM, 6.18 μM, 9.57 μM, and 11.95 μM, respectively, expressed similar activity against HCT-116 cells at the 24 hour time point as compared to carmofur **1** (IC_50_ = 6.16 μM) (**Fig. 4A**). In **Fig. 4A**, most of our carmofur analogs were not well tolerated against the HT-29 and HEK-293 cell lines. Similarly, most compounds expressed little to no inhibition against MDA-MB-468 cells after 24 hours.

However, having an IC_50_ value of 2.91 μM, the octadecyl urethane analog **6** exhibited potent inhibition in MDA-MB-468 cells while the other compounds remained ineffective after 72 hours. The benzyl urethane analog **2** with an IC_50_ value of 3.55 μM performed slightly better than **1** with an IC_50_ value of 6.97 μM against HT-29 cells after 72 hours (**Fig. 4B**). Furthermore, analog **4** with an IC_50_ value of 11.2 μM performed comparable to **1** with an IC_50_ value of 7.15 μM against HEK-293 after 72 hours while the remaining analogs performed worse than **1** (**Fig. 4B**).

### Antiproliferative carmofur analogs cause membrane rupture in model systems

Prompted by the observation that the dodecyl urethane **5** and octadecyl urethane **6** exhibit antiproliferative activity in mammalian cell lines only at the highest soluble concentrations, we developed a novel cell-free membrane disruption assay based on quantification of the concentration-dependent membrane rupture of fluorescently labeled GUVs on a negatively charged glass surface [32]. We systematically characterized membrane rupture for each carmofur analog over concentrations ranging from 11 pM to 2.25 mM by collecting images of GUVs covering a 1 mm × 1 mm area before and after addition of the analogs. **Fig. 6** shows representative images of GUVs bursting at the surface of the glass before (**Fig. 6A**) and 15 minutes after addition of the octadecyl urethane (**Fig. 6B**). The GUVs can be observed as circular, bright red rings with dim red interiors (white arrow in **Fig. 6C**) and the patches that result following membrane rupture can be observed as unsmooth, dim splotches lacking a bright ring (white arrow in **Fig. 6D**). Consistent with the expectation that lipidated compounds may exert their *in vitro* antiproliferative activity by acting as surfactants at high concentrations, we observed consistent membrane rupture in GUVs treated with **5** and **6** (**Fig. 7**) [33–35]. For the remaining compounds, we mostly observe little to no GUV bursting even at the highest soluble concentrations with some rare but perhaps still interesting bursting occurring in heptyl amide **10** (**Supporting Fig. S2**).

**Fig. 6.**
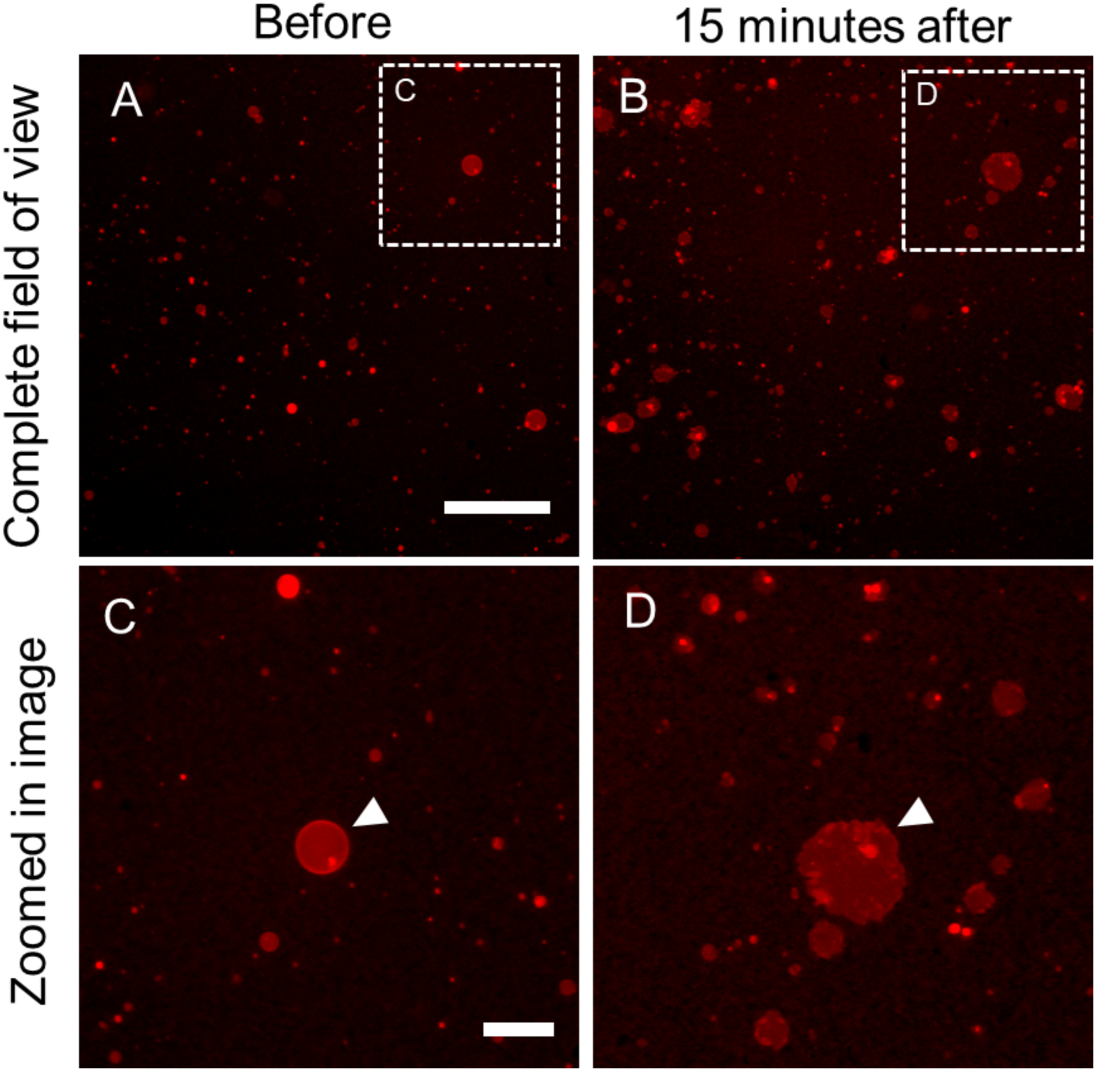
Membrane rupture assay using GUVs. **A,B** Fluorescence microscopy images of GUVs **A** before and **B** 15 minutes after addition of octadecyl urethane **6** at 112 μM. **C,D** Zoomed in images of the regions in **C** (panel **A**) and **D** (panel **B**) showing a larger GUV that burst into a patch (white arrows). Scale bars are **A,B** 200 μm, **C,D** 50 μm

**Fig. 7.**
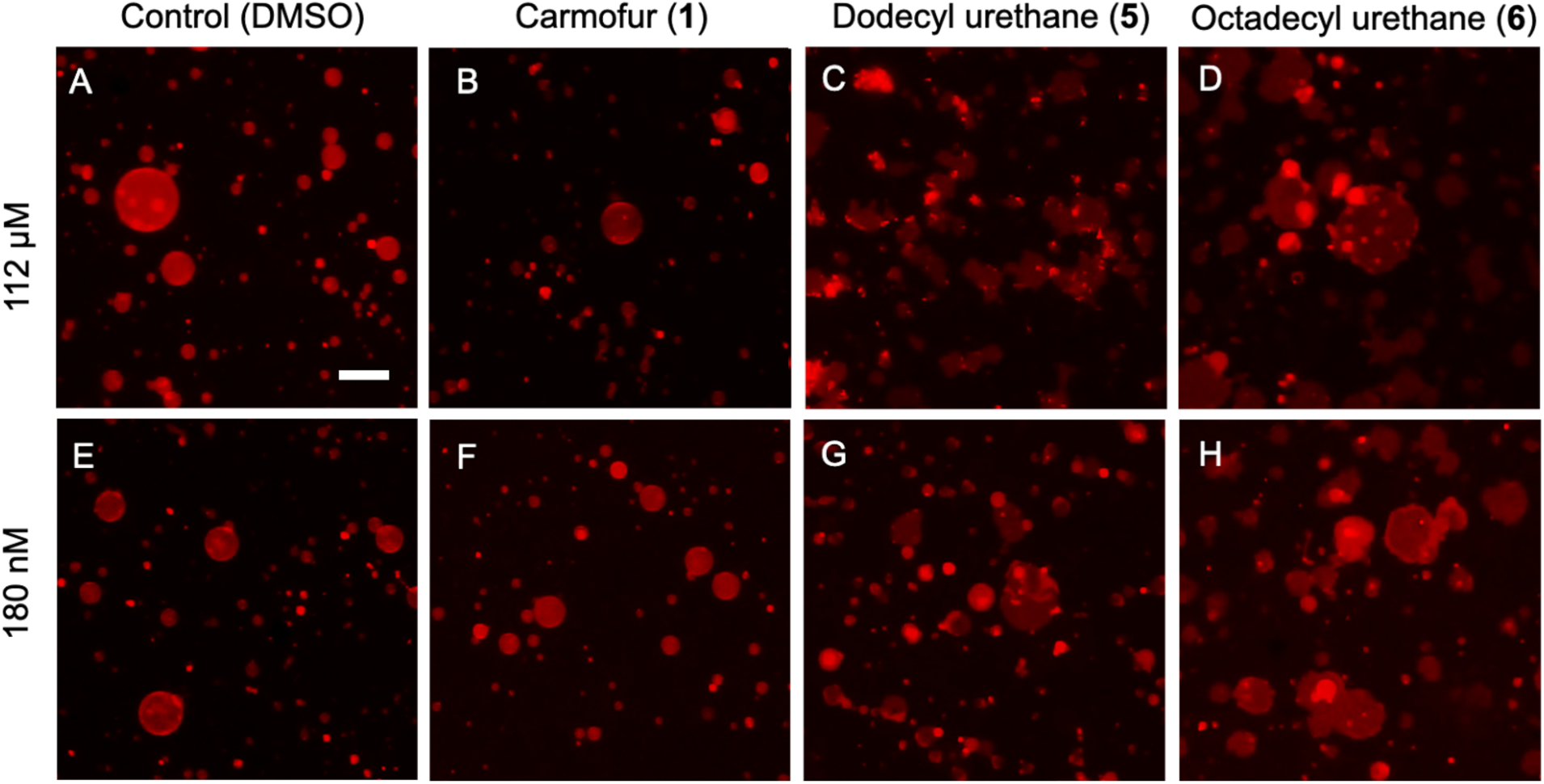
Effect of compound and concentration on GUV rupture. Representative images of GUVs 15 minutes after addition of **A** 0.0225% v/v DMSO, **B** 112 μM carmofur **1**, **C** 112 μM dodecyl urethane **5**, **D** 112 μM octadecyl urethane **6**, **E** 0.0225% v/v DMSO, **F** 180 nM carmofur **1**, **G** 180 nM dodecyl urethane **5**, and **H** 180 nM octadecyl urethane **6**. Scale bar for **A-H** is 50 μm

To further characterize membrane rupture we counted the percentage of GUVs that burst into patches from the images (**Fig. 8**). For each sample we counted approximately 500 GUVs both before and after addition of compound, and we collected data on 3 replicate samples for each compound and each concentration. The bar plots show the mean percentage of GUVs that burst into membrane patches with the error bars representing the standard deviation of the mean.

**Fig. 8.**
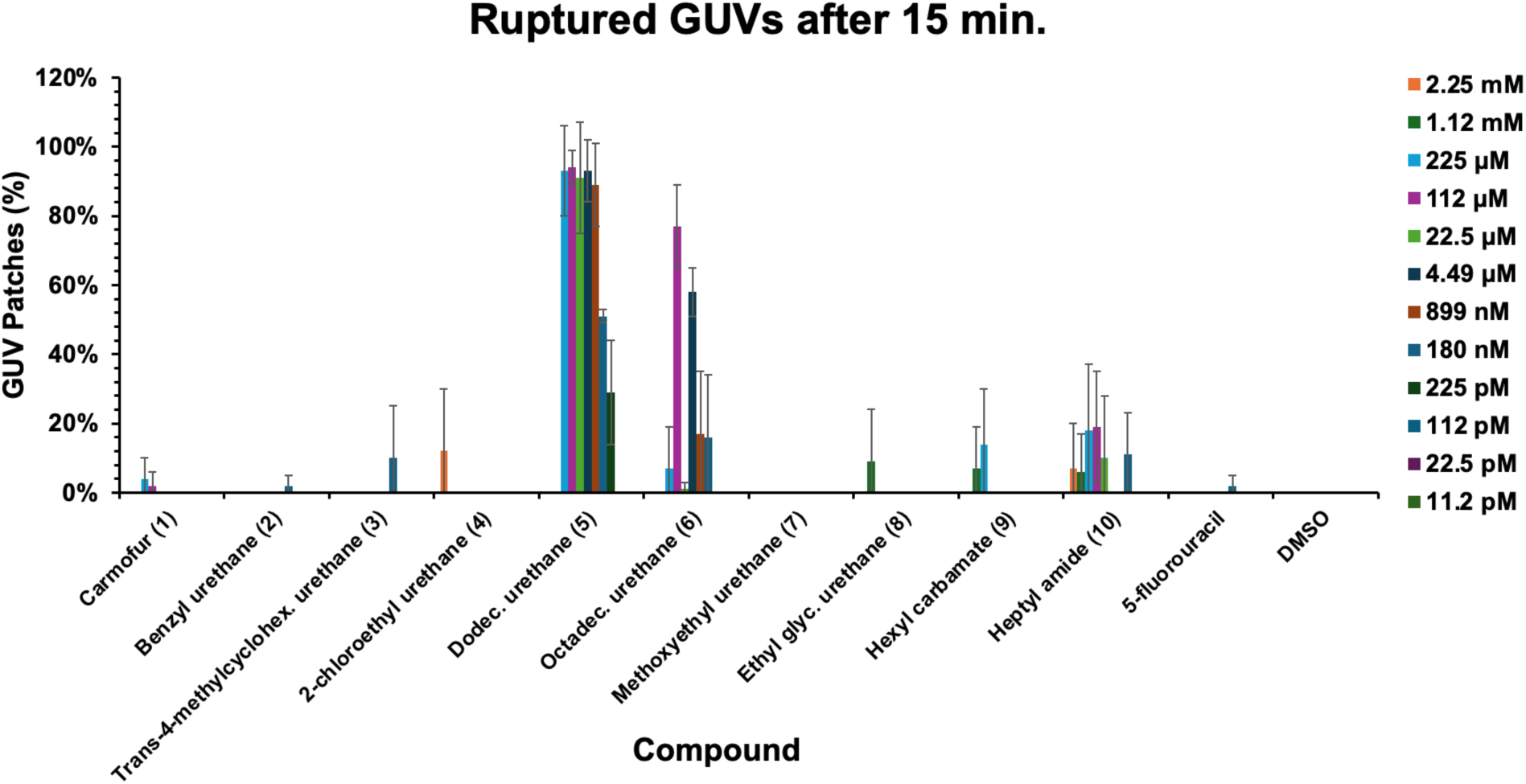
Bar plots of the percentage of GUVs that converted to patches. Average percent of GUV patches formed after a 15 minute incubation with each of the compounds is shown in each bar (n=3). Data is represented as the average ± SD and can be compared with the control (DMSO). In the space between each of the x-axis tick points the concentrations can be read from highest (left) to lowest (right) and are designated by color as shown in the legend. Note: Compounds **5** and **6** were poorly soluble at final concentrations of 2.25 mM and 1.12 mM and were hence excluded at those concentrations from this report

Overall, the bar plots indicate that the lipidated analogs of carmofur result in much higher bursting of GUVs into patches with a decline in bursting activity starting at concentrations around 5 μM but some extension into the picomolar range. Specifically, dodecyl urethane **5** causes the rupture of 29 ± 15 % of the GUVs at a concentration as low as 225 pM and octadecyl urethane **6** causes the rupture of 58 ± 7.0 % of the GUVs at a concentration of 4.49 μM. The quantitative data on the remaining analogs overall supports our qualitative observation that little to no membrane patches form at even the highest soluble concentrations for the other analogs with again the exception of heptyl amide **10** where some percentage of the GUVs do appear to be bursting into patches at the higher concentrations.

## Conclusion

In summary, the present work describes the preparation of a selection of 5-FU derivatives with different alkyl, aryl, heteroalkyl, and single-atom substituted side chains. In a cell free M^pro^ inhibition assay, we identified hexyl carbamate **9** as a potential lead inhibitor, with over two-fold improved M^pro^ inhibitory activity over carmofur **1**, making it a promising candidate for further development as a dual-purpose therapeutic. Benzyl urethane **2,** methoxyethyl urethane **7,** and heptyl amide **10** performed similarly to carmofur **1**, suggesting that the structure-activity relationship around carmofur-analogous SARS-CoV-2 M^pro^ inhibitors involves factors beyond mere electrophilicity. Other compounds in this series exhibited M^pro^ inhibitory activities significantly less potent than **1.**

When evaluated for antiproliferative activity in a selection of four human cell lines, including two model colorectal cancer cell lines HCT-116 and HT-29, a model breast cancer cell line MDA-MB-468, and a noncancerous HEK-293 embryonic kidney cell line, all compounds exhibited antiproliferative activities that were amplified at the 72-hour time point with minimal antiproliferative activity at 24 hours. None of the compounds exhibited improved antiproliferative potency over carmofur **1** at 24 hours, and they were generally more potent in inhibiting HEK-293 cells over colon or breast cancer cell lines in a dose-dependent fashion.

Finally, in efforts to probe the potential mechanism of action of our most lipophilic analogs **5** and **6** as membrane surfactants, we developed a cell-free assay to measure dose-dependent rupture of lipid membranes, modeled by fluorescently labeled GUVs. Compounds **5** and **6** exhibited consistent membrane rupture activity due to their long lipophilic side chains enhancing membrane-disruption properties, which suggests potential further research into their use as surfactant-like agents. This is consistent with the modest *in vitro* antiproliferative activity exhibited by **5** and **6** only observed at the highest evaluated soluble concentrations, which might occur through membrane disruption analogously to what is observed in the model GUV system. Other compounds in this series exhibited minimal to no membrane rupture activity.

The unique activities of the analogs described here provide key structural cues to the diverse biological and biochemical activity of carmofur and its analogs, which builds upon the foundation of known 5-fluorouracil derivatives that will inspire the design and preparation of other carmofur analogs. Namely, the remarkable potency of lipidated 5-fluorouracil derivatives in membrane rupture provides insight into a new potential application of long chain aliphatic 5-FU derivatives that prompts more detailed biophysical examination of these and other similar membrane-disrupting agents, which shall be reported in due course of time.

## Experimental Section

### General synthesis procedure

To a 50 mL oven-dried three-neck flask fitted with a water-jacketed condenser and a Teflon stir bar was added 5-fluorouracil (200.0 mg, 1.54 mmol, 1.0 eq) under nitrogen, and then anhydrous NMP (12.00 mL, 0.13 M) was added via syringe followed by a suitable base Et_3_N or DBU (0.23 mmol, 0.15 eq) and the appropriate isocyanate, chloroformate, or acyl chloride (2.31 mmol, 1.5 eq).The reaction was stirred at 50–90 °C and monitored by ^19^F NMR (0.5–3 h). Upon completion, the reaction mixture was extracted in ethyl acetate (2×50 mL) over brine. The combined organic layers were dried over anhydrous magnesium sulfate and were purified via column chromatography with a hexanes and ethyl acetate eluent. Spectra and detailed experimental procedures are described in the <u>Supporting Information</u>.

### Other chemicals

Carmofur and 5-fluorouracil were purchased from AK Scientific. All solvents and reagents were obtained from commercial suppliers and were used without further purification.

### Characterization of synthesized carmofur analogs

Benzyl urethane **(2)**: Yield: 34 %; GC-MS recorded: 263.222 [M^+^], expected for [C_12_H_10_FN_3_O_3_]^+^: 263.228 [M^+^]; R_*f*_= 0.72 (60 % EtOAc:Hexanes); FT-IR (ATR, cm^-1^): 3299, 3089, 2866, 2799, 2355, 1773, 1743, 1715, 1693, 1651, 1607, 1585, 1520, 1496, 1464, 1455, 1335, 1281, 1264, 1233, 1206, 1113, 1079, 1048, 1028, 1001, 910, 834, 757, 730, 694, 667, 622, 614, 598, 561; ^1^H NMR (500 MHz, Acetone-*d*6) δ 10.92 (s, 1H), 9.57 (s, 1H), 8.42 (d, J = 7.6 Hz, 1H), 7.39 – 7.23 (m, 5H), 4.57 (d, J = 3.9 Hz, 2H); ^13^C NMR (126 MHz, Acetone-*d*6) δ 150.64, 150.04, 142.19, 140.32, 138.33, 128.57, 127.63, 127.40, 122.68, 122.38, 44.48, 44.36; ^19^F NMR (57 MHz, Acetone-*d*6) δ −166.31 (d, *J* = 7.6 Hz).

*Trans*-4-methylcyclohexyl urethane **(3)**: Yield: 56 %; GC-MS recorded: 269.278 [M^+^], expected for [C_12_H_16_FN_3_O_3_]^+^: 269.276 [M^+^]; R_*f*_= 0.60 (60 % EtOAc:Hexanes); FT-IR (ATR, cm^-1^): 3326, 3286, 3079, 3037, 2952, 2927, 2904, 2865, 2818, 1755, 1724, 1698, 1683, 1652, 1624, 1576, 1539, 1507, 1439, 1420, 1377, 1362, 1339, 1311, 1267, 1236, 1213, 1197, 1156, 1106, 1079, 1045, 1005, 968, 953, 901, 883, 857, 795, 766, 749, 736, 705, 639, 621, 595, 569, 562; ^1^H NMR (400 MHz, DMSO-*d*6) δ 11.48 (s, 1H), 9.03 (d, *J* = 7.3 Hz, 1H), 8.31 (d, *J* = 7.4 Hz, 1H), 7.69 (d, *J* = 6.0 Hz, 1H), 3.39 – 3.28 (m, 9H), 1.86 (d, *J* = 12.4 Hz, 2H), 1.64 (d, *J* = 13.3 Hz, 2H), 1.33 – 1.17 (m, 3H), 0.96 (q, *J* = 10.2 Hz, 2H), 0.82 (t, *J* = 6.5 Hz, 3H); ^13^C NMR (101 MHz, DMSO-*d*6) δ 150.97, 149.50, 149.07, 142.57, 126.55, 123.19, 122.82, 50.48, 33.60, 32.34, 31.56, 22.38; ^19^F NMR (57 MHz, Chloroform-*d*) δ −162.65 (d, *J* = 6.2 Hz). 2-Chloroethyl urethane **(4)**: Yield: 57 %; GC-MS recorded: 235.607 [M^+^], expected for [C_7_H_7_ClFN_3_O_3_]^+^: 235.599 [M^+^]; R_*f*_ = 0.60 (30 % EtOAc:Hexanes); FT-IR (ATR, cm^-1^): 3274, 3089, 2929, 2836, 1722, 1691, 1667, 1515, 1456, 1435, 1334, 1267, 1247, 1222, 1205, 1188, 1145, 1101, 1074, 1061, 1036, 943, 883, 872, 846, 786, 758, 742, 655, 609, 581, 572, 564, 553; ^1^H NMR (400 MHz, Acetone-*d*6) δ 10.84 (s, 1H), 9.35 (s, 1H), 8.30 (d, *J* = 7.4 Hz, 1H), 3.68 – 3-60 (m, 4H); ^13^C NMR (101 MHz, Acetone-*d*6) δ 150.49, 150.02, 142.40, 140.05, 122.43, 122.05, 42.71; ^19^F NMR (57 MHz, Chloroform-*d*) δ −161.94 (d, *J* = 4.8 Hz). Spectroscopic data is consistent with previously reported values [36].

Dodecyl urethane **(5)**: Yield: 44 %; GC-MS recorded: 341.411 [M^+^], expected for [C_17_H_28_FN_3_O_3_]^+^: 341.427 [M^+^]; R_*f*_ = 0.90 (60 % EtOAc:Hexanes); FT-IR (ATR, cm^-1^): 3335, 3287, 3099, 3053, 2959, 2919, 2847, 2360, 2344, 1741, 1729, 1683, 1669, 1636, 1612, 1574, 1537, 1462, 1431, 1377, 1362, 1331, 1273, 1239, 1220, 1195, 1168, 1099, 1043, 1026, 932, 863, 838, 805, 778, 760, 751, 742, 724, 668, 630, 615, 600, 581, 570, 564, 553; ^1^H NMR (400 MHz, Chloroform-*d*) δ 9.03 (t, *J* = 4.9 Hz, 1H), 8.49 (d, *J* = 6.7 Hz, 1H), 3.39 (q, *J* = 7.4 Hz, 2H), 1.60 (p, *J* = 7.2 Hz, 2H), 1.42 – 1.17 (m, 22H), 0.88 (t, *J* = 6.9 Hz, 3H); ^13^C NMR (101 MHz, Chloroform-*d*) δ 156.96, 156.68, 150.07, 149.09, 141.99, 139.58, 123.47, 123.10, 41.50, 31.93, 29.65, 29.64, 29.58, 29.51, 29.36, 29.21, 29.15, 26.82, 22.70, 14.12; ^19^F NMR (57 MHz, Chloroform-*d*) δ −162.53 (d, *J* = 7.6 Hz). Spectroscopic data is consistent with previously reported values [37].

Octadecyl urethane **(6)**: Yield: 19 %; GC-MS recorded: 425.598 [M^+^], expected for [C_23_H_40_FN_3_O_3_]^+^: 425.589 [M^+^]; R_*f*_ = 0.90 (50 % EtOAc:Hexanes); FT-IR (ATR, cm^-1^): 2959, 2918, 2847, 2361, 2344, 1746, 1723, 1684, 1669, 1654, 1647, 1636, 1576, 1538, 1507, 1497, 1489, 1462, 1435, 1375, 1338, 1274, 1220, 1197, 1099, 1046, 1031, 1022, 932, 899, 864, 823, 801, 781, 760, 753, 744, 724, 634, 616, 599, 592, 580, 571, 564, 553; ^1^H NMR (400 MHz, Pyridine-*d*5) δ 9.61 (t, *J* = 5.6 Hz, 1H), 8.75 (s, 1H), 3.48 (q, *J* = 6.1 Hz, 2H), 1.62 (p, *J* = 6.9 Hz, 2H), 1.39 – 1.23 (m, 34H), 0.88 (t, *J* = 7.2 Hz, 3H); ^13^C NMR (101 MHz, Pyridine-*d*5) δ 157.91, 157.64, 151.47, 150.09, 140.41, 138.06, 122.72, 122.58, 41.18, 31.89, 31.56, 29.76, 29.75, 29.71, 29.69, 29.64, 29.58, 29.43, 29.38, 29.29, 26.91, 22.70, 14.05; ^19^FNMR (101 MHz, Pyridine-*d*5) δ 157.91, 157.64, 151.47, 150.09, 140.41, 138.06, 122.72, 122.58, 41.18, 31.89, 31.56, 29.76, 29.75, 29.71, 29.69, 29.64, 29.58, 29.43, 29.38, 29.29, 26.91, 22.70, 14.05. Spectroscopic data is consistent with previously reported values [37].

Methoxyethyl urethane **(7)**: Yield: 25 %; HRMS recorded: 231.2100 [M^+^], expected for [C_8_H_10_FN_3_O_4_]^+^: 231.1834 [M^+^]; R_*f*_= 0.60 (30 % EtOAc:Hexanes); FT-IR (ATR, cm^-1^): 3290, 3163, 3072, 2934, 2898, 2866, 2824, 2359, 2342, 1728, 1694, 1559, 1522, 1507, 1496, 1455, 1388, 1362, 1350, 1339, 1317, 1268, 1227, 1218, 1190, 1114, 1103, 1087, 1066, 1019, 956, 927, 897, 868, 833, 781, 752, 743, 676, 609, 567, 556; ^1^H NMR (400 MHz, DMSO-*d*6) δ 11.51 (s, 1H), 10.74 (s, 1H), 7.76 (d, *J* = 6.1 Hz, 1H), 5.97 (t, *J* = 5.7 Hz, 1H), 3.29 (t, *J* = 5.6 Hz, 2H), 3.24 (s, 3H), 3.13 (q, *J* = 5.6 Hz, 2H); ^13^C NMR (101 MHz, DMSO-*d*6) δ 158.54, 158.39, 158.28, 150.54, 141.43, 139.17, 126.93, 126.61, 72.09, 58.34, 39.45; ^19^F NMR (57 MHz, Acetone-*d*6) δ −171.50 (d, *J* = 6.1 Hz).

Ethyl glycinate urethane **(8)**: Yield: 20 %; GC-MS recorded: 259.208 [M^+^], expected for [C_9_H_10_FN_3_O_5_]^+^: 259.193 [M^+^]; R_*f*_= 0.60 (30 % EtOAc:Hexanes); FT-IR (ATR, cm^-1^): 3318, 3074, 2995, 2832, 2358, 2333, 1731, 1715, 1693, 1660, 1510, 1474, 1445, 1397, 1380, 1346, 1264, 1245, 1236, 1195, 1178, 1122, 1065, 1012, 943, 867, 797, 768 727, 657, 612, 578, 563, 553; ^1^H NMR (400 MHz, DMSO-*d*6) δ 11.01 (s, 1H), 7.70 (d, *J* = 6.1 Hz, 1H), 6.45 (t, *J* = 5.9 Hz, 1H), 4.05 (q, *J* = 7.1 Hz, 2H), 3.74 (d, *J* = 6.1 Hz, 2H), 1.15 (t, *J* = 7.1 Hz, 3H); ^13^C NMR (101 MHz, DMSO-*d*6) δ 192.20, 177.29, 171.36, 159.03, 158.26, 142.84, 124.06, 122.80, 60.61, 42.17, 14.51; ^19^F NMR (57 MHz, DMSO-*d*6) δ −171.48 (d, *J* = 7.6 Hz).

Hexyl carbamate **(9)**: Yield: 93 %; GC-MS recorded: 258.243 [M^+^], expected for [C_11_H_15_FN_2_O_4_]^+^: 258.249 [M^+^]; R_*f*_= 0.40 (75 % EtOAc:Hexanes); FT-IR (ATR, cm^-1^): 3201, 3097, 2954, 2933, 2872, 2849, 1739, 1722, 1683, 1465, 1436, 1388, 1341, 1286, 1228, 1195, 1122, 1099, 1042, 1011, 989, 916, 899, 842, 799, 768, 735, 696, 607, 580, 573, 563, 553; ^1^H NMR (400 MHz, Acetone-*d*6) δ 10.48 (s, 1H), 8.03 (d, *J* = 7.1 Hz, 1H), 4.25 (t, *J* = 6.6 Hz, 2H), 1.63 (p, *J* = 7.8 Hz, 2H), 1.33 (p, *J* = 7.5 Hz, 2H), 1.24 – 1.18 (m, 4H), 0.76 (t, *J* = 6.9 Hz, 3H); ^13^C NMR (101 MHz, Acetone-*d*6) δ 150.27, 149.99, 145.63, 124.13, 123.76, 68.99, 31.13, 28.11, 25.11, 22.24, 13.32; ^19^F NMR (57 MHz, Chloroform-*d*) δ −162.06 (d, *J* = 6.1 Hz).

Heptyl amide **(10)**: Yield: 53 %; GC-MS recorded: 243.250 [M^+^], expected for [C_11_H_15_FN_2_O_3_]^+^: 242.250 [M^+^]; R_*f*_= 0.80 (60 % EtOAc:Hexanes); FT-IR (ATR, cm^-1^): 3188, 3108, 3065, 2955, 2925, 2848, 1727, 1682, 1559, 1454, 1431, 1395, 1374, 1339, 1318, 1295, 1281, 1259, 1226, 1128, 1098, 1034, 1018, 928, 851, 796, 775, 755, 742, 721, 643, 604, 564, 552; ^1^H NMR (400 MHz, Chloroform-*d*) δ 8.56 (s, 1H), 8.22 (d, *J* = 6.7 Hz, 1H), 3.05 (t, *J* = 7.3 Hz, 2H), 1.65 (p, *J* = 7.1 Hz, 2H), 1.35 – 1.22 (m, 6H), 0.83 (t, *J* = 6.9 Hz, 3H); ^13^C NMR (101 MHz, Chloroform-*d*) δ 219.78, 171.99, 147.63, 121.90, 121.54, 39.05, 31.47, 28.59, 24.37, 22.47, 14.00; ^19^F NMR (57 MHz, Chloroform-*d*) δ −161.82 (d, *J* = 6.1 Hz).

### Main protease colorimetric assay

To a 96-well flat-bottom plate (Corning Costar) was added 10 μL of compounds **1** to **10** diluted to 25 μM in 4 % DMSO and buffer prepared according to manufacturer instructions for Cayman Chemical SARS-CoV-2 Main Protease Assay Buffer (Item No. 701961), 0.25 μL DMSO, and 15 μL of 10X SARS-CoV-2 Main Protease (recombinant) solution (Cayman Chemical, Item No. 701963, Batch 0673677-1) were added in triplicate. After incubation at 37 °C with 5 % CO_2_ for an hour, 25 μL of M^pro^ chromogenic substrate peptide TSAVLQ-pNA (Sigma Aldrich, SAE0172, Lot No. 0000159244) at 800 μg/mL was added to all wells. A negative control was established with the addition of only substrate, buffer, and DMSO, and the positive control contained the substrate, buffer, M^pro^ enzyme, and DMSO. Three spectrometric readings of the plate were obtained before and after adding the substrate using a Labsystems Multiskan plate reader at 410 nm over 160 minutes with 20 minute timepoints. Percent M^pro^ activity was calculated by dividing the drugged wells’ absorbance values relative to the negative control over those of the positive control wells.

### Molecular docking

Computer models of **1** through **10** were performed on the Schrödinger small molecule discovery suite following methods previously described by the Hoye group [38]. Thermodynamically acceptable three-dimensional conformations were identified through a Monte Carlo conformational search using MacroModel with the OPLS2005 force field and standard precision [39–41]. Following this, each structure was further DFT optimized at the B3LYP / 6-31G(d,p)++ level of theory (Jaguar) with an implicit aqueous solvation model (PCM) [42]. X-ray structures of SARS-CoV-2 main protease (PDB: 7CAM and 7BUY) and human acid ceramidase (PDB: 6MHM) were prepared using the protein preparation wizard, and structures of **1** through **10** were docked with standard parameters on Schrödinger Glide [9,16,27–31,43].

### Cell culture

Human colorectal cancer (HCT-116 and HT-29) cell lines were obtained from the European Collection of Authenticated Cell Cultures (ECACC) and maintained in McCoy’s 5A Media (Tribioscience; Sunnyvale, CA) supplemented with 10 % fetal bovine serum (FBS) from Gibco and 1 % penicillin-streptomycin (Tribioscience; Sunnyvale, CA). Human breast cancer (MDA-MB-468) and human embryonic kidney (HEK-293) cell lines were obtained from the American Type Culture Collection (ATCC) and maintained in Dulbecco’s Modified Eagle’s Medium (DMEM) from Tribioscience (Sunnyvale, CA) supplemented with 10 % fetal bovine serum and 1 % penicillin-streptomycin. All cells were cultured in 25 cm^2^ and 75 cm^2^ flasks in a humidified incubator at 37 °C with 5 % CO_2_.

### MTT cell viability assay

HCT-116, HT-29, MDA-MB-468, and HEK-293 cells were seeded at 24,000 cells per well in 96-well plates (Thermo Scientific Nunclon Delta Surface). 5-Fluorouracil and compounds **1** to **10** were added in dimethylsulfoxide (DMSO, final concentration of 0.5 % v/v) in triplicate. A negative control was established by dosing cells with undrugged, 0.5% v/v DMSO. After incubation for 24 or 72 hours, a mixture of MTT (3-(4,5-dimethylthiazol-2-yl)-2,5-diphenyl-2H-tetrazolium bromide) and 1X phosphate buffered saline (PBS) (Tribioscience; Sunnyvale, CA) at 5 mg/mL was prepared and added to each well. The cells were then incubated for 1 hour, and the cell medium and MTT solution were aspirated before 100 μL of DMSO was added to each well. The plate was incubated for an additional 15 minutes, and three spectrometric readings were obtained with a Molecular Devices SPECTRAmax 250 Microplate Spectrophotometer at 570 nm. Cell viability was calculated by taking the ratio of the absorbance values of the drugged well over those of the negative control, and results are reported as an average and standard deviation of triplicate experiments.

### GUV rupture fluorescence imaging assay

The GUVs were assembled based on literature procedures using a 1 mg/mL mixture of DOPC:RhodamineDPPE (99.5:0.5 mol %) [32]. To prepare the samples for imaging we passivated the wells of a 96-well glass bottom plate (Whatman 96-Well Glass Bottom Plate) with 1 mg/mL casein solution in 1X PBS. After incubation for 1 hour, 85 μL of 95 mM glucose was added to each of the wells and then added 2 μL of the GUV solution in 100 mM sucrose. We allowed the GUVs to sediment for 1 hour and collected images of the GUVs at 555 nm using a Zeiss Axiovert 200M fluorescence microscope with a Zeiss A-plan 10x/0.25 Ph1 M27 objective and a Zeiss FLUAR 5x/0.25 M27 objective. 2 μL of 5-fluorouracil and compounds **1** to **10** at the appropriate concentration were subsequently added in triplicate. A negative control was established with 0.0225 % v/v DMSO. Following a 15 minute incubation, we collected representative images of the GUVs in each well to characterize membrane disruption. The percent of GUV patches was determined by counting the number of patches in a 650 μm × 650 μm region and dividing by the total number of GUVs before rupture.

### Statistical analysis

All statistical analyses were performed using GraphPad Prism 10.3.1. Data in each group were compared using a two-tailed unpaired student’s t test. P-values <0.05 were considered statistically significant.

## Author Contributions

T.G., A.L., X.W., L.Z., L.X., Y.L., A.R., and E.N. synthesized compounds reported in this manuscript. T.G., A.L., X.W., N.B., L.Z., L.X., Y.L., A.R., J.V., C.L., and E.N. developed research methodology and performed reaction condition screening. T.G. and A.L. performed the SARS-CoV-2 Mpro assay. N.B. performed computer modeling and docking studies. T.G., N.B., L.Z., A.L., L.X., and E.N. performed antiproliferative assays (MTT) on cell lines. K.S., K.S., and J.P. performed GUV imaging experiments. J.P. developed GUV imaging workflow, and K.S. performed GUV membrane rupture quantification. T.G., N.B., K.S., J.P., and E.N. prepared the manuscript figures and wrote the manuscript text. All authors reviewed the manuscript.

## Acknowledgements

The authors gratefully acknowledge the Stanford University Nuclear Magnetic Resonance facility for providing facilities for high field NMR. Additionally, the authors acknowledge Tribioscience Inc. (Sunnyvale, CA) for support of the materials used in this research and Nanalysis Corporation (Calgary, AB) for support of benchtop NMR capabilities that enabled this work.

## Conflict of Interest

The authors declare no conflicts of interest.

**Supporting information** https://1drv.ms/w/c/4c28190939c62071/EfBMKRgYFapItPLCY4zYttsBOalE9M5OYK1eX8ZE6IKNuw Characterization and spectroscopic data for all compounds, supplementary figures, and computer modeling information

## References

1. Shelton J, Lu X, Hollenbaugh JA, Cho JH, Amblard F, Schinazi RF. Metabolism, Biochemical Actions, and Chemical Synthesis of Anticancer Nucleosides, Nucleotides, and Base Analogs. Chem Rev. 2016;116(23):14379–455. 10.1021/acs.chemrev.6b00209.

2. Galmarini MC, Mackey RJ, Dumontet C. Nucleoside analogues and nucleobases in cancer treatment. The Lancet Oncol. 2002;3(7):415–24. 10.1016/S1470-2045(02)00788-X.

3. Seley-Radtke KL, Yates MK. The evolution of nucleoside analogue antivirals: A review for chemists and non-chemists. Part 1: Early structural modifications to the nucleoside scaffold. Antiviral Res. 2018;154:66–86. 10.1016/j.antiviral.2018.04.004.

4. Yates MK, Seley-Radtke KL. The evolution of antiviral nucleoside analogues: A review for chemists and non-chemists. Part II: Complex modifications to the nucleoside scaffold. Antiviral Res. 2019;162:5–21. 10.1016/j.antiviral.2018.11.016.

5. Geraghty RJ, Aliota MT, Bonnac LF. Broad-spectrum antiviral strategies and nucleoside analogues. Viruses. 2021;13(4):667. 10.3390/v13040667.

6. Islam MM, Mirza SP. Versatile use of Carmofur: A comprehensive review of its chemistry and pharmacology. Drug Dev Res. 2022;83(7):1505–18. 10.1002/ddr.21984.

7. Nishiyama M, Takagami S, Kim R, Kirihara Y, Saeki T, Jinushi K, et al. [Inhibition of thymidylate synthetase and antiproliferative effect by 1-hexylcarbamoyl-5-fluorouracil]. Gan To Kagaku Ryoho. 1988;15(11):3109–13.

8. Realini N, Solorzano C, Pagliuca C, Pizzirani D, Armirotti A, Luciani R, et al. Discovery of highly potent acid ceramidase inhibitors with in vitro tumor chemosensitizing activity. Sci Rep. 2013;3. 10.1038/srep01035.

9. Dementiev A, Joachimiak A, Nguyen H, Gorelik A, Illes K, Shabani S, et al. Molecular Mechanism of Inhibition of Acid Ceramidase by Carmofur. J Med Chem. 2019;62(2):987–92. https://pubs.acs.org/doi/10.1021/acs.jmedchem.8b01723.

10. Pizzirani D, Pagliuca C, Realini N, Branduardi D, Bottegoni G, Mor M, et al. Discovery of a new class of highly potent inhibitors of acid ceramidase: Synthesis and structure-activity relationship (SAR). J Med Chem. 2013;56(9):3518–30. 10.1021/jm301879g.

11. Govindarajah N, Clifford R, Bowden D, Sutton PA, Parsons JL, Vimalachandran D. Sphingolipids and acid ceramidase as therapeutic targets in cancer therapy. Crit Rev Oncol Hematol. 2019;138:104–11. 10.1016/j.critrevonc.2019.03.018.

12. Vijayan Y, Lankadasari MB, Harikumar KB. Acid ceramidase: a novel therapeutic target in cancer. Curr Top Med Chem. 2019;19(17):1512–20. 10.2174/1568026619666190227222930.

13. Jin Z, Du X, Xu Y, Deng Y, Liu M, Zhao Y, et al. Structure of Mpro from SARS-CoV-2 and discovery of its inhibitors. Nature. 2020;582(7811):289–93. 10.1038/s41586-020-2223-y.

14. Wang MY, Zhao R, Gao LJ, Gao XF, Wang DP, Cao JM. SARS-CoV-2: Structure, Biology, and Structure-Based Therapeutics Development. Front Cell Infect Microbiol. 2020;10:587269. 10.3389/fcimb.2020.587269.

15. Zhu Y. SARS-CoV-2: phylogenetic status, mutations and therapeutic research based on spike protein. Eur Rev Med Pharmacol Sci. 2021;25(18):5843–52. 10.26355/eurrev_202109_26803.

16. Jin Z, Zhao Y, Sun Y, Zhang B, Wang H, Wu Y, et al. Structural basis for the inhibition of SARS-CoV-2 main protease by antineoplastic drug carmofur. Nat Struct Mol Biol. 2020;27:529–32. 10.1038/s41594-020-0440-6.

17. Yang H, Rao Z. Structural biology of SARS-CoV-2 and implications for therapeutic development. Nat Rev Microbiol. 2021;19:685–700. 10.1038/s41579-021-00630-8.

18. Kang KM, Jang Y, Lee SS, Jin MS, Jun CD, Kim M, et al. Discovery of antiviral SARS-CoV-2 main protease inhibitors by structure-guided hit-to-lead optimization of carmofur. Eur J Med Chem. 2023;260:115720. 10.1016/j.ejmech.2023.115720.

19. Chen R, Singh P, Su S, Kocalar S, Wang X, Mandava N, et al. Benchtop 19F Nuclear Magnetic Resonance (NMR) Spectroscopy Provides Mechanistic Insight into the Biginelli Condensation toward the Chemical Synthesis of Novel Trifluorinated Dihydro- and Tetrahydropyrimidinones as Antiproliferative Agents. ACS Omega. 2023;8(11):10545–54. https://pubs.acs.org/doi/10.1021/acsomega.3c00290.

20. Chyu A, Xi S, Kim J, et al. Benchtop 19F NMR Spectroscopy optimized Knorr pyrazole synthesis of Celecoxib and Mavacoxib, 3-(trifluoromethyl) pyrazolyl benzenesulfonamides Non-Steroidal Anti-Inflammatory Drugs (NSAIDs). ChemRxiv. 2024; This content is a preprint and has not been peer-reviewed. 10.26434/chemrxiv-2024-d1grv.

21. Wang X, Vu J, Luk C, Njoo, E. Benchtop ^19^F nuclear magnetic resonance spectroscopy enabled kinetic studies and optimization of the synthesis of carmofur. Can J Chem. 2023;101(8):518–24. 10.1139/cjc-2022-0266.

22. Ozaki S, Kong X, Watanabe Y, et al. 5-Fluorouracil Derivatives. XXII. Synthesis and Antitumor Activities of 1-Carbamoyl-5-fluorouracils. Chem Pharm Bull. 1997;45(8):1372–5. 10.1248/cpb.45.1372.

23. Hoshi A, Masaaki I, Nakamura A, Inomata M, Kuretani K. Antitumor Activity of 1-Alkylcarbamoyl Derivatives of 5-Fluorouracil against L1210 Leukemia. Chem Pharm Bull. 1978;26(1):161–5. 10.1248/cpb.26.161.

24. Zalupski M, Baker LH. Ifosfamide. J Natl Cancer Inst. 1988;80(8):556–66. 10.1093/jnci/80.8.556.

25. Mazur L, Opydo-Chanek M, Stojak M. Glufosfamide as a new oxazaphosphorine anticancer agent. Anticancer Drugs. 2011;22(6):488–93. 10.1097/cad.0b013e328345e1e0.

26. Thomsen CL, Thøgersen J, Keiding SR. Ultrafast Charge-Transfer Dynamics: Studies of p-Nitroaniline in Water and Dioxane. 1998; 102(7):1062–7. https://pubs.acs.org/doi/10.1021/jp972492g.

27. Wang Y-C, Yang W-H, Yang C-S, Hou M-H, Tsai C-L, Chou Y-Z, et al. Structural basis of SARS-CoV-2 main protease inhibition by a broad-spectrum anti-coronaviral drug. Am J Cancer Res. 2020;10(8):2535–45.

28. Friesner RA, Banks JL, Murphy RB, Halgren TA, Klicic JJ, Mainz DT, et al. Glide: A New Approach for Rapid, Accurate Docking and Scoring. 1. Method and Assessment of Docking Accuracy. J Med Chem. 2004;47(7):1739–49. 10.1021/jm0306430.

29. Halgren TA, Murphy RB, Friesner RA, Beard HS, Frye LL, Pollard WT, et al. Glide: A New Approach for Rapid, Accurate Docking and Scoring. 2. Enrichment Factors in Database Screening. J Med Chem. 2004;47(7):1750–9. 10.1021/jm030644s.

30. Friesner RA, Murphy RB, Repasky MP, Frye LL, Greenwood JR, Halgren TA, et al. Extra precision glide: Docking and scoring incorporating a model of hydrophobic enclosure for protein-ligand complexes. J Med Chem. 2006;49(21):6177–96. 10.1021/jm051256o.

31. Yang Y, Yao K, Repasky MP, Leswing K, Abel R, Shoichet BK, et al. Efficient Exploration of Chemical Space with Docking and Deep Learning. J Chem Theory Comput. 2021;17(11):7106–19. 10.1021/acs.jctc.1c00810.

32. Pazzi J, Subramaniam AB. Nanoscale Curvature Promotes High Yield Spontaneous Formation of Cell-Mimetic Giant Vesicles on Nanocellulose Paper. ACS Appl Mater Interfaces. 2020;12(50):56549–61. https://pubs.acs.org/doi/10.1021/acsami.0c14485.

33. Angelico R, Ceglie A, Cuomo F, Cardellicchio C, Mascolo G, Colafemmina G. Catanionic Systems from Conversion of Nucleotides into Nucleo-lipids. Langmuir. 2008;24(6):2348–55. https://pubs.acs.org/doi/10.1021/la702580j.

34. Khiati S, Pierre N, Andriamanarivo S, Grinstaff MW, Arazam N, Nallet F, et al. Anionic Nucleotide-Lipids for In Vitro DNA transfection. Bioconjug Chem. 2009;20(9):1765–72. https://pubs.acs.org/doi/10.1021/bc900163s.

35. Khiati S, Luvino D, Oumzil K, Chauffert B, Camplo M, Barthélémy P. Nucleoside-Lipid-Based Nanoparticles for Cisplatin Delivery. ACS Nano. 2011;5(11):8649–55. https://pubs.acs.org/doi/10.1021/nn202291k.

36. Inventors; Pyrimidine Derivative And Its Preparation. JPS5663966A. 1981 May 30.

37. Ozaki S, Ike Y, Mizuno H, Ishikawa K, Mori H, et al. 5-Fluorouracil Derivatives. I. The Synthesis of 1-Carbamoyl-5-fluorouracils. Bull Chem Soc Jpn. 1977;50(9):2406–12. 10.1246/bcsj.50.2406.

38. Willoughby PH, Jansma MJ, Hoye TR. A guide to small-molecule structure assignment through computation of (1 H and 13 C) NMR chemical shifts. Nat Protoc. 2014;9(3):643–60. 10.1038/nprot.2014.042.

39. Mohamadi F, Richards NGJ, Guida WC, Liskamp R, Lipton M, Caufield C, et al. MacroModel-An Integrated Software System for Modeling Organic and Bioorganic Molecules Using Molecular Mechanics. J Comput Chem. 1990;11(4):440–67. 10.1002/jcc.540110405.

40. Chen IJ, Foloppe N. Tackling the conformational sampling of larger flexible compounds and macrocycles in pharmacology and drug discovery. Bioorg Med Chem. 2013;21(24):7898–920. 10.1016/j.bmc.2013.10.003.

41. Watts KS, Dalal P, Tebben AJ, Cheney DL, Shelley JC. Macrocycle conformational sampling with macromodel. J Chem Inf Model. 2014;54(10):2680–96. 10.1021/ci5001696.

42. Bochevarov AD, Harder E, Hughes TF, Greenwood JR, Braden DA, Philipp DM, et al. Jaguar: A high-performance quantum chemistry software program with strengths in life and materials sciences. Int J Quantum Chem. 2013;113(18):2110–42. 10.1002/qua.24481.

43. Sastry GM, Adzhigirey M, Day T, Annabhimoju R, Sherman W. Protein and ligand preparation: Parameters, protocols, and influence on virtual screening enrichments. J Comput Aided Mol Des. 2013;27(3):221–34. 10.1007/s10822-013-9644-8.

