## Supporting Information Document for "Synthesis and Evaluation of Carmofur Analogs as Antiproliferative Agents, Inhibitors to the Main Protease (M^pro^) of SARS-CoV-2, and Membrane Rupture-Inducing Agents"

A. General Information SI-1
   a. Materials SI-1
   b. Equipment SI-1
   c. Computational Methods SI-1
B. Supplementary Figures SI-2
C. Experimental Procedures SI-3
   a. Kinetics SI-3
   b. $M^{pro}$ Colorimetric Assay Procedure SI-3
   c. Cell Culture SI-4
   d. GUV Rupture Assay Procedure SI-4
   e. Chemical Synthesis SI-5
      i. Benzyl urethane **(2)** SI-5
      ii. Trans-4-methylcyclohexyl urethane **(3)** SI-11
      iii. 2-Chloroethyl urethane **(4)** SI-16
      iv. Dodecyl urethane **(5)** SI-21
      v. Octadecyl urethane **(6)** SI-26
      vi. Methoxyethyl urethane **(7)** SI-31
      vii. Ethyl glycinate urethane **(8)** SI-36
      viii. Hexyl carbamate **(9)** SI-41
      ix. Heptyl amide **(10)** SI-46

### General Information:

#### A. Materials:

Solvents and reagents used in all reactions and purification processes were ACS grade or higher, were used without additional purification, and were purchased from Sigma Aldrich, Acros Organics, Oakwood Chemical, or AK Scientific. Deuterated solvents were purchased from Cambridge Isotope Laboratories, Acros Organics, or Martek Isotopes and were used without further purification. All other reagents, catalysts, and chemicals were purchased from commercial sources and used without further purification unless otherwise stated.

#### B. Equipment:

$^{1}$H, $^{19}$F, $^{13}$C{$^{1}$H} NMR spectra were acquired on a Varian INOVA 400 MHz nuclear magnetic resonance spectrometer, a Bruker Avance Neo 400 MHz nuclear magnetic resonance spectrometer, a JEOL 500 MHz nuclear magnetic resonance spectrometer, or a Nanalysis NMReady 60Pro 60 MHz multinuclear benchtop nuclear magnetic resonance spectrometer and were processed on the MestreNova software package. $^{1}$H, $^{19}$F, and $^{13}$C chemical shifts are reported in parts per million (ppm) relative to the residual solvent peak (Chloroform-*d* = 7.26 ppm, Acetone-*d*6 = 2.04 ppm, Pyridine-*d*5 = 8.74 ppm, DMSO-*d*6 = 2.49 ppm) as follows: chemical shift (δ), multiplicity (app = apparent, b = broad, s = singlet, d = doublet, t = triplet, q = quartet, m = multiplet, or combinations thereof), coupling constant(s) in Hz, integration. $^{13}$C chemical shifts are reported relative to the residual solvent peak (Chloroform-*d* = 77.0 ppm, Acetone-*d*6 = 29.8 ppm, Pyridine-*d*5 = 123.5 ppm, DMSO-*d*6 = 39.52 ppm). Mass spectra were obtained using a Thermo Scientific LTQ-XL linear ion trap mass spectrometer equipped with a Thermo Surveyor high performance liquid chromatography system (LC-MS), a Waters Quattro Micromass triple quadrupole mass spectrometer equipped with a Waters Alliance 1300 series high performance liquid chromatography system (LC-MS), or a DSQ single quadrupole mass spectrometer equipped with a Thermo Scientific Finnigan Trace GC Ultra Multi-channel gas chromatograph (GC-MS). Infrared spectra were collected on a Thermo Scientific Nicolet iS5 Fourier transform infrared (FT-IR) spectrometer equipped with a Thermo iD5 attenuated total reflectance (ATR) assembly.

#### C. Computational Methods:

All computer models for analogs evaluated in this study were performed on the Schrödinger small molecule discovery suite following methods described by the Hoye group [1]. Thermodynamically acceptable three-dimensional conformations of 5-fluorouracil analogs **1** through **10** were identified through a Monte Carlo conformational search using MacroModel with the OPLS2005 force field and standard precision [2–4]. Following this, each structure was further DFT optimized at the B3LYP / 6-31G(d,p)++ level of theory (Jaguar) with an implicit aqueous solvation model (PCM) [5]. X-ray structures of SARS-CoV-2 main protease (PDB:

7CAM and 7BUY) and human acid ceramidase (PDB: 6MHM) were prepared using the protein preparation wizard, and structures of **1** through **10** were docked with standard parameters with Schrodinger Glide [5–13].

### Supplementary Figures:

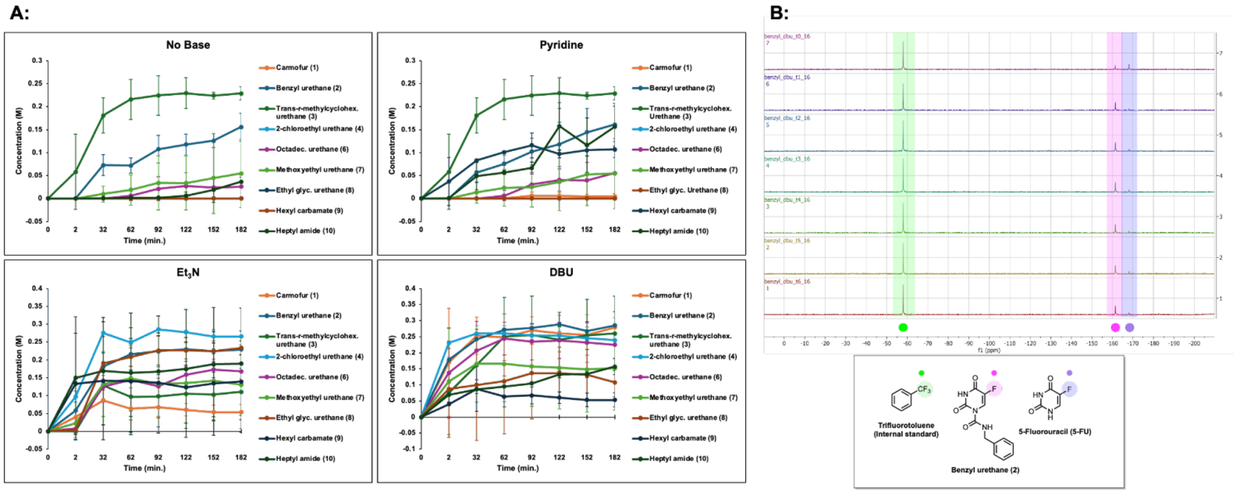


**S1 Benchtop $^{19}$F NMR enables quantitative reaction monitoring of carmofur analogs**
Concentrations of carmofur and its analogs were measured by benchtop $^{19}$F NMR every 30 minutes for up to 3 hours. B) Representative time course $^{19}$F NMR demonstrates peak to peak conversion of 5-fluorouracil (highlighted in purple) to its benzyl urethane analog 2 (labeled pink)

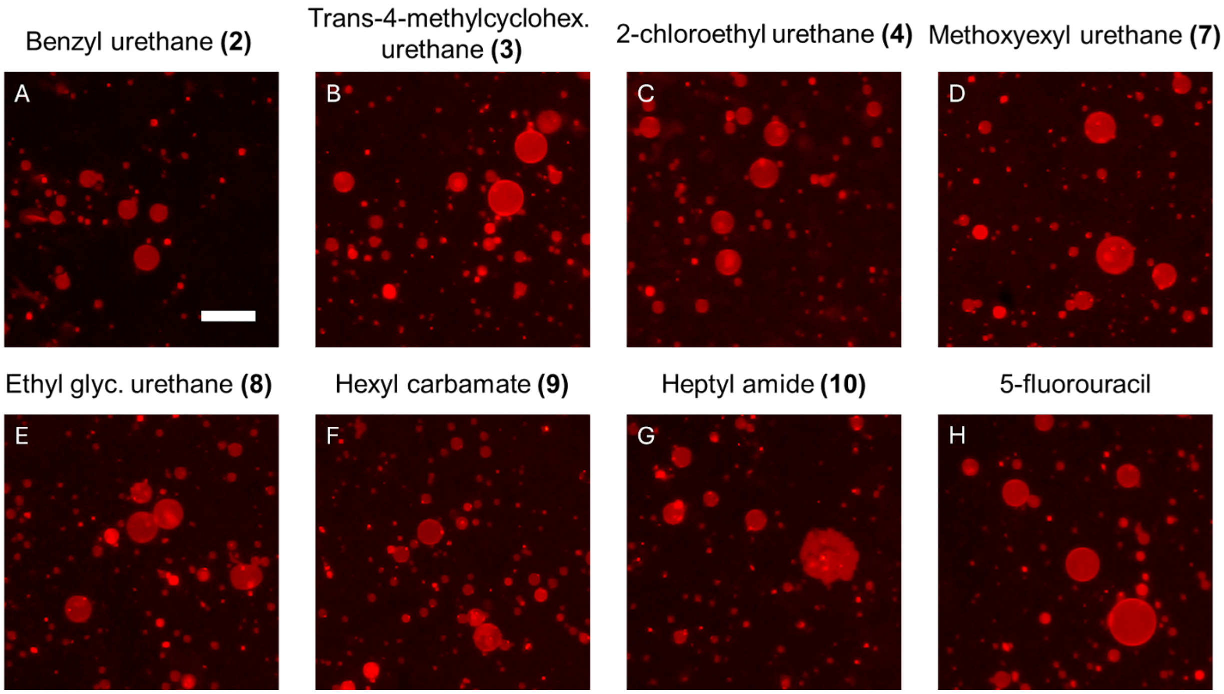

**S2 Membrane rupture assay using GUVs** Representative images of GUVs 15 minutes after addition of **A** 2.25 mM benzyl urethane **2**, **B** trans-4-methylcyclohex. urethane **3**, **C** 2-chloroethyl urethane **4**, **D** methoxyethyl urethane **7**, **E** ethyl glycinate urethane **8**, **F** hexyl carbamate **9**, **G** heptyl amide **10**, and **H** 5-fluorouracil. Scale bar for **A-H** is 50 μm

### Experimental Procedures:

#### A. Kinetics:

A stock solution of 0.3 M 5-fluorouracil and 0.3 M α,α,α-trifluorotoluene (Sigma Aldrich, 98 %) was prepared in anhydrous N-methylpyrrolidinone (NMP) (Sigma Aldrich) to which the appropriate base was added at 0.045 M: pyridine (Sigma Aldrich), triethylamine (Sigma Aldrich), or 1,8-diazabicyclo[5.4.0]undec-7-ene (AK Scientific). In addition, 0.45 M solutions of the isocyanates, chloroformates, and acyl chlorides were prepared in anhydrous NMP (Sigma Aldrich). To a 4 mm NMR tube was sequentially added the stock solution of 5-fluorouracil, base, and α,α,α-trifluorotoluene and the solution of the substituted isocyanate, chloroformate, or acyl chloride. An initial $^{19}F$ NMR using 16 scans and a 2.27 s scan delay was taken, and the NMR tube was subsequently placed into a water bath heated to 60 °C. $^{19}F$ NMR spectra were taken in 30 minute intervals over a 3 hour period, and the peaks were integrated against the internal standard α,α,α-trifluorotoluene (Sigma Aldrich, 98 %) to determine relative concentrations according to methods our laboratory has previously established [14].

#### B. $M^{pro}$ Colorimetric Assay Procedure:

$M^{pro}$ inhibitory activities were evaluated colorimetrically using a spectrophotometric method with a $M^{pro}$ chromogenic substrate peptide TSAVLQ-pNA (Sigma Aldrich, SAE0172, Lot 0000159244). Compounds **1** to **10** were diluted to a final concentration of 25 μM in 4 % DMSO and buffer prepared according to Cayman Chemical's SARS-CoV-2 Main Protease Assay Buffer (Item No. 701961). The lyophilized chromogenic substrate powder was diluted to 800 μg/mL with 100 mM dPBS, and the 100 mM SARS-CoV-2 Main Protease (recombinant) (Cayman Chemical, Item No. 701963, Batch 0673677-1) solution in HEPES was further diluted to a 10X working solution (5.95 mM) using 100 mM dPBS. To a 96-well flat-bottom plate (Corning Costar), 10 μL of each compound, 0.25 μL DMSO, and 15 μL of the $M^{pro}$ enzyme were added in triplicate. After the plate incubated at 37 °C with 5 % $CO_2$ for an hour, 25 μL of the substrate was then added to all wells. A negative control was established with the addition of only substrate, buffer, and 1 % DMSO, and a positive control contained the substrate, buffer, $M^{pro}$ enzyme, and 1 % DMSO. Three spectrometric readings of the plate were obtained before and after adding the substrate using a Labsystems Multiskan plate reader at 410 nm over 160 minutes with 20 minute timepoints. Percent $M^{pro}$ activity was then calculated using the ratio of the drugged wells' absorbance values relative to the negative control over those of the positive control wells.

#### C. Cell Culture:

Human colorectal cancer (HCT-116 and HT-29) cell lines were obtained from the European Collection of Authenticated Cell Cultures (ECACC) and maintained in McCoy's 5A Media (Tribioscience; Sunnyvale, CA) supplemented with 10 % fetal bovine serum (FBS) from Gibco and 1 % penicillin-streptomycin from Tribioscience (Sunnyvale, CA). Human breast cancer (MDA-MB-468) and human embryonic kidney (HEK-293) cell lines were obtained from the American Type Culture Collection and cultured in Dulbecco's Modified Eagle's Medium (DMEM) from Tribioscience supplemented with 10 % fetal bovine serum (FBS) and 1 % penicillin-streptomycin. All cells were grown to 80 % confluency in 25 $cm^2$ and 75 $cm^2$ flasks (Bio Basics) in a humidified incubator at 37 °C in 5 % $CO_2$.

#### D. MTT Procedure:

Carmofur and 5-fluorouracil obtained from AK Scientific and Compounds **2** to **10** were synthetically prepared (Section F) and serial diluted down to concentrations of 100 mM, 50 mM, 10 mM, 5 mM, 1 mM, 0.5 mM, and 0.1 mM. The HCT-116, HT-29, MDA-MB-468, and HEK-293 cells were grown to confluence at 37 °C with 5 % $CO_2$ and seeded at 40–60 % confluency in a sterile 96-well plate (Thermo Scientific Nunclon Delta Surface). Following a 24 hour incubation period, the drug medium solution was prepared through 200-fold dilution, and the cell media in each well was replaced with 100 or 150 µL of drug medium solution in triplicate. A negative control was established by dosing cells with undrugged, 0.5% v/v DMSO. The plates were incubated for 24 hours and 72 hours, and a mixture of thiazolyl blue tetrazolium bromide (MTT) from AK Scientific and phosphate buffered saline (PBS) from Tribioscience at 5 mg/mL was prepared. 10 µL of the MTT solution was added to each well. The cells were incubated for 1 hour, and the cell medium and MTT solution were aspirated before 100 µL of DMSO was added to each well. The plate was then incubated for another 15 minutes, and three spectrometric readings were obtained with a Molecular Devices SPECTRAmax 250 Microplate Spectrophotometer at 570 nm. Cell viability was calculated by taking the ratio of the absorbance values of the drugged well over those of the negative control. $IC_{50}$ values could not be calculated accurately for some compounds as they exhibited only limited antiproliferative activity except at the highest concentrations.

#### E. GUV Rupture Fluorescence Imaging Assay Procedure:

To investigate the mechanism of cell potency of 5-fluorouracil and compounds **1** to **10**, we prepared giant unilamellar vesicles (GUVs) as model lipid membrane test systems. The GUVs were assembled based on literature procedures using a 1 mg/mL mixture of DOPC:RhodamineDPPE (99.5:0.5 mol %) [15]. To prepare the samples for imaging we passivated the wells of a 96-well glass bottom plate (Whatman 96-Well Glass Bottom Plate) with 1 mg/mL casein solution in 1X PBS. After a 1 hour incubation, we added 85 µL of 95 mM glucose to each of the wells and then added 2 µL of the GUV solution in 100 mM sucrose. After allowing the GUVs to sediment for 1 hour resting on the surface of the glass, we collected images of the GUVs before addition of the compounds using a Zeiss Axiovert 200M fluorescence microscope with a Zeiss A-plan 10x/0.25 Ph1 M27 objective and a Zeiss FLUAR 5x/0.25 M27 objective to excite and collect the fluorescence signal from the red-labeled GUVs at 555 nm. We then added 2 µL of 5-fluorouracil and compounds **1** to **10** at concentrations of 100 mM, 50 mM, 10 mM, 5 mM, 1 mM, 0.5 mM, 0.1 mM, 0.05 mM, 0.01 mM, 5 µM, 1 µM, 0.5 µM and observed the GUV rupturing events. A negative control was established with 0.0225 % v/v DMSO. Each sample was repeated in triplicate. Following a 15 minute incubation, we collected representative images of the GUVs in each well to characterize membrane disruption. The percent of GUV patches was determined by counting the number of patches in a 650 µm × 650 µm region and dividing by the total number of GUVs before rupture.

#### F. Chemical Synthesis:

*Preparation of the Benzyl urethane analog of Carmofur:*

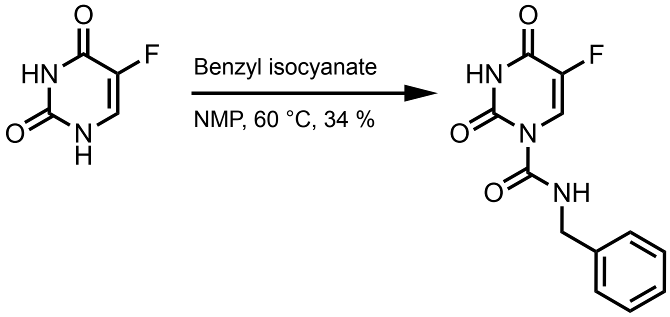


2

Following the general procedure yielded a white solid (34 % yield).

MW: 263.228 g/mol

**Chemicals:**
5-Fluorouracil (AK Scientific, 95 %): used without further purification
N-methylpyrrolidinone (Sigma Aldrich, 99.5 %, SureSeal bottle): used without further purification
Benzyl isocyanate (AK Scientific, 95 %): used without further purification

**Procedure:**
To a 50 mL oven-dried three-neck flask fitted with a water-jacketed condenser and a Teflon stir bar was added 5-fluorouracil (200.0 mg, 1.54 mmol, 1.0 eq) under nitrogen, and then anhydrous N-methylpyrrolidinone (12.00 mL, 0.13 M) was added via syringe followed by benzyl isocyanate (0.284 mL, 2.31 mmol, 1.5 eq). The reaction was heated to 60 °C in a silicone oil bath and tracked by aliquot $^{19}$F NMR spectroscopy until reaction completion after 30 minutes. The

reaction mixture was then quenched with 1.0 M brine (50 mL), aq. 1.0 M HCl (50 mL), and extracted with ethyl acetate (2x50 mL). The combined organic layers were dried over anhydrous magnesium sulfate, filtered, and concentrated *in vacuo*. The resulting residue was directly loaded onto a silica column (2.3 cm dia., 10 cm stack) and purified by flash chromatography (0 % → 30 % EtOAc in hexanes) to afford the title compound as a white powder (138 mg, 34 % yield).

**Characterization Data for Compound 2:**

**TLC** $R_f$ = 0.72 (60 % EtOAc/Hexanes), UV active

**GC-MS:** Calculated for $[C_{12}H_{10}FN_3O_3]^+$ $[M]^+$: 263.228; found: 263.222

**FT-IR (ATR, $cm^{-1}$):** 3299, 3089, 2866, 2799, 2355, 1773, 1743, 1715, 1693, 1651, 1607, 1585, 1520, 1496, 1464, 1455, 1335, 1281, 1264, 1233, 1206, 1113, 1079, 1048, 1028, 1001, 910, 834, 757, 730, 694, 667, 622, 614, 598, 561

**$^1$H NMR** (500 MHz, Acetone-*d6*) δ 10.92 (s, 1H), 9.57 (s, 1H), 8.42 (d, J = 7.6 Hz, 1H), 7.39 – 7.23 (m, 5H), 4.57 (d, J = 3.9 Hz, 2H).

**$^{19}$F NMR** (57 MHz, Acetone-*d6*) δ -166.31 (d, *J* = 7.6 Hz).

**$^{13}$C NMR** (126 MHz, Acetone-*d6*) δ 150.64, 150.04, 142.19, 140.32, 138.33, 128.57, 127.63, 127.40, 122.68, 122.38, 44.48, 44.36.

**¹H NMR for Compound 2** (500 MHz, Acetone-*d6*)**:**

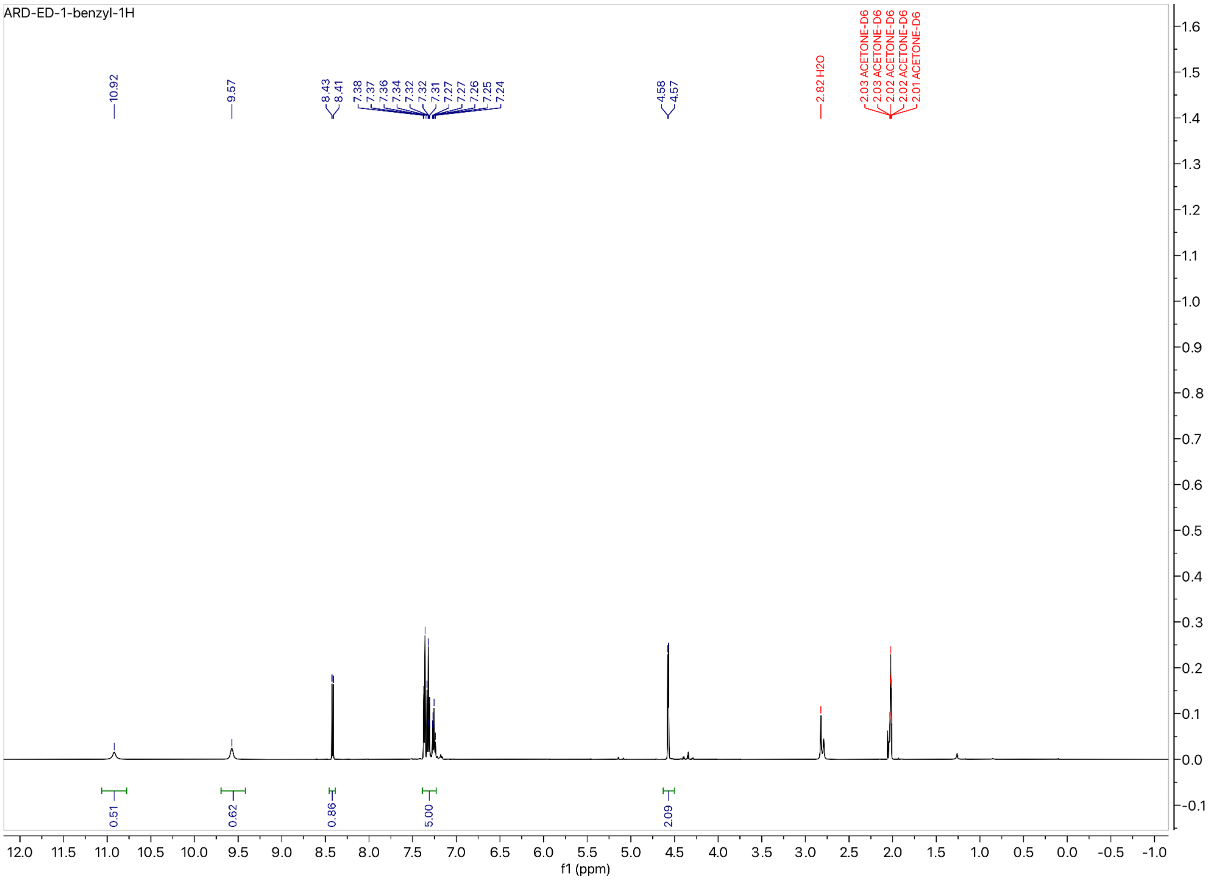

**[19]F NMR for Compound 2** (57 MHz, Acetone-*d*6):

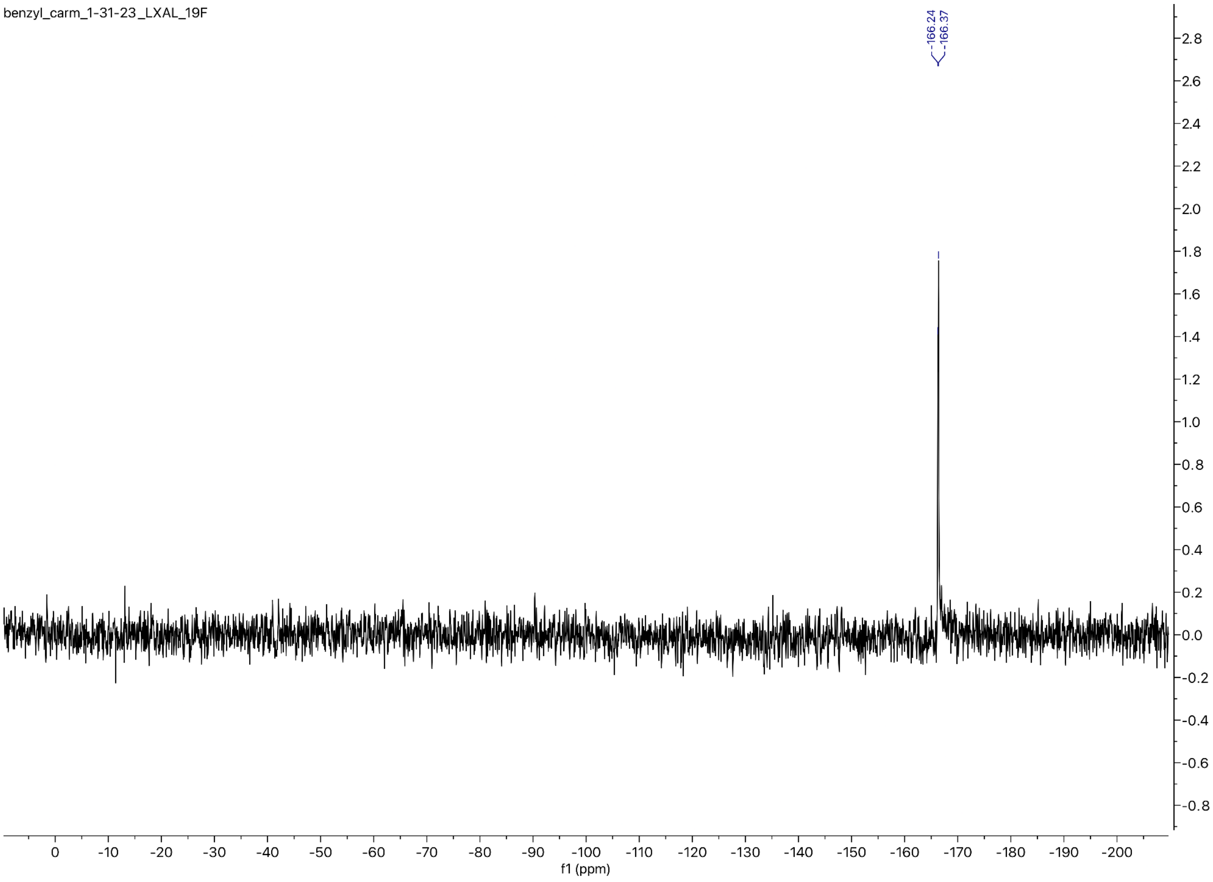

**$^{13}$C NMR for Compound 2** (101 MHz, Acetone-*d6*):

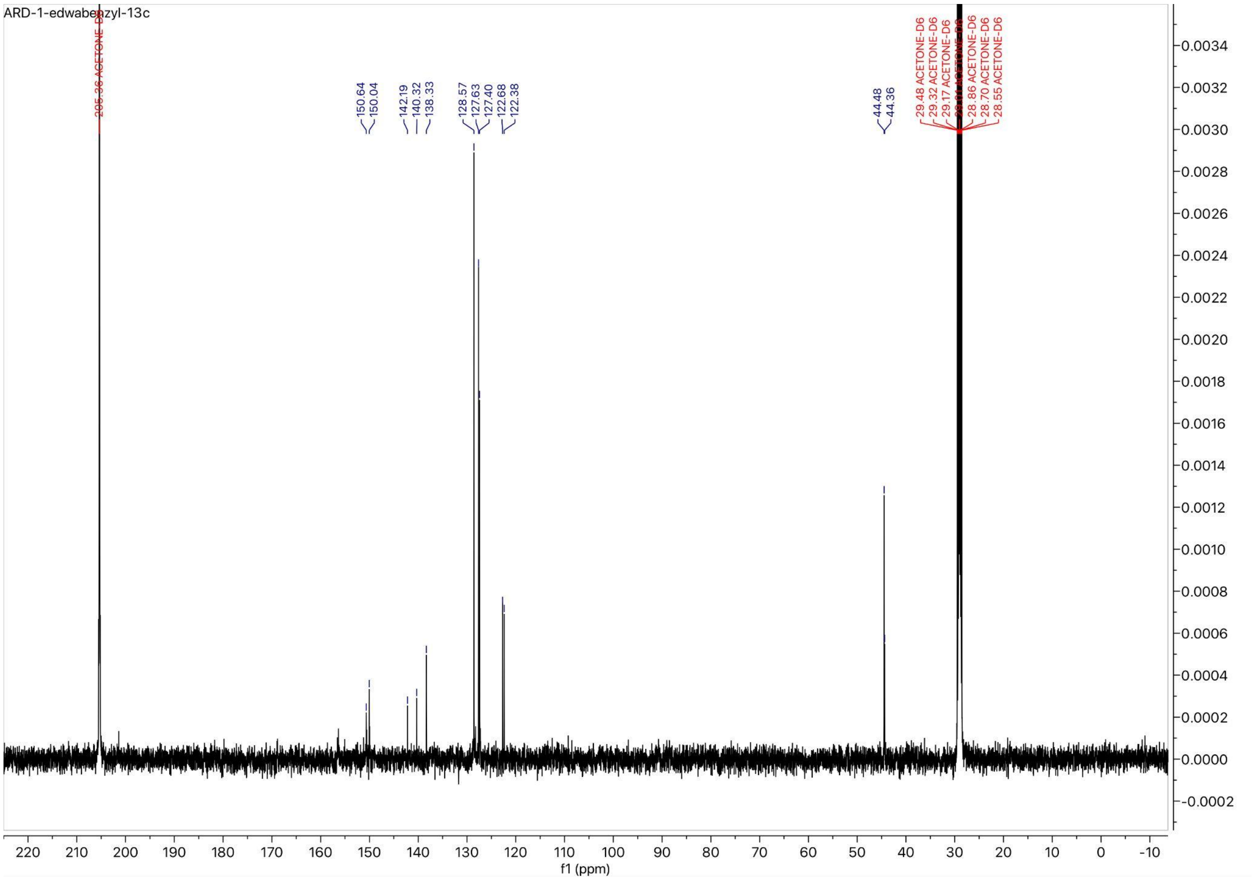

*Preparation of the Trans-4-methylcyclohexyl urethane analog of Carmofur:*

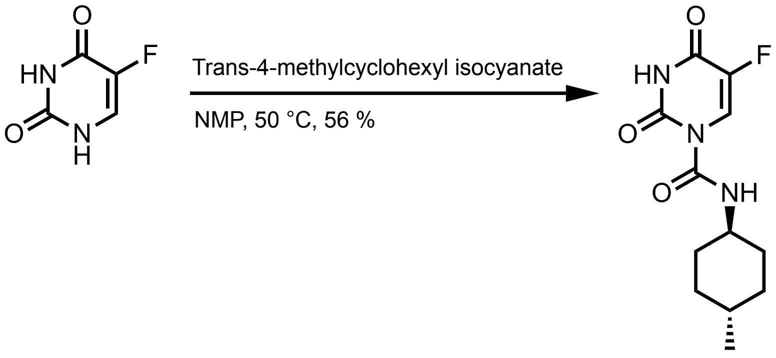


3

Following the general procedure yielded a white solid (56 % yield)

MW: 269.276 g/mol

**Chemicals:**
5-Fluorouracil (AK Scientific, 95 %): used without further purification
N-methylpyrrolidinone (Sigma Aldrich, 99.5 %, SureSeal bottle): used without further purification
Trans-4-methylcyclohexyl isocyanate (AK Scientific, 95 %): used without further purification

**Procedure:**
To a 50 mL oven-dried three-neck flask fitted with a water-jacketed condenser and a Teflon stir bar was added 5-fluorouracil (200.0 mg, 1.54 mmol, 1.0 eq) under nitrogen, and then anhydrous N-methylpyrrolidinone (12.00 mL, 0.13 M) was added via syringe followed by trans-4-methylcyclohexyl isocyanate (0.429 mL, 3.08 mmol, 2.0 eq). The reaction was heated to 50 °C in a silicone oil bath and tracked by aliquot $^{19}$F NMR spectroscopy until reaction completion after 150 minutes. The reaction mixture was then quenched with 1.0 M brine (50 mL), aq. 1.0 M HCl (50 mL), and extracted with ethyl acetate (2x50 mL). The combined organic layers were dried over anhydrous magnesium sulfate, filtered, and concentrated *in vacuo*. The resulting residue was directly loaded onto a silica column (2.3 cm dia., 10 cm stack) and purified by flash chromatography (0 % → 35 % EtOAc in hexanes) to afford the title compound as a white powder (230. mg, 56 % yield).

**Characterization Data for Compound 3:**

**TLC** $R_f$ = 0.60 (60 % EtOAc/Hexanes), UV active

**GC-MS:** Calculated for $[C_{12}H_{16}FN_3O_3]^+$ $[M]^+$: 269.276; found: 269.278

**FT-IR (ATR, cm$^{-1}$):** 3326, 3286, 3079, 3037, 2952, 2927, 2904, 2865, 2818, 1755, 1724, 1698, 1683, 1652, 1624, 1576, 1539, 1507, 1439, 1420, 1377, 1362, 1339, 1311, 1267, 1236, 1213, 1197, 1156, 1106, 1079, 1045, 1005, 968, 953, 901, 883, 857, 795, 766, 749, 736, 705, 639, 621, 595, 569, 562

**$^{1}$H NMR** (400 MHz, DMSO-*d*6) δ 11.48 (s, 1H), 9.03 (d, *J* = 7.3 Hz, 1H), 8.31 (d, *J* = 7.4 Hz, 1H), 7.69 (d, *J* = 6.0 Hz, 1H), 3.39 – 3.28 (m, 9H), 1.86 (d, *J* = 12.4 Hz, 2H), 1.64 (d, *J* = 13.3 Hz, 2H), 1.33 – 1.17 (m, 3H), 0.96 (q, *J* = 10.2 Hz, 2H), 0.82 (t, *J* = 6.5 Hz, 3H).

**$^{19}$F NMR** (57 MHz, Chloroform-*d*) δ -162.65 (d, *J* = 6.2 Hz).

**$^{13}$C NMR** (101 MHz, DMSO-*d*6) δ 150.97, 149.50, 149.07, 142.57, 126.55, 123.19, 122.82, 50.48, 33.60, 32.34, 31.56, 22.38.

**$^1$H NMR for Compound 3** (400 MHz, DMSO-*d*6):

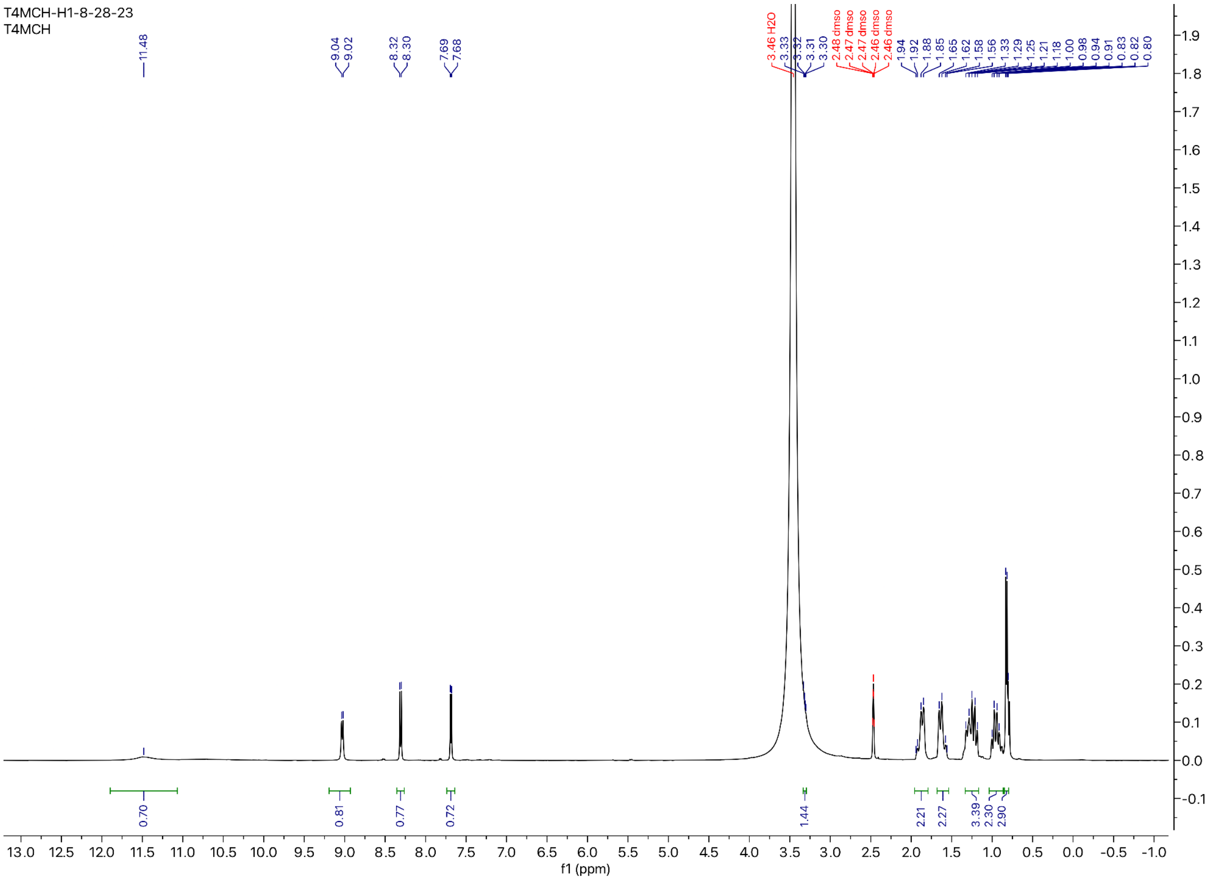

**$^{19}$F NMR for Compound 3** (57 MHz, Chloroform-*d*)

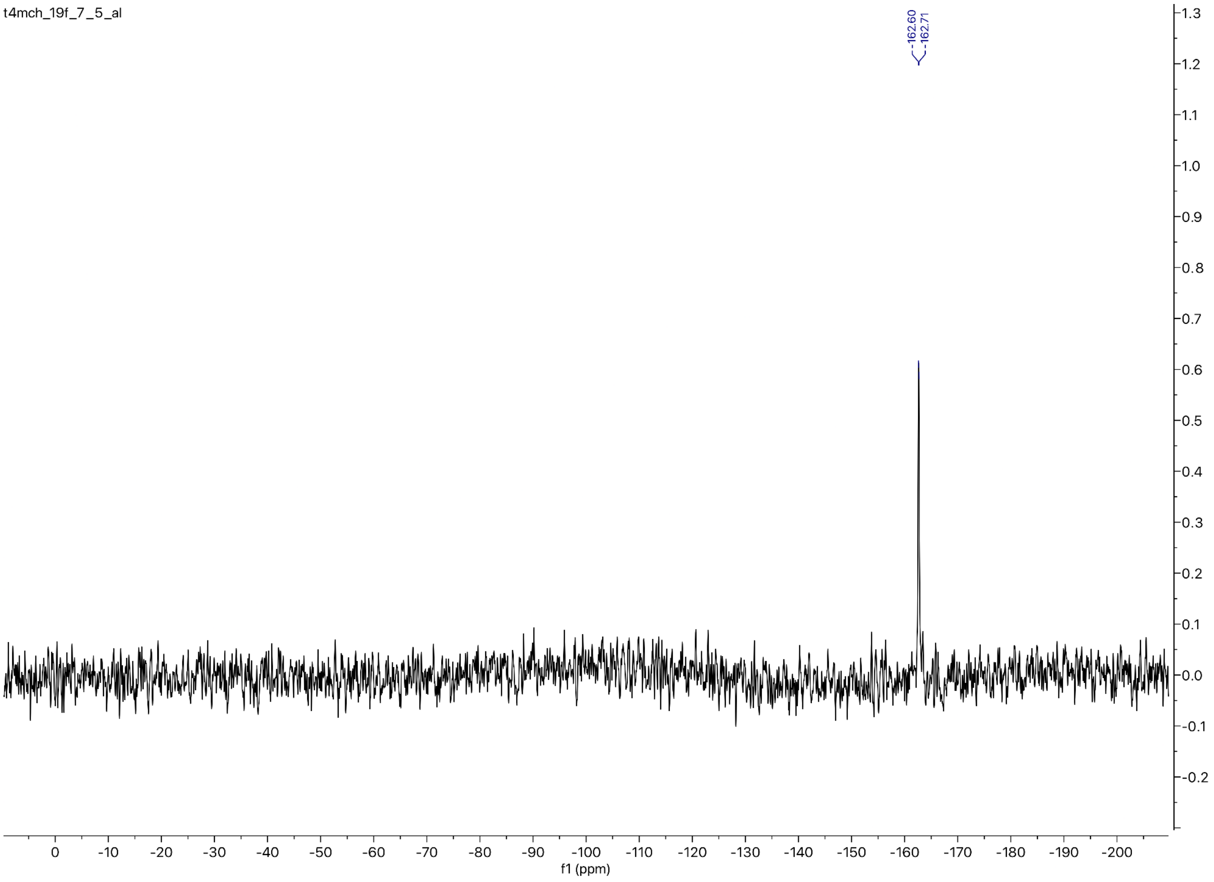

**$^{13}$C NMR for Compound 3** (101 MHz, DMSO-*d6*)

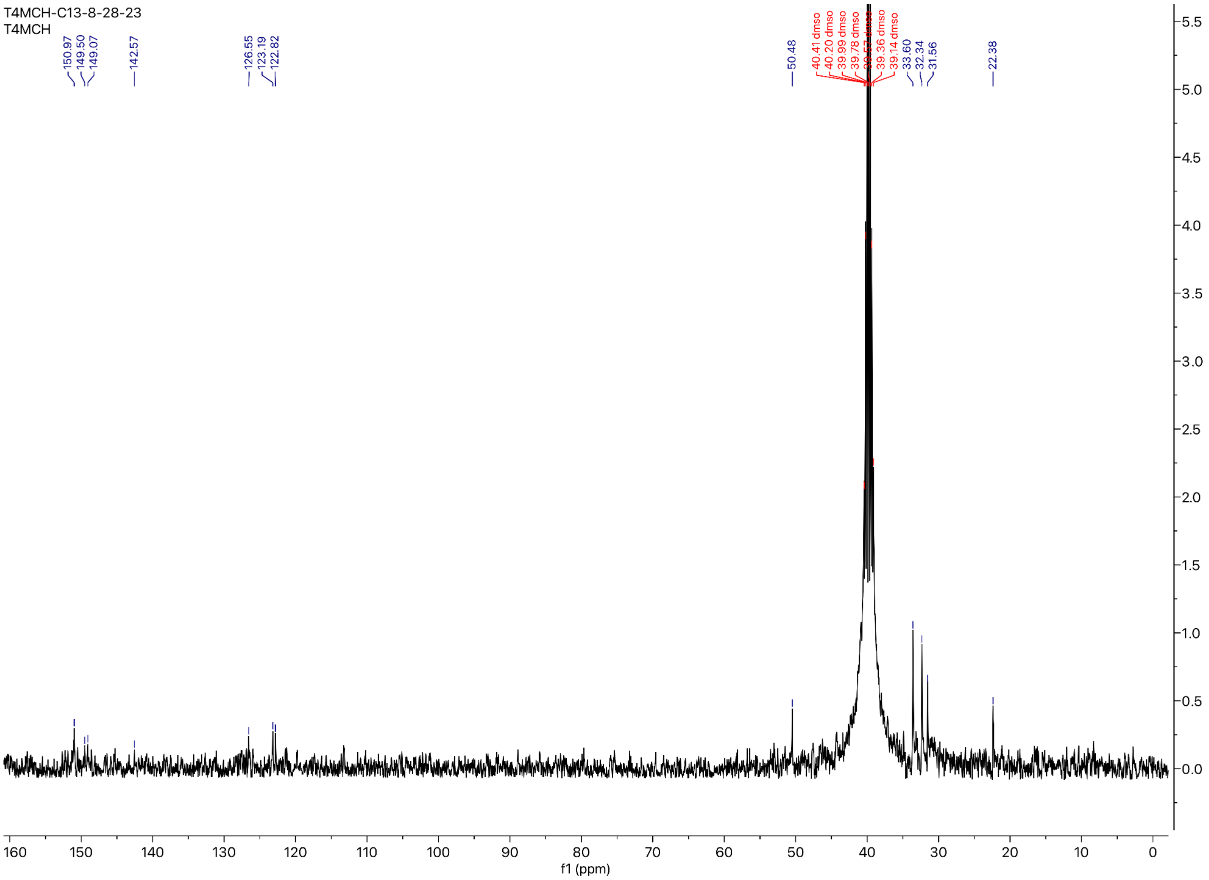

#### *Preparation of the 2-Chloroethyl urethane analog of Carmofur:*

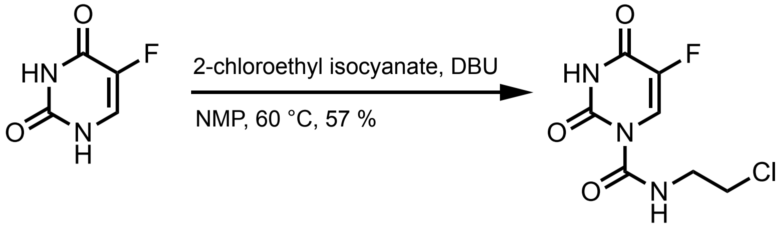


**4**

Following the general procedure yielded a white solid (57 % yield)

MW: 235.599 g/mol

**Chemicals:**
5-Fluorouracil (AK Scientific, 95 %): used without further purification
N-methylpyrrolidinone (Sigma Aldrich, 99.5 %, SureSeal bottle): used without further purification
1,8-Diazabicyclo[5.4.0]undec-7-ene (AK Scientific, 99 %): used without further purification
2-Chloroethyl isocyanate (AK Scientific, 95 %): used without further purification

**Procedure:**
To a 50 mL oven-dried three-neck flask fitted with a water-jacketed condenser and a Teflon stir bar was added 5-fluorouracil (400.0 mg, 3.08 mmol, 1.0 eq) under nitrogen, and then anhydrous N-methylpyrrolidinone (10.00 mL, 0.31 M) was added via syringe followed by 1,8-diazabicyclo[5.4.0]undec-7-ene (0.069 mL, 0.46 mmol, 0.15 eq) and 2-chloroethyl isocyanate (0.200 mL, 2.34 mmol, 0.76 eq). The reaction was heated to 60 °C in a silicone oil bath and tracked by aliquot $^{19}$F NMR spectroscopy until reaction completion after 120 minutes. The reaction mixture was then quenched with 1.0 M brine (50 mL) and extracted with ethyl acetate (2x50 mL). The combined organic layers were dried over anhydrous magnesium sulfate, filtered, and concentrated *in vacuo*. The resulting residue was directly loaded onto a silica column (2.3 cm dia., 10 cm stack) and purified by flash chromatography (10 % → 50 % EtOAc in hexanes) to afford the title compound as a white powder (410. mg, 57 % yield).

**Characterization Data for Compound 4:**

**TLC $R_f$**: 0.60 (30 % EtOAc/Hexanes), UV Active

**GC-MS:** Calculated for $[C_7H_7ClFN_3O_3]^+$ $[M]^+$: 235.599; found: 235.607

**FT-IR (ATR, $cm^{-1}$)**: 3274, 3089, 2929, 2836, 1722, 1691, 1667, 1515, 1456, 1435, 1334, 1267, 1247, 1222, 1205, 1188, 1145, 1101, 1074, 1061, 1036, 943, 883, 872, 846, 786, 758, 742, 655, 609, 581, 572, 564, 553

**$^1$H NMR** (400 MHz, Acetone-*d6*) δ 10.84 (s, 1H), 9.35 (s, 1H), 8.30 (d, *J* = 7.4 Hz, 1H), 3.68 – 3.60 (m, 4H).

**$^{19}$F NMR** (57 MHz, Chloroform-*d*) δ -161.94 (d, *J* = 4.8 Hz).

**$^{13}$C NMR** (101 MHz, Acetone-*d6*) δ 150.49, 150.02, 142.40, 140.05, 122.43, 122.05, 42.71.

**$^{1}$H NMR for Compound 4** (400 MHz, Acetone-*d6*):

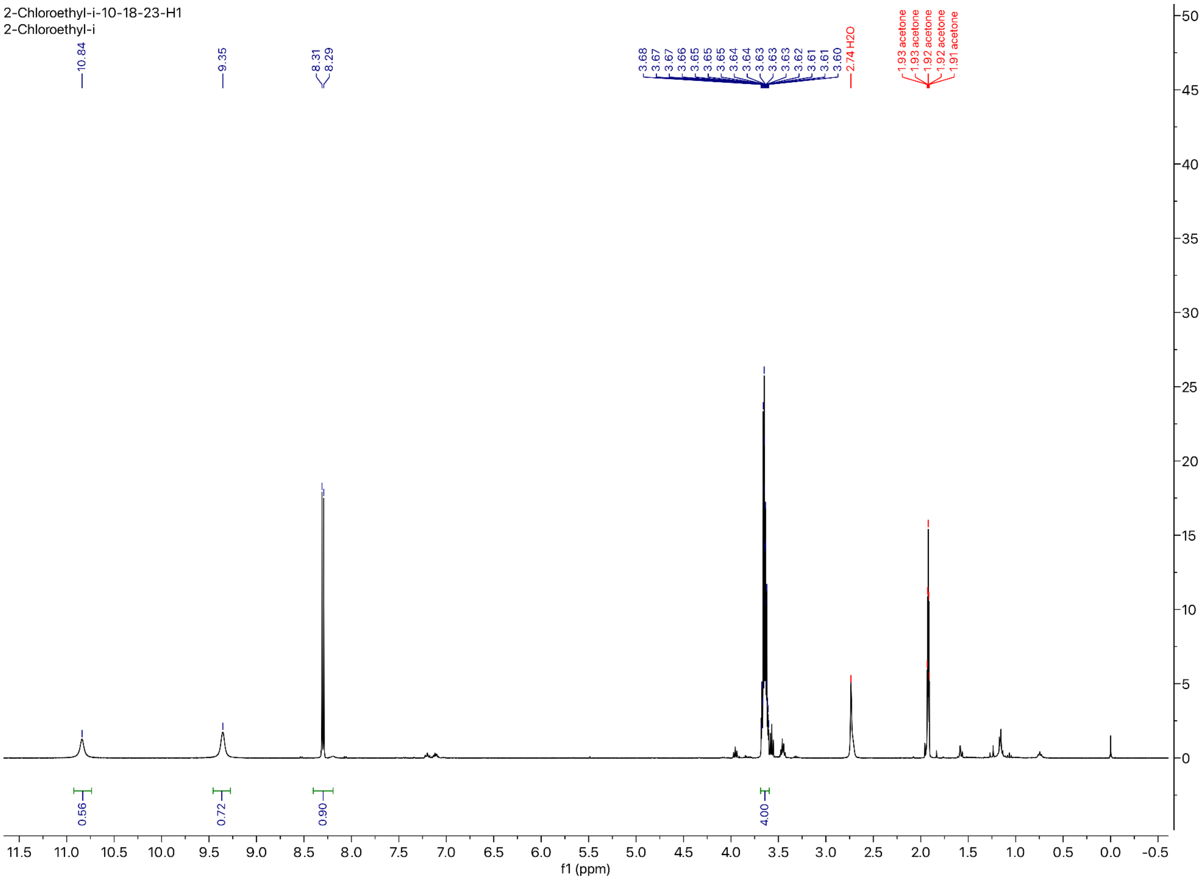

**$^{19}$F NMR for Compound 4** (57 MHz, Chloroform-*d*)**:**

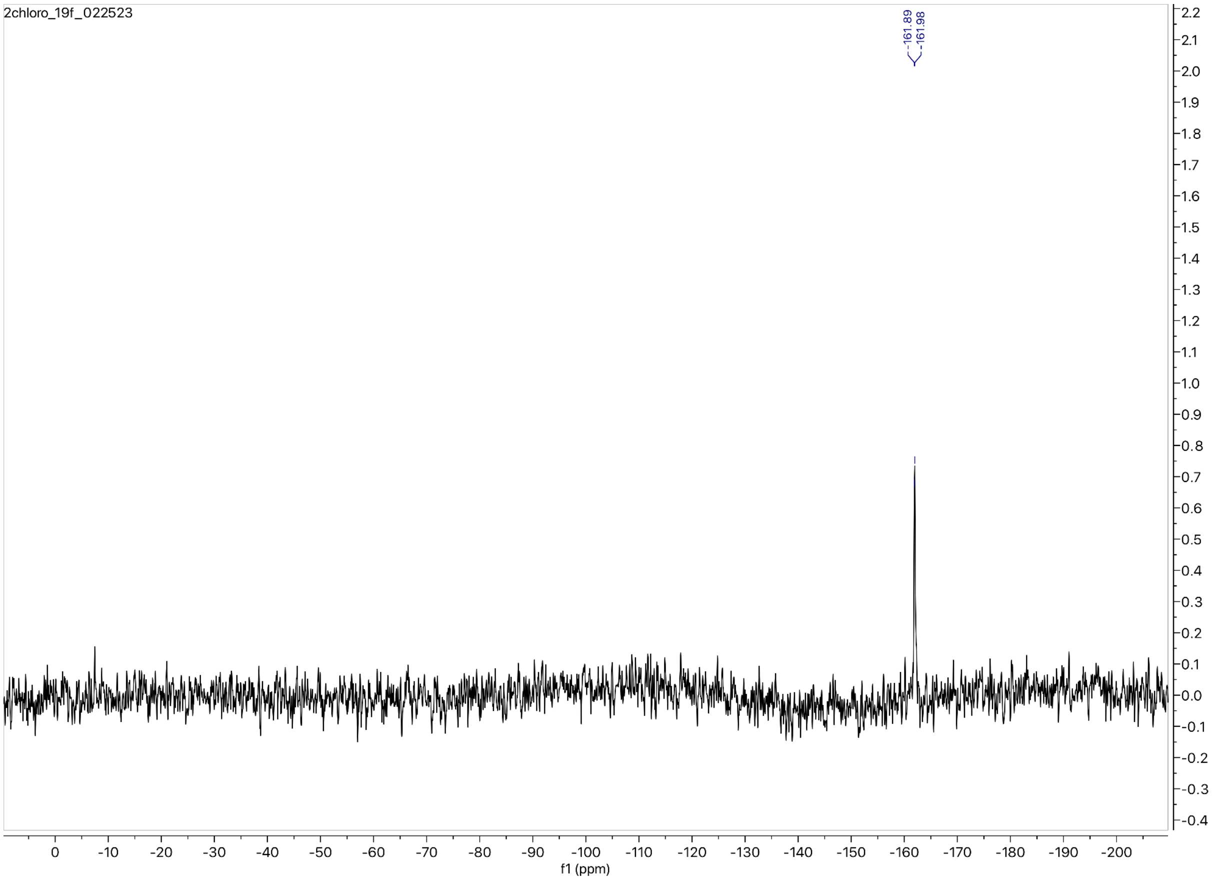

**$^{13}$C NMR for Compound 4** (101 MHz, Acetone-*d6*):

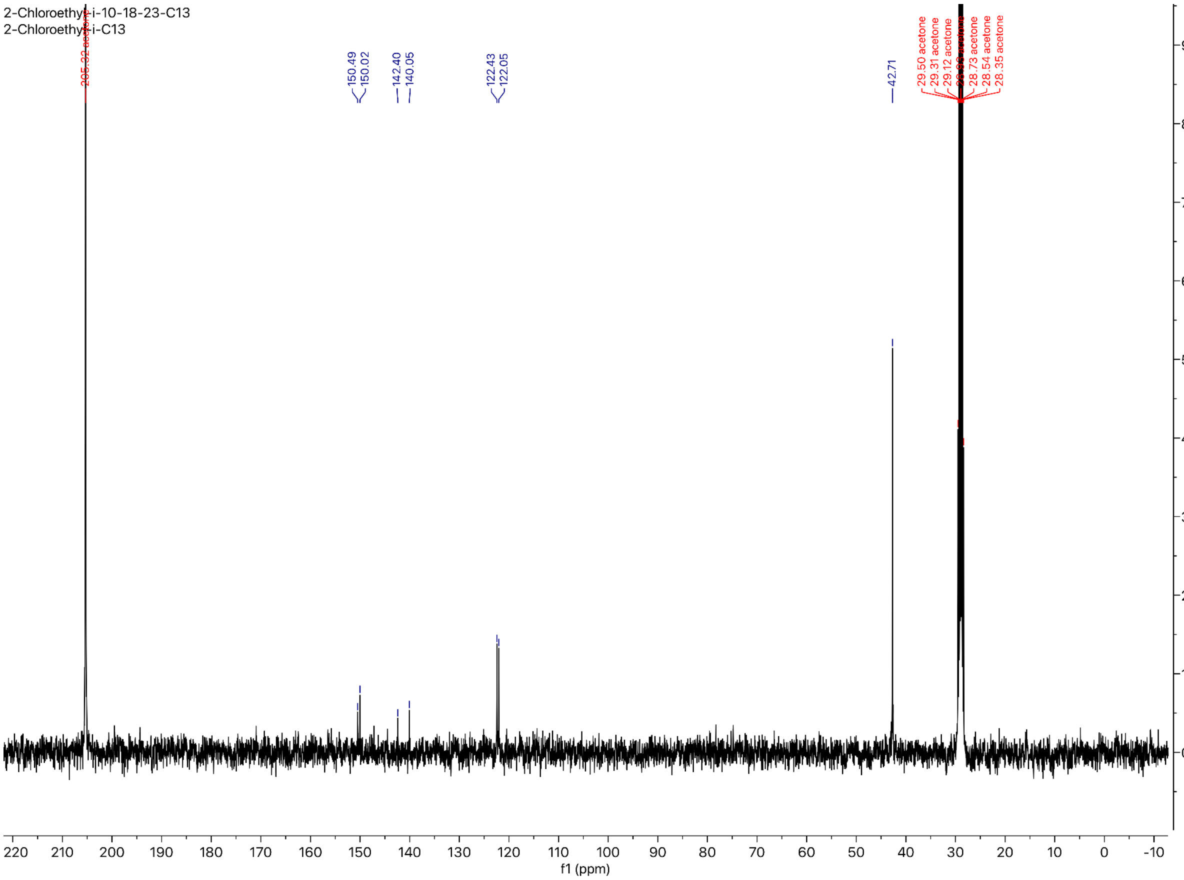

#### *Preparation of the Dodecyl urethane analog of Carmofur:*

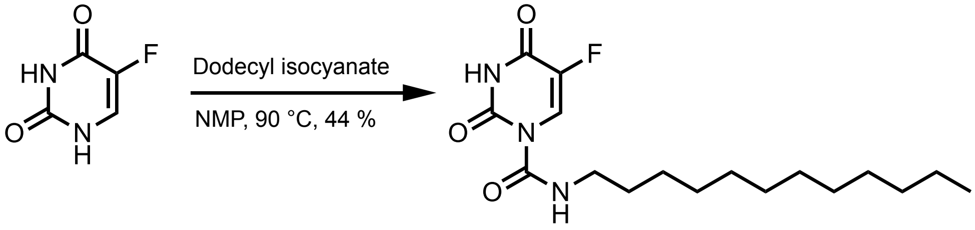


**5**

Following the general procedure yielded a white solid (44 % yield)

MW: 341.427 g/mol

**Chemicals:**

5-Fluorouracil (AK Scientific, 95 %): used without further purification
N-methylpyrrolidinone (Sigma Aldrich, 99.5 %, SureSeal bottle): used without further purification
Dodecyl isocyanate (AK Scientific, 95 %): used without further purification

**Procedure:**

To a 50 mL oven-dried three-neck flask fitted with a water-jacketed condenser and a Teflon stir bar was added 5-fluorouracil (200.0 mg, 1.54 mmol, 1.0 eq) under nitrogen, and then anhydrous N-methylpyrrolidinone (12.00 mL, 0.13 M) was added via syringe followed by dodecyl isocyanate (0.557 mL, 2.31 mmol, 1.5 eq). The reaction was heated to 90 °C in a silicone oil bath and tracked by aliquot $^{19}F$ NMR spectroscopy until reaction completion after 150 minutes. The reaction mixture was then quenched with 1.0 M brine (50 mL), aq. 1.0 M HCl (50 mL), and extracted with ethyl acetate (2x50 mL). The combined organic layers were dried over anhydrous magnesium sulfate, filtered, and concentrated *in vacuo*. The resulting residue was directly loaded onto a silica column (2.3 cm dia., 10 cm stack) and purified by flash chromatography (0 % → 30 % EtOAc in hexanes) to afford the title compound as a white powder (231 mg, 44 % yield).

**Characterization Data for Compound 5:**

**TLC $R_f$**: 0.90 (60 % EtOAc/Hex), UV Active

**GC-MS:** Calculated for $[C_{17}H_{28}FN_3O_3]^+$ $[M]^+$: 341.427; found: 341.411

**FT-IR (ATR, $cm^{-1}$):** 3335, 3287, 3099, 3053, 2959, 2919, 2847, 2360, 2344, 1741, 1729, 1683, 1669, 1636, 1612, 1574, 1537, 1462, 1431, 1377, 1362, 1331, 1273, 1239, 1220, 1195, 1168, 1099, 1043, 1026, 932, 863, 838, 805, 778, 760, 751, 742, 724, 668, 630, 615, 600, 581, 570, 564, 553

**$^1$H NMR** (400 MHz, Chloroform-*d*) δ 9.03 (t, *J* = 4.9 Hz, 1H), 8.49 (d, *J* = 6.7 Hz, 1H), 3.39 (q, *J* = 7.4 Hz, 2H), 1.60 (p, *J* = 7.2 Hz, 2H), 1.42 – 1.17 (m, 22H), 0.88 (t, *J* = 6.9 Hz, 3H).

**$^{19}$F NMR** (57 MHz, Chloroform-*d*) δ -162.53 (d, *J* = 7.6 Hz).

**$^{13}$C NMR** (101 MHz, Chloroform-*d*) δ 156.96, 156.68, 150.07, 149.09, 141.99, 139.58, 123.47, 123.10, 41.50, 31.93, 29.65, 29.64, 29.58, 29.51, 29.36, 29.21, 29.15, 26.82, 22.70, 14.12.

**[1]H NMR for Compound 5** (400 MHz, Chloroform-*d*):

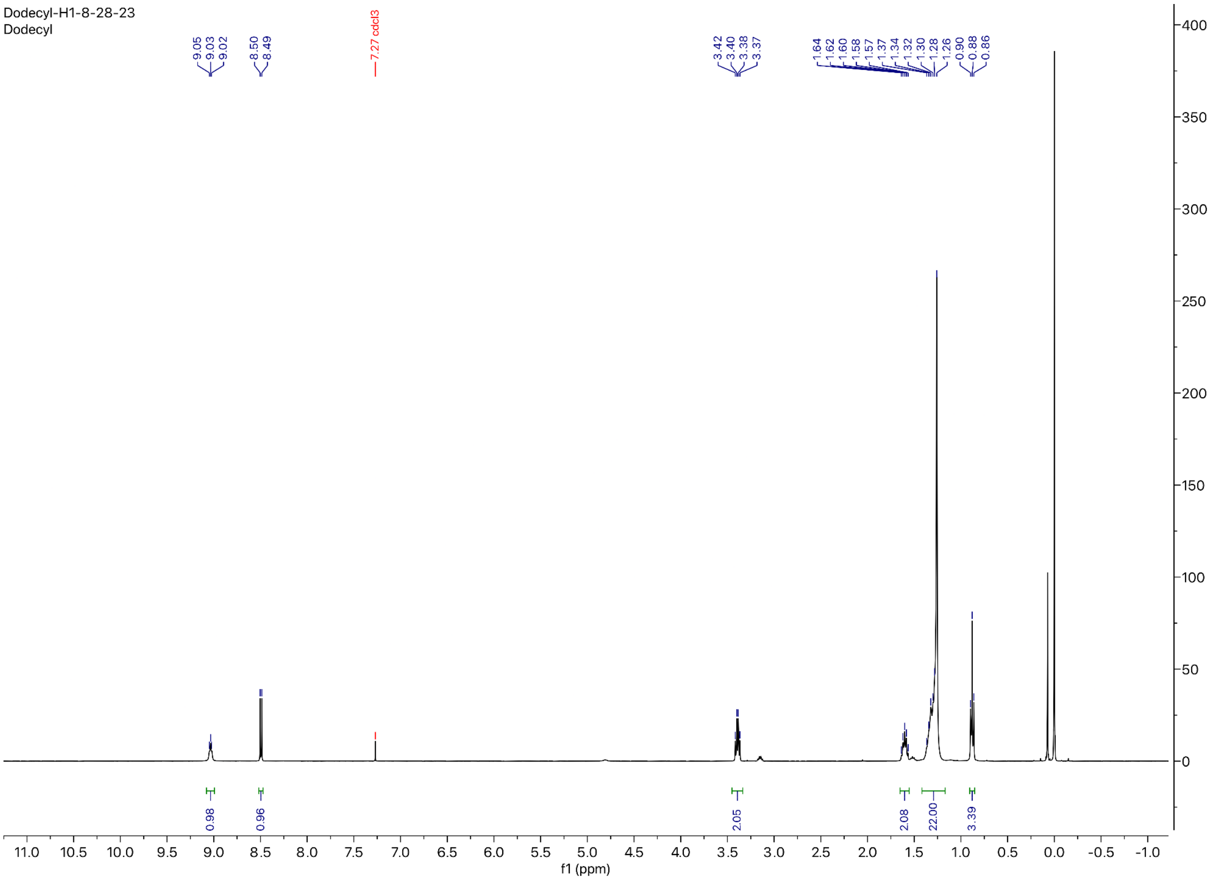

**$^{19}$F NMR for Compound 5** (57 MHz, Chloroform-*d*)

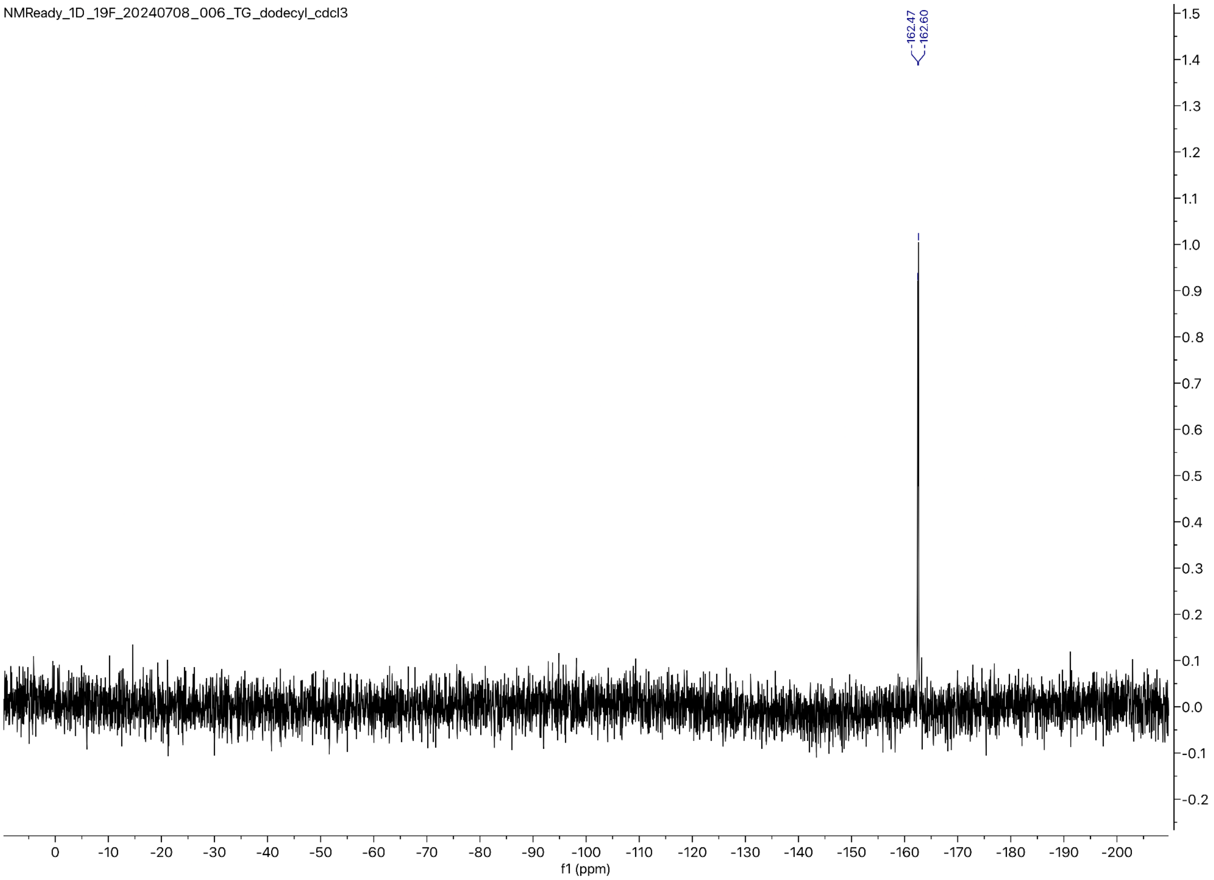

**$^{13}$C NMR for Compound 5** (101 MHz, Chloroform-*d*):

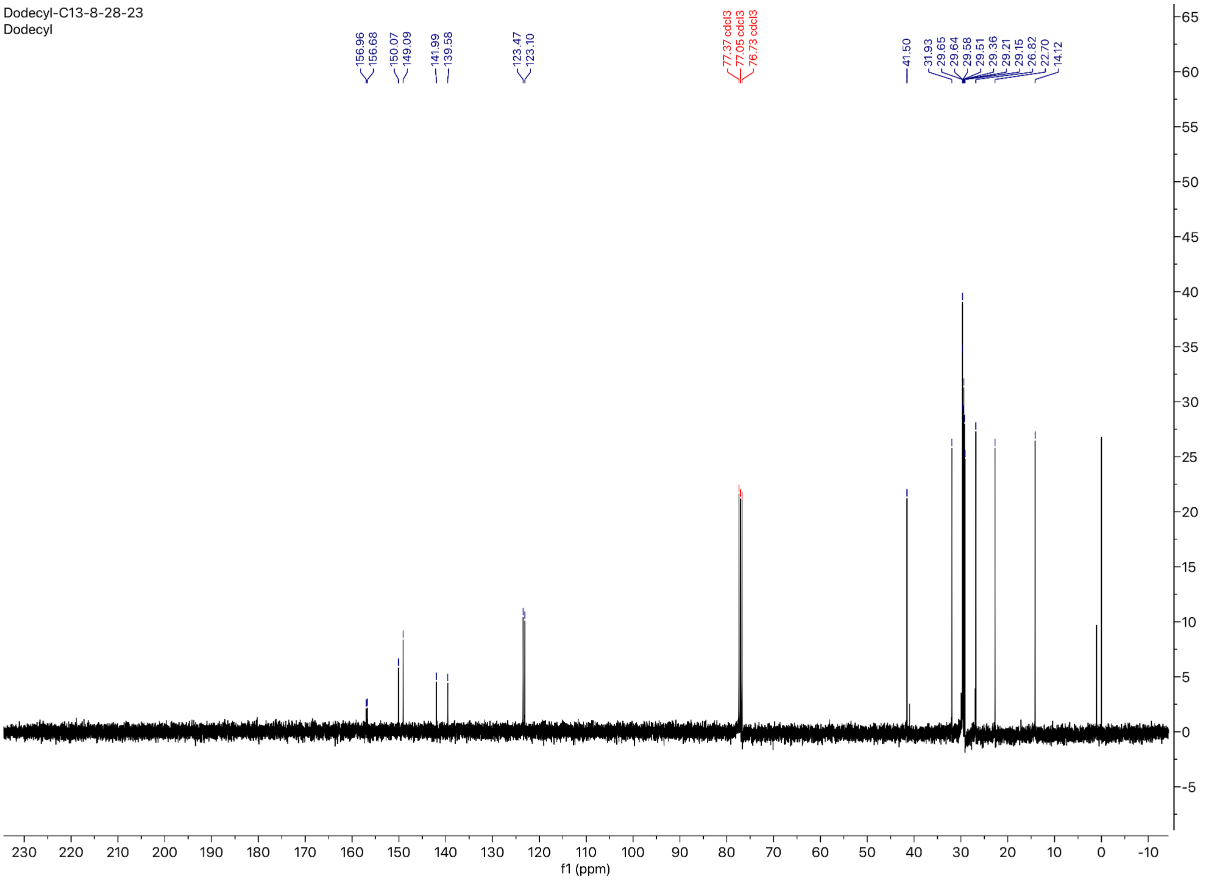

*Preparation of the Octadecyl urethane analog of Carmofur:*

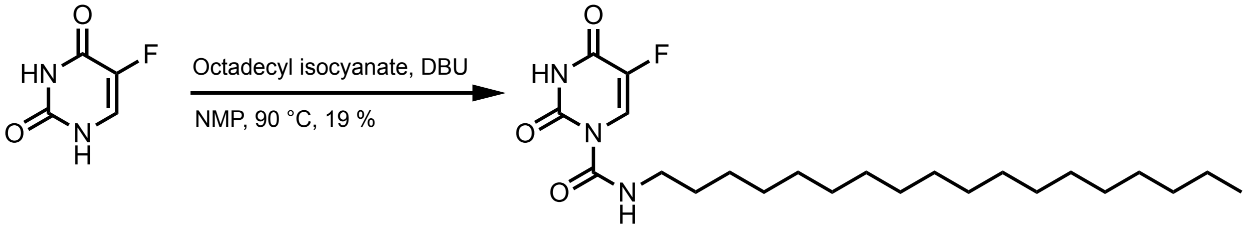


6

Following the general procedure yielded a white solid (19 % yield)

MW: 425.589 g/mol

**Chemicals:**
5-Fluorouracil (AK Scientific, 95 %): used without further purification
N-methylpyrrolidinone (Sigma Aldrich, 99.5 %, SureSeal bottle): used without further purification
1,8-Diazabicyclo[5.4.0]undec-7-ene (AK Scientific, 99 %): used without further purification
Octadecyl isocyanate (AK Scientific, >80 %): used without further purification

**Procedure:**
To a 50 mL oven-dried three-neck flask fitted with a water-jacketed condenser and a Teflon stir bar was added 5-fluorouracil (100.0 mg, 0.77 mmol, 1.0 eq) under nitrogen, and then anhydrous N-methylpyrrolidinone (3.00 mL, 0.26 M) was added via syringe followed by 1,8-diazabicyclo[5.4.0]undec-7-ene (0.017 mL, 0.12 mmol, 0.15 eq) and octadecyl isocyanate (0.402 mL, 1.15 mmol, 1.5 eq). The reaction was heated to 90 °C in a silicone oil bath and tracked by aliquot $^{19}$F NMR spectroscopy until reaction completion after 150 minutes. The reaction mixture was then quenched with 1.0 M brine (50 mL), aq. 1.0 M HCl (50 mL), and extracted with ethyl acetate (2x50 mL). The combined organic layers were dried over anhydrous magnesium sulfate, filtered, and concentrated *in vacuo*. The resulting residue was directly loaded onto a silica column (2.3 cm dia., 10 cm stack) and purified by flash chromatography (0 % → 30 % EtOAc in hexanes) to afford the title compound as a white powder (63 mg, 19 % yield).

**Characterization Data for Compound 6:**

**TLC $R_f$**: 0.90 (50 % EtOAc/Hexanes), UV Active

**GC-MS:** Calculated for $[C_{23}H_{40}FN_3O_3]^+$ $[M]^+$: 425.589; found: 425.598

**FT-IR (ATR, cm$^{-1}$):** 2959, 2918, 2847, 2361, 2344, 1746, 1723, 1684, 1669, 1654, 1647, 1636, 1576, 1538, 1507, 1497, 1489, 1462, 1435, 1375, 1338, 1274, 1220, 1197, 1099, 1046, 1031, 1022, 932, 899, 864, 823, 801, 781, 760, 753, 744, 724, 634, 616, 599, 592, 580, 571, 564, 553

**$^1$H NMR** (400 MHz, Pyridine-*d*5) δ 9.61 (t, *J* = 5.6 Hz, 1H), 8.75 (s, 1H), 3.48 (q, *J* = 6.1 Hz, 2H), 1.62 (p, *J* = 6.9 Hz, 2H), 1.39 – 1.23 (m, 34H), 0.88 (t, *J* = 7.2 Hz, 3H).

**$^{19}$F NMR** (57 MHz, Pyridine-*d*5) δ -163.06 (d, *J* = 9.2 Hz).

**$^{13}$C NMR** (101 MHz, Pyridine-*d*5) δ 157.91, 157.64, 151.47, 150.09, 140.41, 138.06, 122.72, 122.58, 41.18, 31.89, 31.56, 29.76, 29.75, 29.71, 29.69, 29.64, 29.58, 29.43, 29.38, 29.29, 26.91, 22.70, 14.05.

**$^{1}$H NMR for Compound 6** (400 MHz, Pyridine-*d*5):

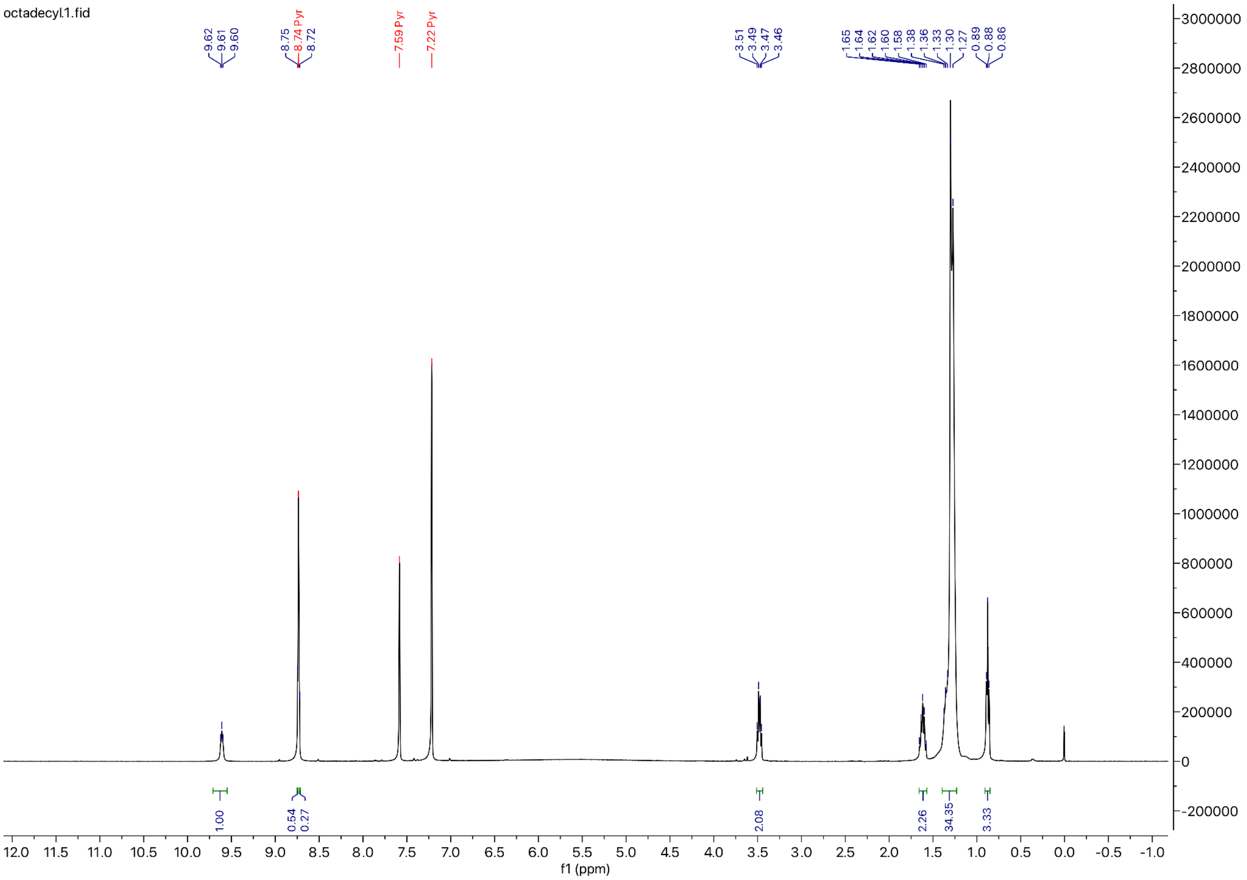

**$^{19}$F NMR for Compound 6** (57 MHz, Pyridine-*d*5)**:**

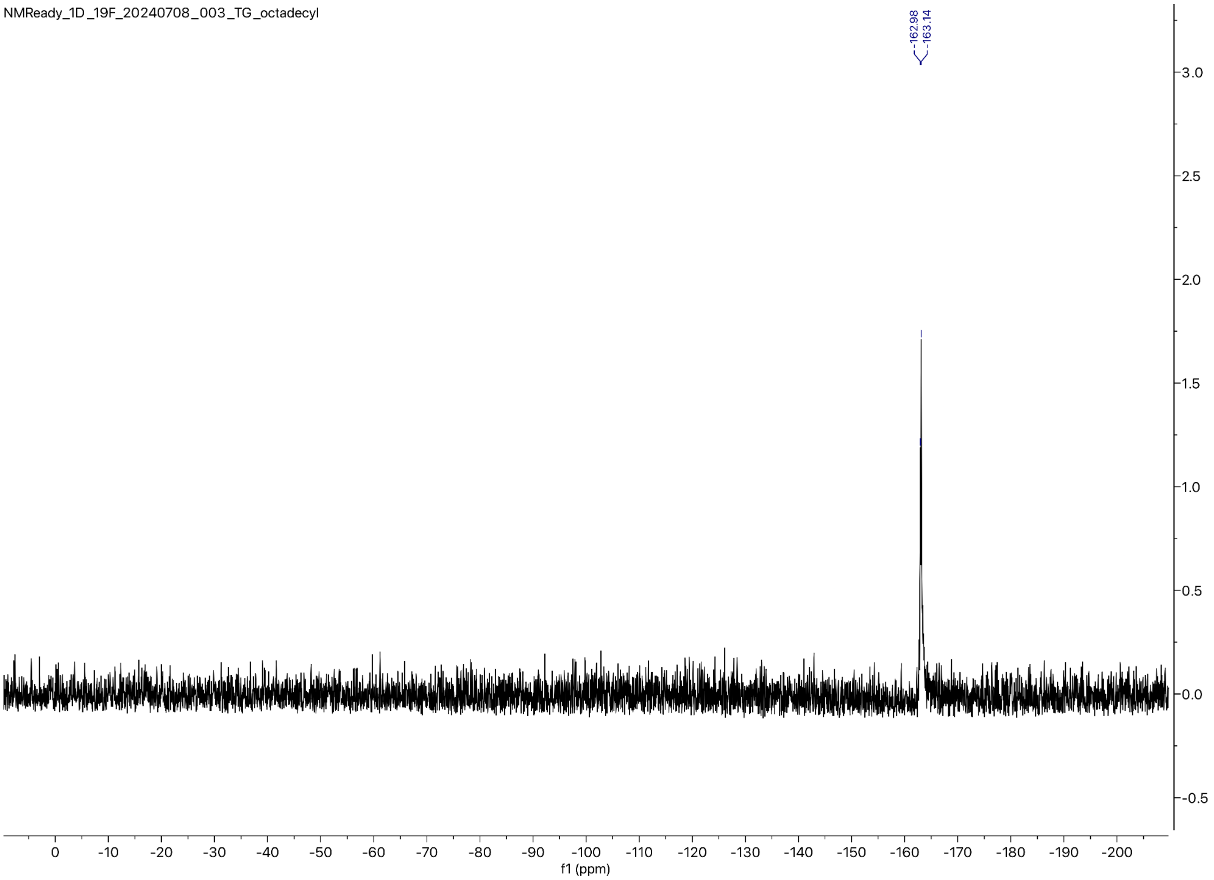

**$^{13}$C NMR for Compound 6** (101 MHz, Pyridine-*d5*):

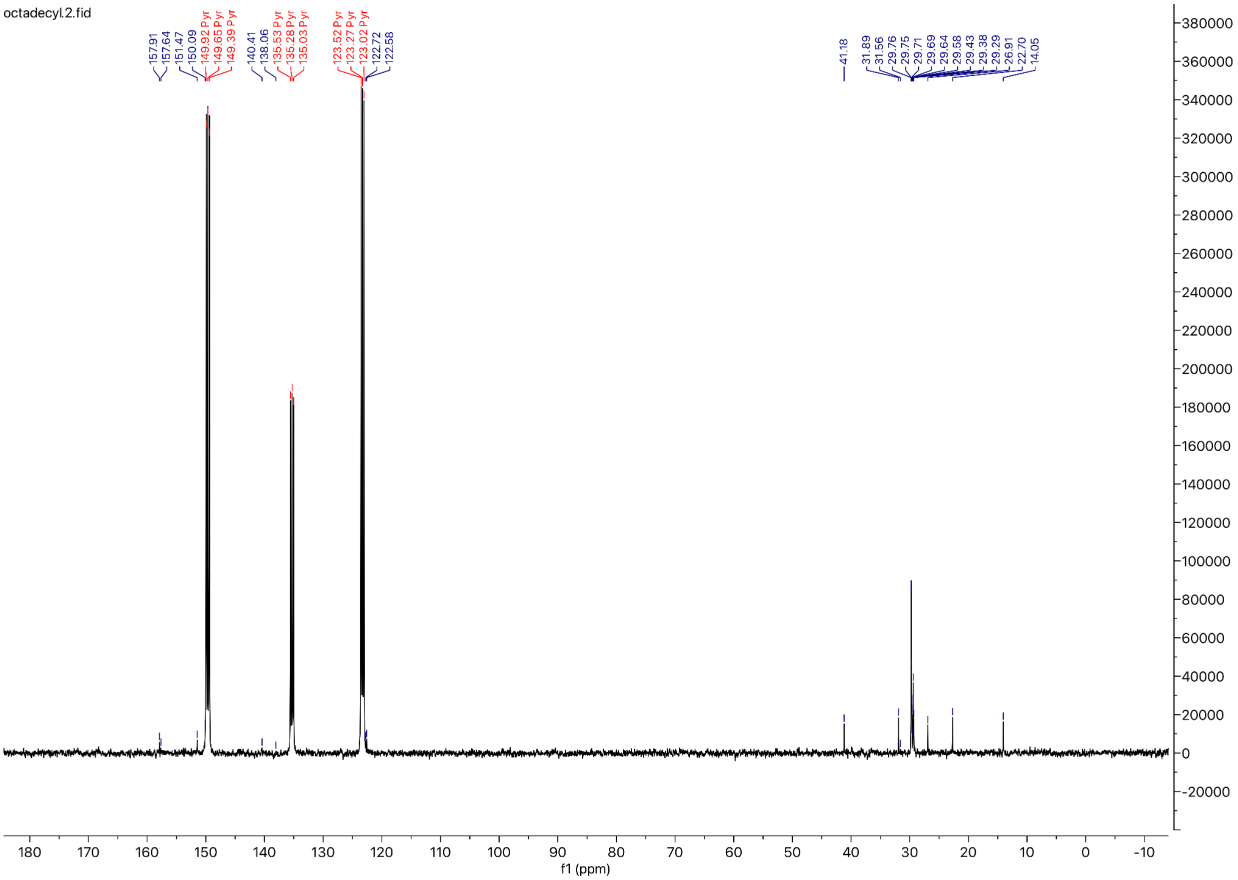

*Preparation of the Methoxyethyl urethane analog of Carmofur:*

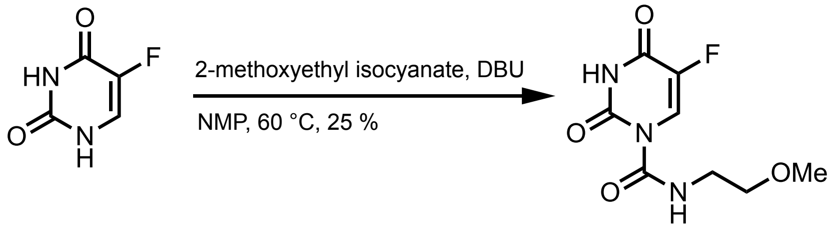


7

Following the general procedure yielded a white solid (25 %)

MW: 231.183 g/mol

**Chemicals:**

5-Fluorouracil (AK Scientific, 95 %): used without further purification
N-methylpyrrolidinone (Sigma Aldrich, 99.5 %, SureSeal bottle): used without further purification
1,8-Diazabicyclo[5.4.0]undec-7-ene (AK Scientific, 99 %): used without further purification
2-Methoxyethyl isocyanate (Oakwood Chemical, 97 %): used without further purification

**Procedure:**

To a 50 mL oven-dried three-neck flask fitted with a water-jacketed condenser and a Teflon stir bar was added 5-fluorouracil (300.0 mg, 2.31 mmol, 1.0 eq) under nitrogen, and then anhydrous N-methylpyrrolidinone (8.00 mL, 0.29 M) was added via syringe followed by 1,8-diazabicyclo[5.4.0]undec-7-ene (0.051 mL, 0.35 mmol, 0.15 eq) and 2-methoxyethyl isocyanate (0.360 mL, 3.46 mmol, 1.5 eq). The reaction was heated to 60 °C in a silicone oil bath and tracked by aliquot $^{19}F$ NMR spectroscopy until reaction completion after 120 minutes. The reaction mixture was then quenched with 1.0 M brine (50 mL), aq. 1.0 M HCl (50 mL), and extracted with ethyl acetate (2x50 mL). The combined organic layers were dried over anhydrous magnesium sulfate, filtered, and concentrated *in vacuo*. The resulting residue was directly loaded onto a silica column (2.3 cm dia., 10 cm stack) and purified by flash chromatography (0 % → 40 % EtOAc in hexanes) to afford the title compound as a white powder (57 mg, 25 % yield).

**Characterization Data for Compound 7**

**TLC $R_f$**: 0.60 (30 % EtOAc/Hexanes), UV Active

**HRMS:** Calculated for $[C_8H_{10}FN_3O_4]^+$ $[M]^+$: 231.1834; found: 231.2100

**FT-IR (ATR, $cm^{-1}$):** 3290, 3163, 3072, 2934, 2898, 2866, 2824, 2359, 2342, 1728, 1694, 1559, 1522, 1507, 1496, 1455, 1388, 1362, 1350, 1339, 1317, 1268, 1227, 1218, 1190, 1114, 1103, 1087, 1066, 1019, 956, 927, 897, 868, 833, 781, 752, 743, 676, 609, 567, 556

**$^1$H NMR** (400 MHz, DMSO-*d6*) δ 11.51 (s, 1H), 10.74 (s, 1H), 7.76 (d, *J* = 6.1 Hz, 1H), 5.97 (t, *J* = 5.7 Hz, 1H), 3.29 (t, *J* = 5.6 Hz, 2H), 3.24 (s, 3H), 3.13 (q, *J* = 5.6 Hz, 2H).

**$^{19}$F NMR** (57 MHz, Acetone-*d6*) δ -171.50 (d, *J* = 6.1 Hz).

**$^{13}$C NMR** (101 MHz, DMSO-*d6*) δ 158.54, 158.39, 158.28, 150.54, 141.43, 139.17, 126.93, 126.61, 72.09, 58.34, 39.45.

**$^{1}$H NMR for Compound 7** (400 MHz, DMSO-*d6*)**:**

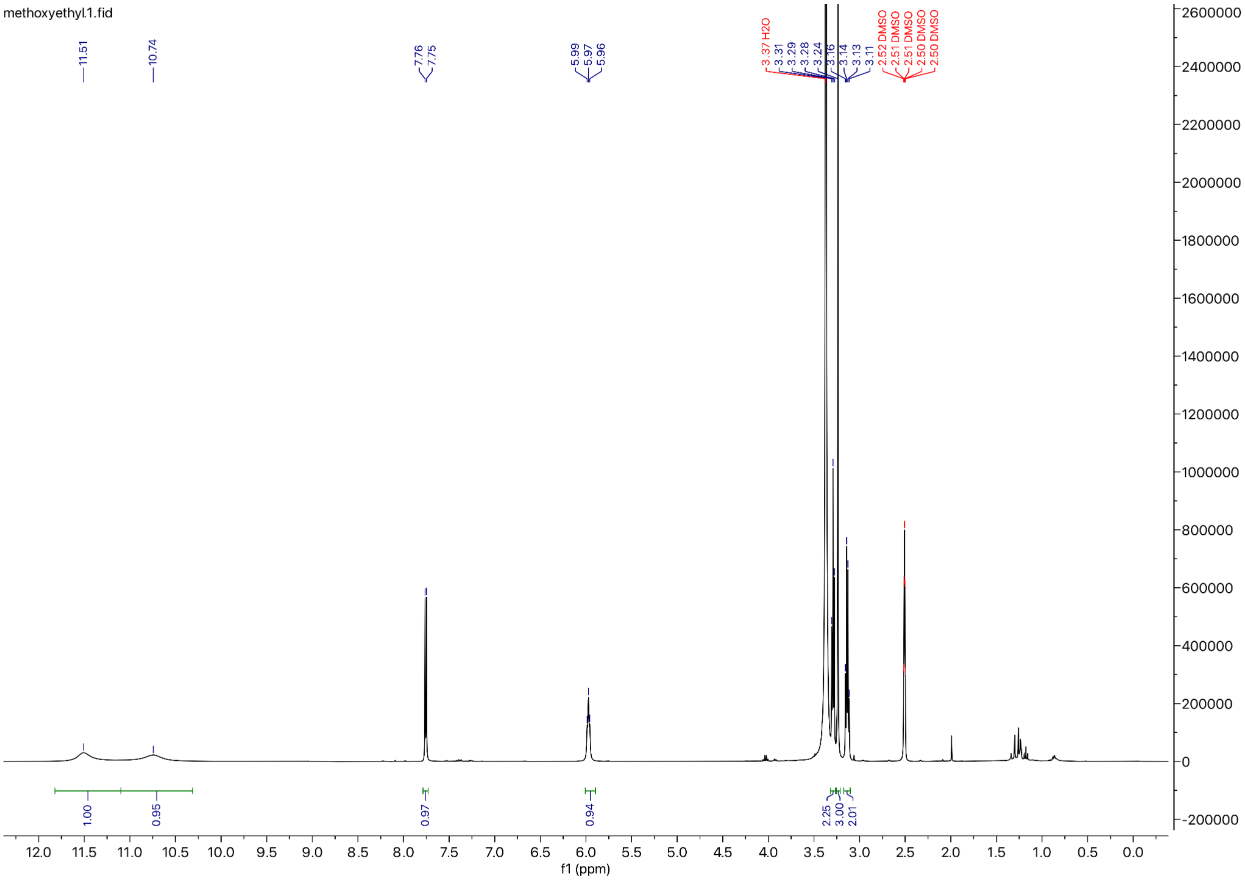

**$^{19}$F NMR for Compound 7** (57 MHz, Acetone-*d*6)**:**

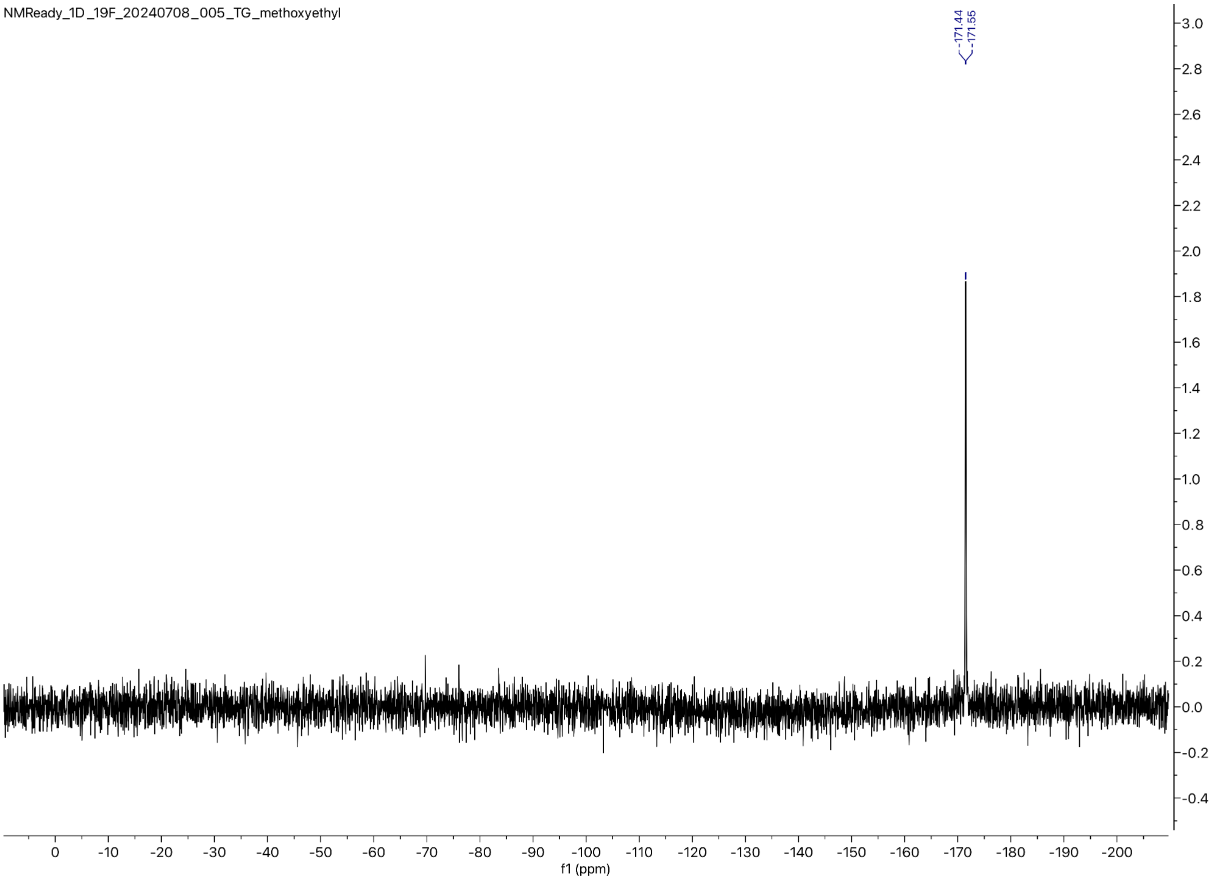

**$^{13}$C NMR for Compound 7** (101 MHz, Acetone-*d6*)**:**

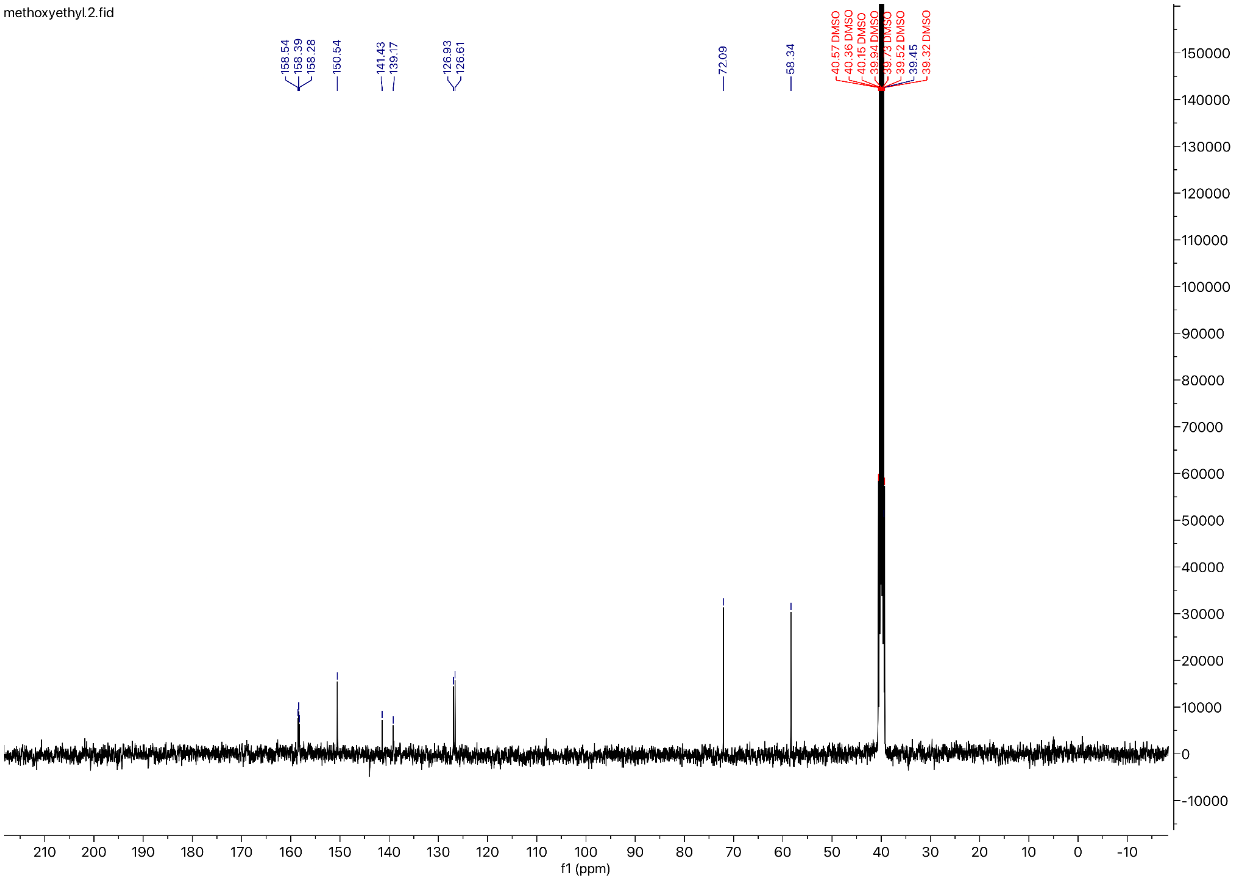

#### *Preparation of the Ethyl glycinate urethane analog of Carmofur*

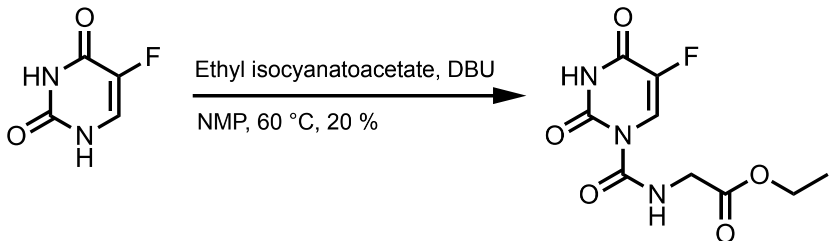


8

Following the general procedure yielded a white solid (20 % yield)

MW: 259.193 g/mol

**Chemicals:**

5-Fluorouracil (AK Scientific, 95 %): used without further purification
N-methylpyrrolidinone (Sigma Aldrich, 99.5 %, SureSeal bottle): used without further purification
1,8-Diazabicyclo[5.4.0]undec-7-ene (AK Scientific, 99 %): used without further purification
Ethyl isocyanatoacetate (Sigma Aldrich, 98 %): used without further purification

**Procedure:**

To a 50 mL oven-dried three-neck flask fitted with a water-jacketed condenser and a Teflon stir bar was added 5-fluorouracil (300.0 mg, 2.31 mmol, 1.0 eq) under nitrogen, and then anhydrous N-methylpyrrolidinone (8.00 mL, 0.29 M) was added via syringe followed by 1,8-diazabicyclo[5.4.0]undec-7-ene (0.051 mL, 0.35 mmol, 0.15 eq) and ethyl isocyanatoacetate (0.388 mL, 3.46 mmol, 1.5 eq). The reaction was heated to 60 ℃ in a silicone oil bath and tracked by aliquot $^{19}$F NMR spectroscopy until reaction completion after 120 minutes. The reaction mixture was then quenched with 1.0 M brine (50 mL), aq. 1.0 M HCl (50 mL), and extracted with ethyl acetate (2x50 mL). The combined organic layers were dried over anhydrous magnesium sulfate, filtered, and concentrated *in vacuo*. The resulting residue was directly loaded onto a silica column (2.3 cm dia., 10 cm stack) and purified by flash chromatography (0 % → 40 % EtOAc in hexanes) to afford the title compound as a white powder (120. mg, 20 % yield).

**Characterization Data for Compound 8:**

**TLC $R_f$**: 0.60 (30 % EtOAc/Hexanes), UV Active

**GC-MS:** Calculated for $[C_9H_{10}FN_3O_5]^+$ $[M]^+$: 259.193; found: 259.208

**FT-IR (ATR, $cm^{-1}$):** 3318, 3074, 2995, 2832, 2358, 2333, 1731, 1715, 1693, 1660, 1510, 1474, 1445, 1397, 1380, 1346, 1264, 1245, 1236, 1195, 1178, 1122, 1065, 1012, 943, 867, 797, 768 727, 657, 612, 578, 563, 553

**$^1$H NMR** (400 MHz, DMSO-*d6*) δ 11.01 (s, 1H), 7.70 (d, *J* = 6.1 Hz, 1H), 6.45 (t, *J* = 5.9 Hz, 1H), 4.05 (q, *J* = 7.1 Hz, 2H), 3.74 (d, *J* = 6.1 Hz, 2H), 1.15 (t, *J* = 7.1 Hz, 3H).

**$^{19}$F NMR** (57 MHz, DMSO-*d6*) δ -171.48 (d, *J* = 7.6 Hz).

**$^{13}$C NMR** (101 MHz, DMSO-*d6*) δ 192.20, 177.29, 171.36, 159.03, 158.26, 142.84, 124.06, 122.80, 60.61, 42.17, 14.51.

**$^1$H NMR for Compound 8** (400 MHz, DMSO-*d6*)**:**

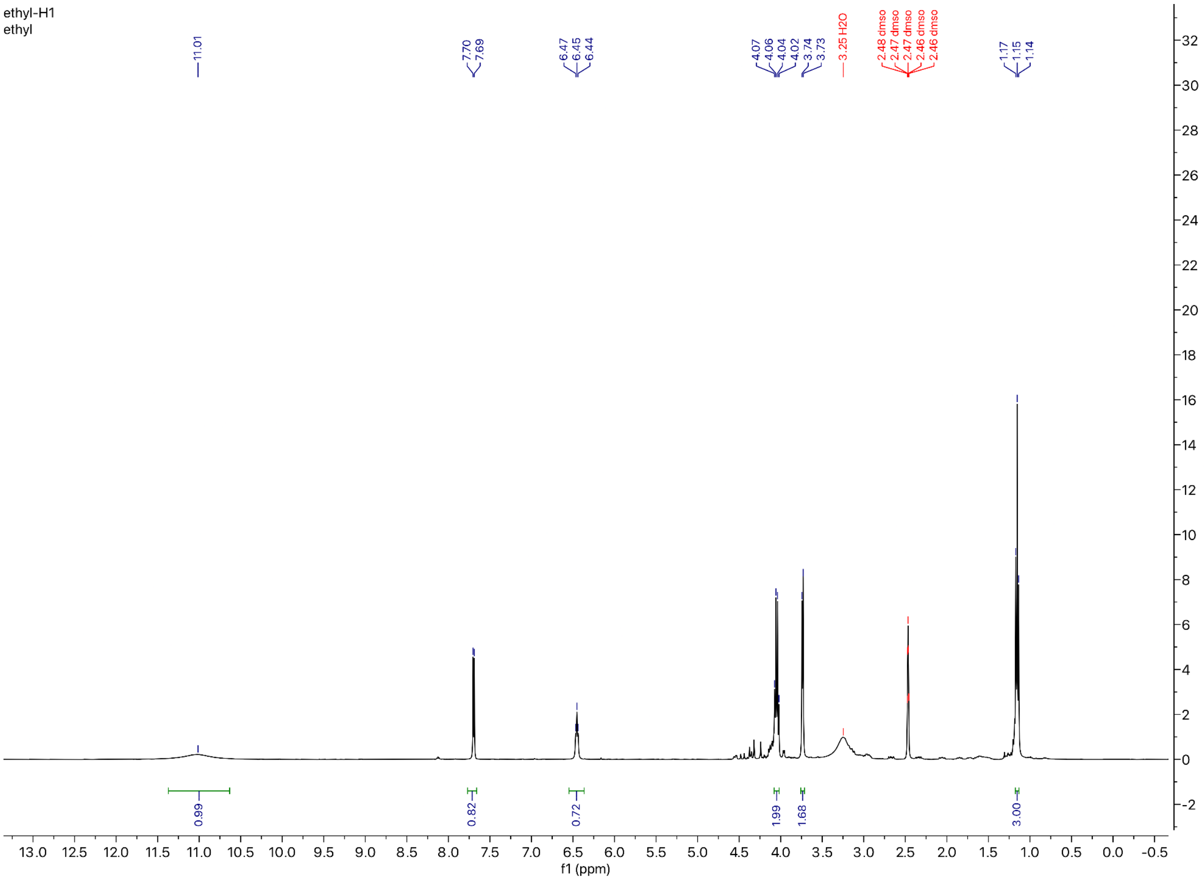

**$^{19}$F NMR for Compound 8** (57 MHz, DMSO-*d*6)**:**

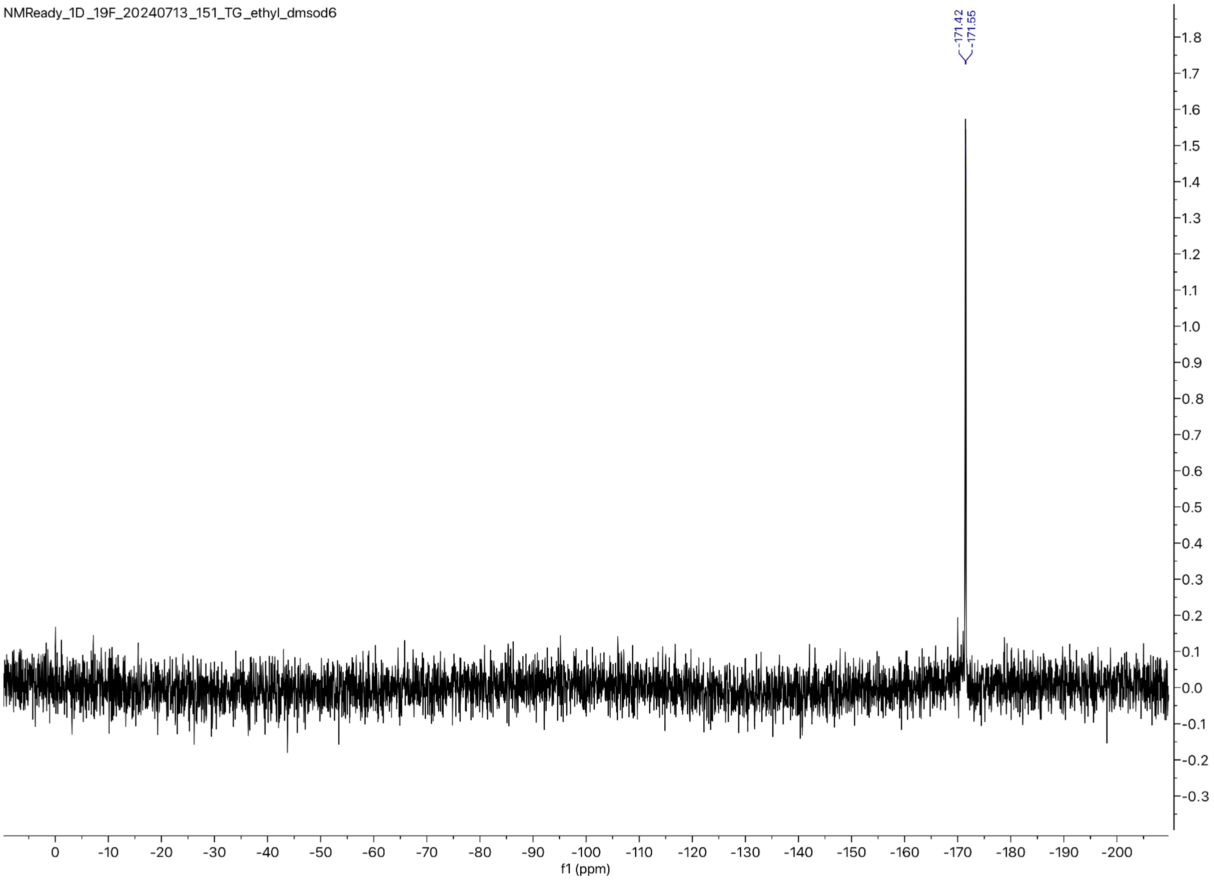

**$^{13}$C NMR for Compound 8** (101 MHz, DMSO-*d6*)**:**

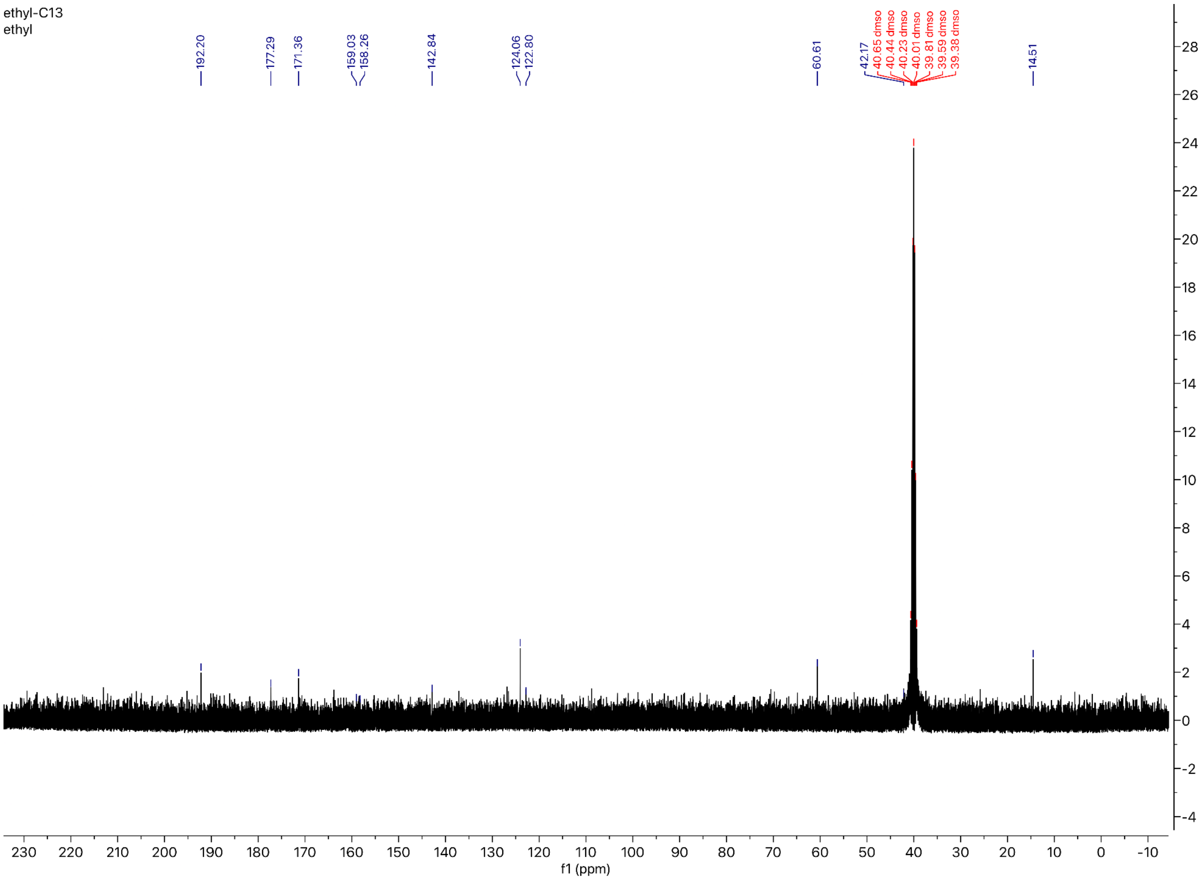

*Preparation of the Hexyl carbamate analog of Carmofur:*

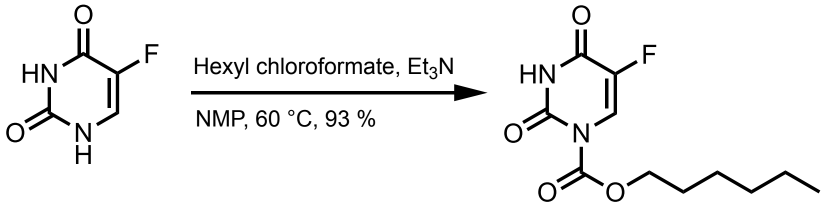


**9**

Following the general procedure yielded a white solid (93 % yield)

MW: 258.249 g/mol

**Chemicals:**
5-Fluorouracil (AK Scientific, 95 %): used without further purification
N-methylpyrrolidinone (Sigma Aldrich, 99.5 %, SureSeal bottle): used without further purification
Triethylamine (Sigma Aldrich, 99.5 %): used without further purification
Hexyl chloroformate (AK Scientific, 95 %): used without further purification

**Procedure:**
To a 50 mL oven-dried three-neck flask fitted with a water-jacketed condenser and a Teflon stir bar was added 5-fluorouracil (630.0 mg, 4.84 mmol, 1.0 eq) under nitrogen, and then anhydrous N-methylpyrrolidinone (20.00 mL, 0.24 M) was added via syringe followed by triethylamine (0.700 mL, 5.03 mmol, 1.0 eq) and hexyl chloroformate (1.193 mL, 7.30 mmol, 1.51 eq). The reaction was heated to 60 °C in a silicone oil bath and tracked by aliquot $^{19}F$ NMR spectroscopy until reaction completion after 180 minutes. The reaction mixture was then quenched with 1.0 M brine (100 mL) and extracted with ethyl acetate (2x50 mL). The combined organic layers were dried over anhydrous magnesium sulfate, filtered, and concentrated *in vacuo*. The resulting residue was directly loaded onto a silica column (2.3 cm dia., 10 cm stack) and purified by flash chromatography (10 % → 30 % EtOAc in hexanes) to afford the title compound as a white powder (1.159 g, 93 % yield).

**Characterization Data for Compound 9:**

**TLC:** $R_f$ = 0.40 (75 % EtOAc/Hexanes), UV Active

**GC-MS:** Calculated for $[C_{11}H_{15}FN_2O_4]^+$ $[M]^+$: 258.249; found: 258.243

**FT-IR (ATR, $cm^{-1}$):** 3201, 3097, 2954, 2933, 2872, 2849, 1739, 1722, 1683, 1465, 1436, 1388, 1341, 1286, 1228, 1195, 1122, 1099, 1042, 1011, 989, 916, 899, 842, 799, 768, 735, 696, 607, 580, 573, 563, 553

**$^1$H NMR** (400 MHz, Acetone-*d6*) δ 10.48 (s, 1H), 8.03 (d, *J* = 7.1 Hz, 1H), 4.25 (t, *J* = 6.6 Hz, 2H), 1.63 (p, *J* = 7.8 Hz, 2H), 1.33 (p, *J* = 7.5 Hz, 2H), 1.24 – 1.18 (m, 4H), 0.76 (t, *J* = 6.9 Hz, 3H).

**$^{19}$F NMR** (57 MHz, Chloroform-*d*) δ -162.06 (d, *J* = 6.1 Hz).

**$^{13}$C NMR** (101 MHz, Acetone-*d6*) δ 150.27, 149.99, 145.63, 124.13, 123.76, 68.99, 31.13, 28.11, 25.11, 22.24, 13.32.

**$^1$H NMR for Compound 9** (400 MHz, Acetone-*d6*):

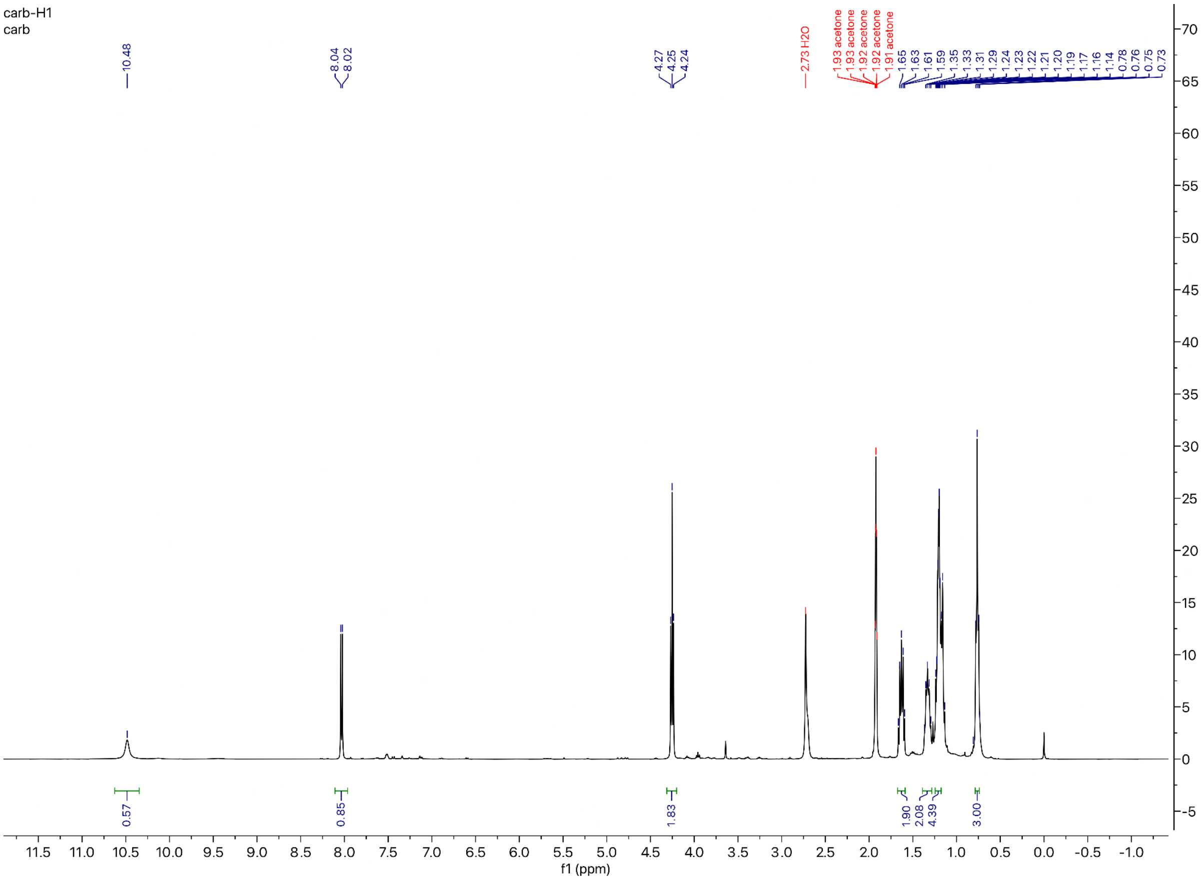

**$^{19}$F NMR for Compound 9** (57 MHz, Chloroform-*d*):

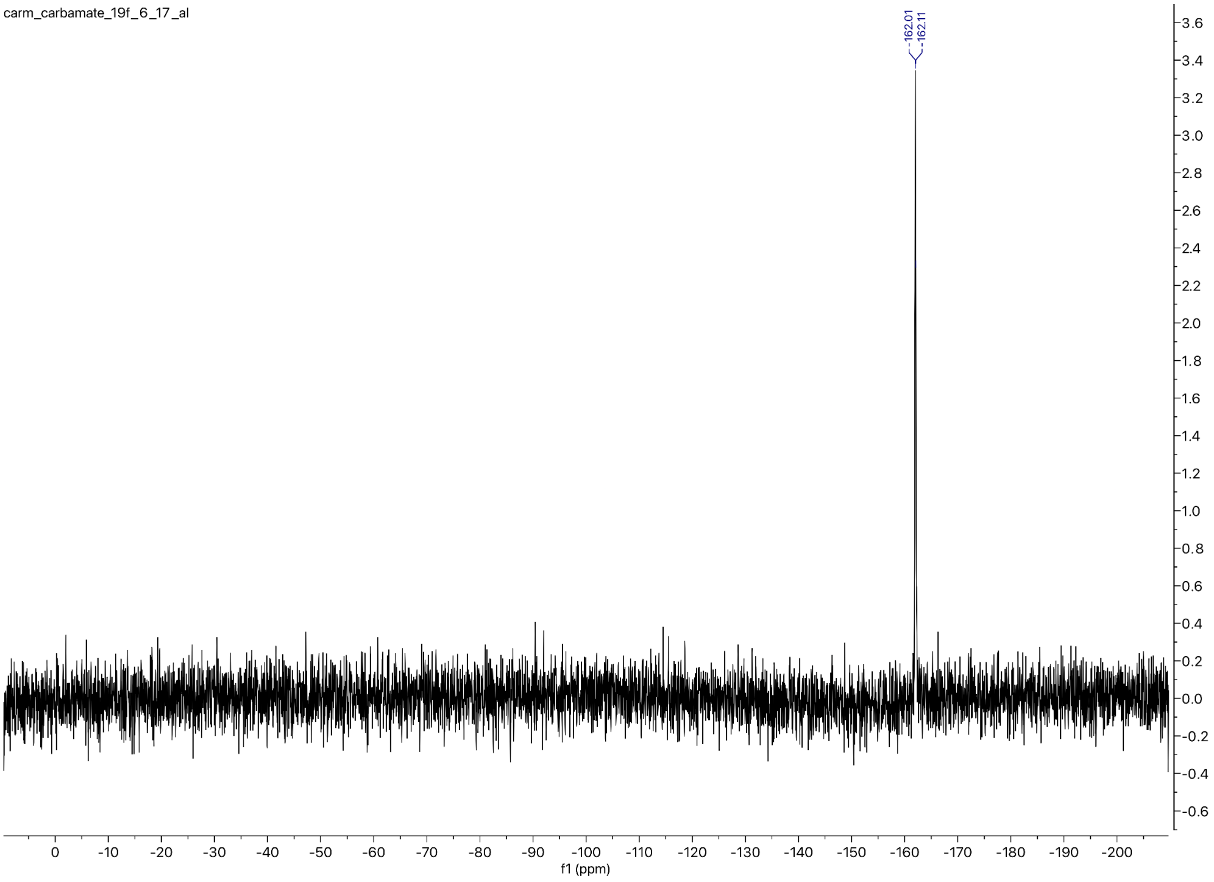

**$^{13}$C NMR for Compound 9** (101 MHz, Acetone-*d6*):

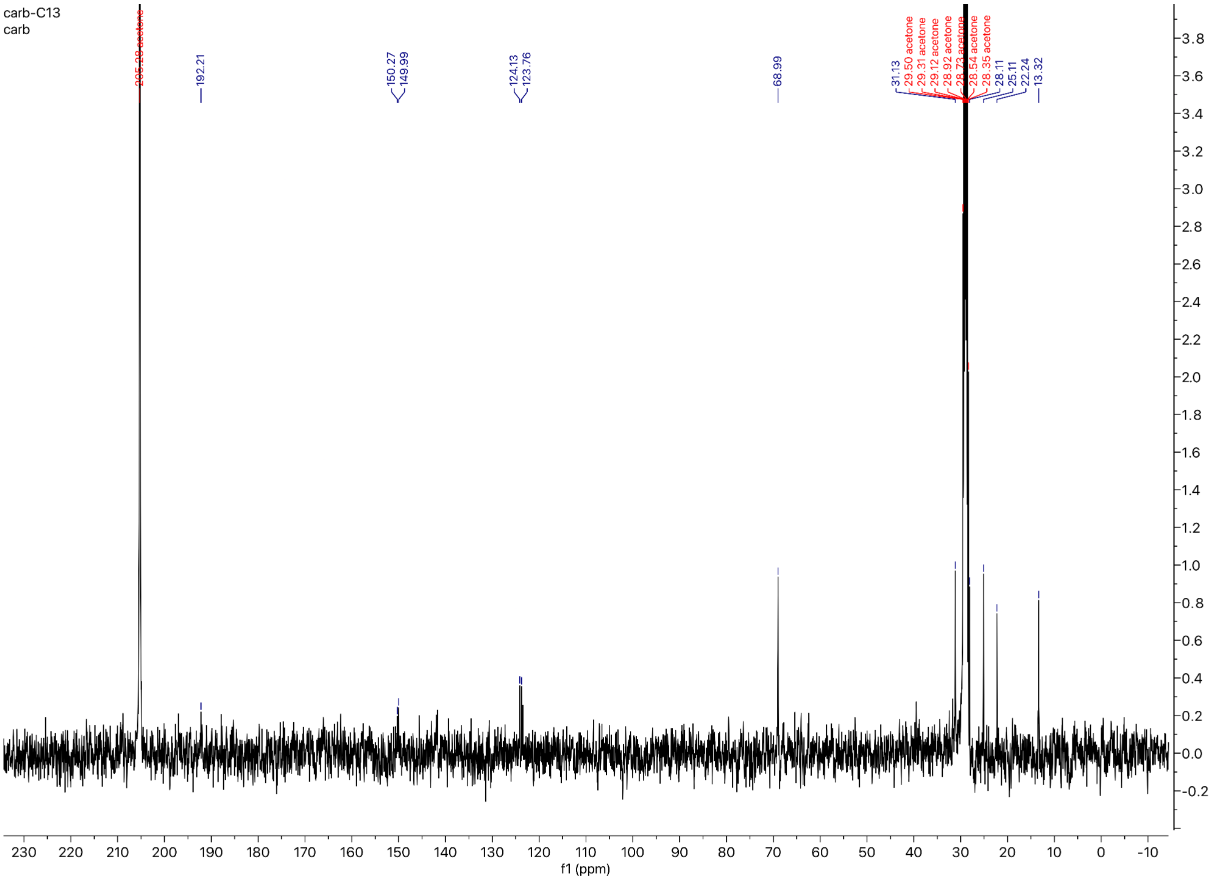

#### *Preparation of the Heptyl amide analog of Carmofur:*

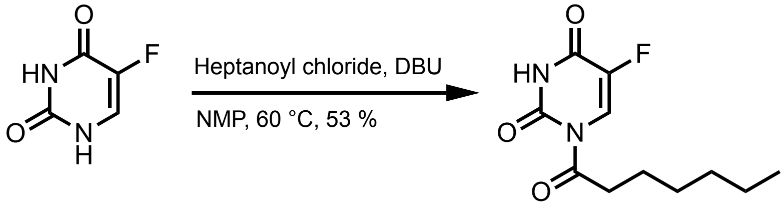


**10**

Following the general procedure yielded a white solid (53 % yield)

MW: 242.250 g/mol

**Chemicals:**
5-Fluorouracil (AK Scientific, 95 %): used without further purification
N-methylpyrrolidinone (Sigma Aldrich, 99.5 %, SureSeal bottle): used without further purification
1,8-Diazabicyclo[5.4.0]undec-7-ene (AK Scientific, 99 %): used without further purification
Heptanoyl chloride (AK Scientific, 95 %): used without further purification

**Procedure:**
To a 50 mL oven-dried three-neck flask fitted with a water-jacketed condenser and a Teflon stir bar was added 5-fluorouracil (200.0 mg, 1.54 mmol, 1.0 eq) under nitrogen, and then anhydrous N-methylpyrrolidinone (12.00 mL, 0.13 M) was added via syringe followed by 1,8-diazabicyclo[5.4.0]undec-7-ene (0.034 mL, 0.23 mmol, 0.15 eq) and heptanoyl chloride (0.476 mL, 3.08 mmol, 2.0 eq). The reaction was heated to 60 °C in a silicone oil bath and tracked by aliquot $^{19}$F NMR spectroscopy until reaction completion after 150 minutes. The reaction mixture was then quenched with 1.0 M brine (50 mL), aq. 1.0 M HCl (50 mL), and extracted with ethyl acetate (2x50 mL). The combined organic layers were dried over anhydrous magnesium sulfate, filtered, and concentrated *in vacuo*. The resulting residue was directly loaded onto a silica column (2.3 cm dia., 10 cm stack) and purified by flash chromatography (0 % → 30 % EtOAc in hexanes) to afford the title compound as a white powder (199 mg, 53 % yield).

**Characterization Data for Compound 10:**

**TLC $R_f$:** 0.80 (60 % EtOAc/Hexanes), UV Active

**GC-MS:** Calculated for $[C_{11}H_{15}FN_2O_3]^+$ $[M]^+$: 242.250; found: 243.250

**FT-IR (ATR, $cm^{-1}$):** 3188, 3108, 3065, 2955, 2925, 2848, 1727, 1682, 1559, 1454, 1431, 1395, 1374, 1339, 1318, 1295, 1281, 1259, 1226, 1128, 1098, 1034, 1018, 928, 851, 796, 775, 755, 742, 721, 643, 604, 564, 552

**$^1$H NMR** (400 MHz, Chloroform-*d*) δ 8.56 (s, 1H), 8.22 (d, *J* = 6.7 Hz, 1H), 3.05 (t, *J* = 7.3 Hz, 2H), 1.65 (p, *J* = 7.1 Hz, 2H), 1.35 – 1.22 (m, 6H), 0.83 (t, *J* = 6.9 Hz, 3H).

**$^{19}$F NMR** (57 MHz, Chloroform-*d*) δ -161.82 (d, *J* = 6.1 Hz).

**$^{13}$C NMR** (101 MHz, Chloroform-*d*) δ 219.78, 171.99, 147.63, 121.90, 121.54, 39.05, 31.47, 28.59, 24.37, 22.47, 14.00.

**[1]H NMR for Compound 10** (400 MHz, Chloroform-*d*):

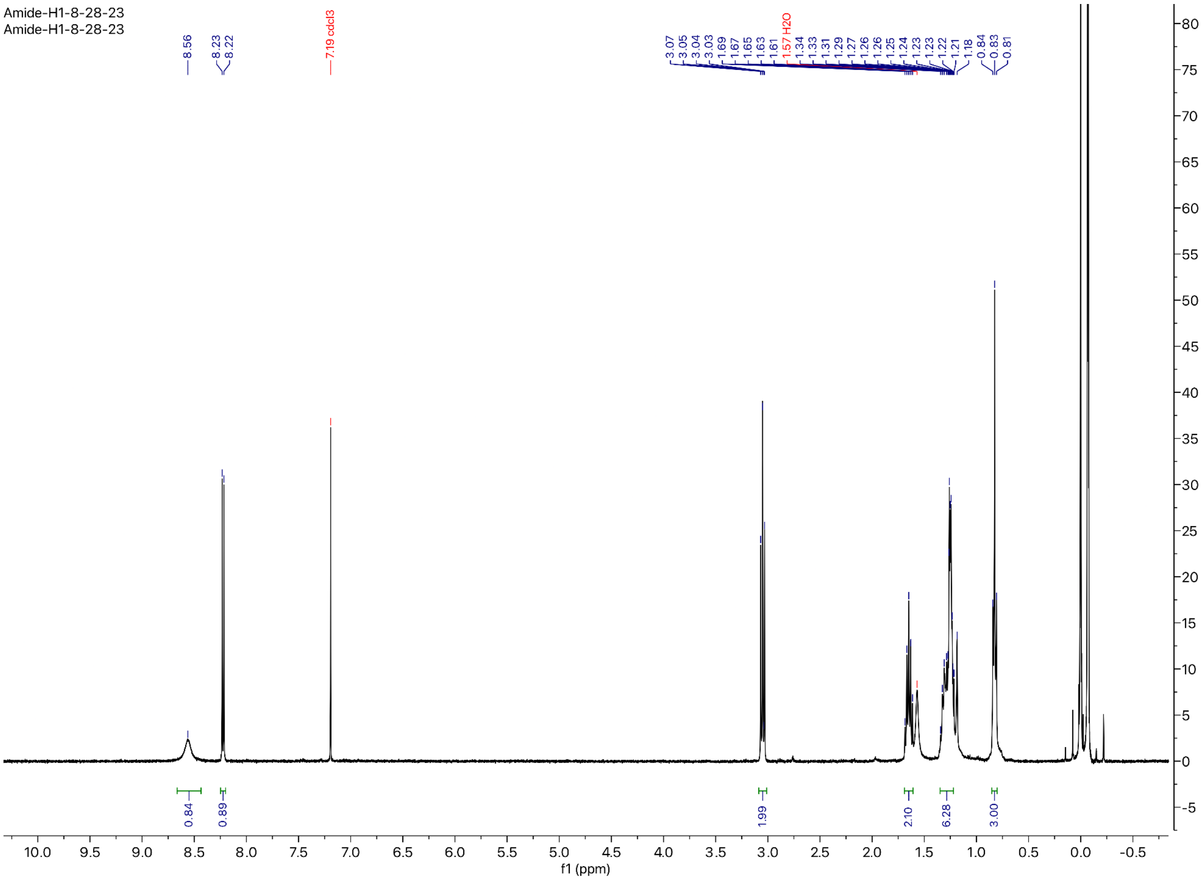

**$^{19}$F NMR for Compound 10** (57 MHz, Chloroform-*d*):

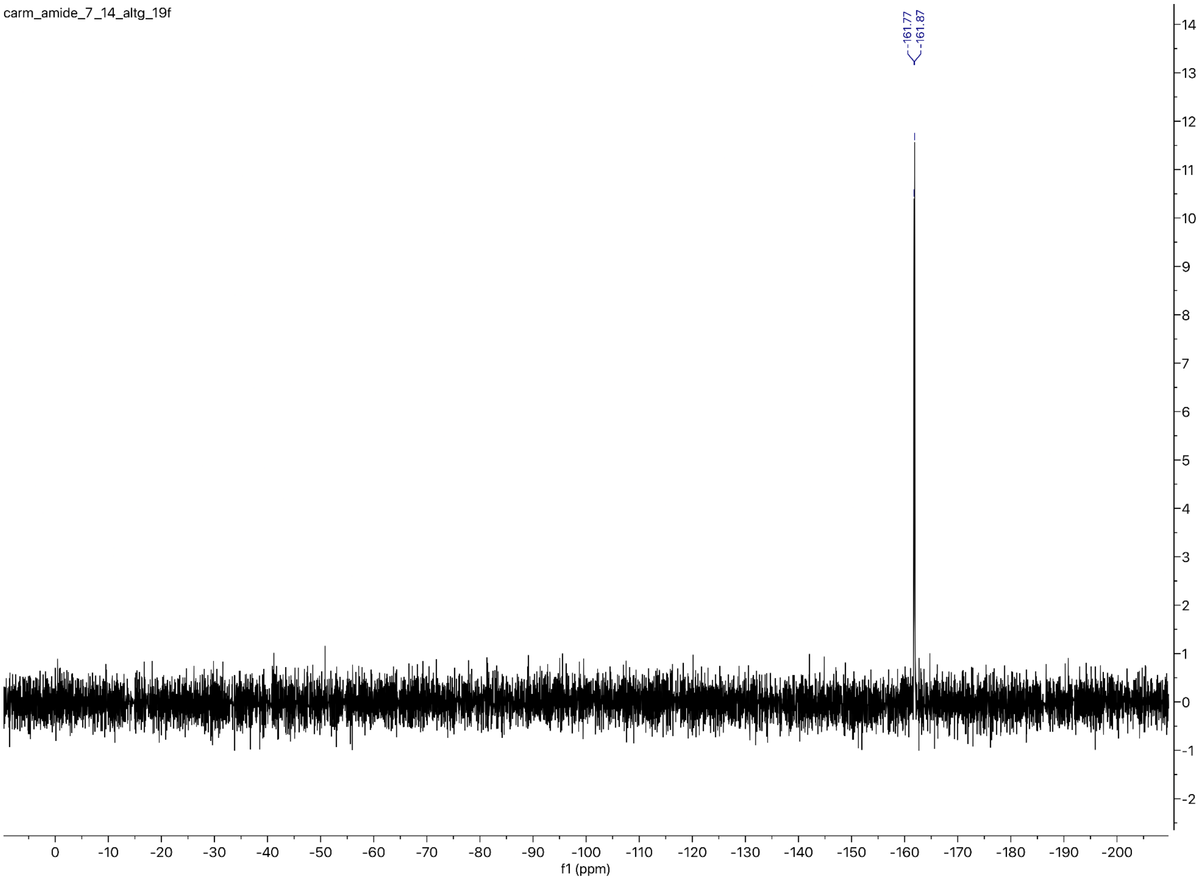

**$^{13}$C NMR for Compound 10** (101 MHz, Chloroform-*d*):

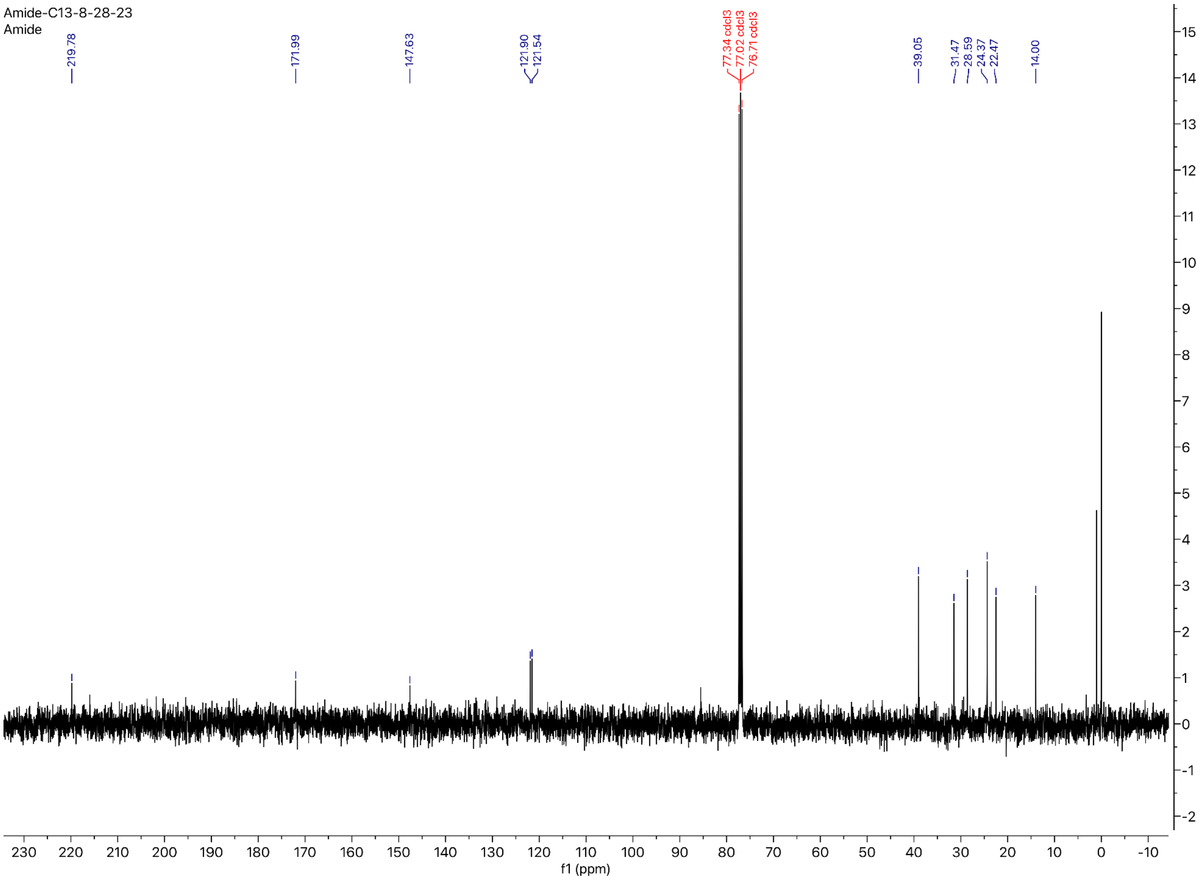

### References:


1. Willoughby PH, Jansma MJ, Hoye TR. A guide to small-molecule structure assignment through computation of (1 H and 13 C) NMR chemical shifts. Nat Protoc. 2014;9(3):643–60. https://doi.org/10.1038/nprot.2014.042.
2. Mohamadi F, Richards NGJ, Guida WC, Liskamp R, Lipton M, Caufield C, et al. MacroModel-An Integrated Software System for Modeling Organic and Bioorganic Molecules Using Molecular Mechanics. J Comput Chem. 1990;11(4):440–67. https://doi.org/10.1002/jcc.540110405.
3. Chen IJ, Foloppe N. Tackling the conformational sampling of larger flexible compounds and macrocycles in pharmacology and drug discovery. Bioorg Med Chem. 2013;21(24):7898–920. https://doi.org/10.1016/j.bmc.2013.10.003.
4. Watts KS, Dalal P, Tebben AJ, Cheney DL, Shelley JC. Macrocycle conformational sampling with macromodel. J Chem Inf Model. 2014;54(10):2680–96. https://doi.org/10.1021/ci5001696.
5. Bochevarov AD, Harder E, Hughes TF, Greenwood JR, Braden DA, Philipp DM, et al. Jaguar: A high-performance quantum chemistry software program with strengths in life and materials sciences. Int J Quantum Chem. 2013;113(18):2110–42. https://doi.org/10.1002/qua.24481.
6. Wang Y-C, Yang W-H, Yang C-S, Hou M-H, Tsai C-L, Chou Y-Z, et al. Structural basis of SARS-CoV-2 main protease inhibition by a broad-spectrum anti-coronaviral drug. Am J Cancer Res. 2020;10(8):2535–45.
7. Jin Z, Zhao Y, Sun Y, Zhang B, Wang H, Wu Y, et al. Structural basis for the inhibition of SARS-CoV-2 main protease by antineoplastic drug carmofur. Nat Struct Mol Biol. 2020;27(6):529–32. https://doi.org/10.1038/s41594-020-0440-6.
8. Dementiev A, Joachimiak A, Nguyen H, Gorelik A, Illes K, Shabani S, et al. Molecular Mechanism of Inhibition of Acid Ceramidase by Carmofur. J Med Chem. 2019;62(2):987–92. https://doi.org/10.1021/acs.jmedchem.8b01723.
9. Madhavi Sastry G, Adzhigirey M, Day T, Annabhimoju R, Sherman W. Protein and ligand preparation: Parameters, protocols, and influence on virtual screening enrichments. J Comput Aided Mol Des. 2013;27(3):221–34. https://doi.org/10.1007/s10822-013-9644-8.
10. Friesner RA, Banks JL, Murphy RB, Halgren TA, Klicic JJ, Mainz DT, et al. Glide: A New Approach for Rapid, Accurate Docking and Scoring. 1. Method and Assessment of Docking Accuracy. J Med Chem. 2004;47(7):1739–49. https://doi.org/10.1021/jm0306430.
11. Halgren TA, Murphy RB, Friesner RA, Beard HS, Frye LL, Pollard WT, et al. Glide: A New Approach for Rapid, Accurate Docking and Scoring. 2. Enrichment Factors in Database Screening. J Med Chem. 2004;47(7):1750–9. https://doi.org/10.1021/jm030644s.
12. Friesner RA, Murphy RB, Repasky MP, Frye LL, Greenwood JR, Halgren TA, et al. Extra precision glide: Docking and scoring incorporating a model of hydrophobic enclosure for

protein-ligand complexes. J Med Chem. 2006;49(21):6177–96.
https://doi.org/10.1021/jm051256o.
13. Yang Y, Yao K, Repasky MP, Leswing K, Abel R, Shoichet BK, et al. Efficient Exploration of Chemical Space with Docking and Deep Learning. J Chem Theory Comput. 2021;17(11):7106–19. https://doi.org/10.1021/acs.jctc.1c00810.
14. Wang X, Vu J, Luk C, Njoo, E. Benchtop 19F nuclear magnetic resonance spectroscopy enabled kinetic studies and optimization of the synthesis of carmofur. Can J Chem. 2023;101(8):518–24. https://doi.org/10.1139/cjc-2022-0266.
15. Pazzi J, Subramaniam AB. Nanoscale Curvature Promotes High Yield Spontaneous Formation of Cell-Mimetic Giant Vesicles on Nanocellulose Paper. ACS Appl Mater Interfaces. 2020;12(50):56549–61. https://pubs.acs.org/doi/10.1021/acsami.0c14485.
